# Expanding the genetic code with diverse backbone structures across diverse sequence contexts

**DOI:** 10.64898/2026.04.16.718949

**Authors:** Carlos Piedrafita, Alexandre Dickson, Daniel Richter, Caroline Weber, Thomas S. Elliott, Ziqi Liu, Fan Zhang, Yichen Li, Daniel L. Dunkelmann, Tomos Morgan, Kim C. Liu, Jason W. Chin

## Abstract

Expanding the genetic code to enable the selective and specific incorporation of non-canonical monomers (ncMs), beyond α-L amino acids with variant sidechains, is a key outstanding challenge. Here we discover orthogonal aminoacyl-tRNA synthetases that selectively and specifically acylate their cognate orthogonal tRNA in vivo with eleven new ncMs spanning five different chemical classes: α,α-disubstituted-amino acids, malonic acids, carboxylic acids, β^2^-amino acids and N-cyclic amino acids. We demonstrate that co-translational incorporation of α,α-disubstituted-amino acids, β^2^-amino acids, β^3^-amino acids and N-cyclic amino acids is strongly dependent on the codons either side of the codon used to direct ncM incorporation, with several ncMs incorporated at less than 1% of sequence contexts. We evolve orthogonal tRNAs that enable the incorporation of previously unincorporated ncMs, enable the incorporation of ncMs at >95% of sequence contexts and, increase the incorporation efficiency at challenging sequence contexts up to 40-fold. We demonstrate the encoded cellular synthesis of proteins and macrocycles containing ncMs and, explicitly demonstrate that our evolved tRNAs provide direct access to a wider range of genetically encoded macrocyclic sequences containing ncMs. Our results provide a foundation for composing, discovering and manufacturing proteins and peptides with functions augmented by ncMs.

## Introduction

The genetic code of living organisms has been expanded to direct the incorporation of diverse α-L-amino acids with variant side chains (non-canonical amino acids, ncAAs)^1,2^, and their close analogs, into proteins, peptides, macrocycles and genetically encoded polymers^3–6^. In contrast, the in vivo, genetically directed, site-specific incorporation of non-canonical monomers (ncMs), with diverse backbone structures, has proved exceptionally challenging. While substantial progress has been made on encoding ncMs in in vitro reactions^7–14^, these approaches cannot generally be extended to live cells.

Recent work has demonstrated the in vivo acylation of several ncMs (β^3^-amino acids, β^3^-hydroxy acids, β^2^-amino acids, β^2^-hydroxy acids, α,α-disubstituted amino acids and malonic acids) on to tRNAs by orthogonal synthetases^15–18^. The development of tRNA display (**Supplementary Fig. 1**)^15^ and other methods for the direct selection of orthogonal synthetases that specifically and selectively acylate their cognate orthogonal tRNA with ncMs has been crucial for expanding the scope of monomers that can be acylated onto tRNAs, whether-or-not these monomers are substrates for translation^15,19^. It has been proposed that the direct discovery of synthetases that acylate orthogonal tRNAs with ncMs will unlock the ability to select other translational components that enhance or enable the co-translational incorporation of the ncMs^15^, but this has not been demonstrated.

A subset of the ncMs (β^3^-amino acids, β^2^-hydroxy acids and α,α-disubstituted amino acids) that can be acylated onto a tRNA have been co-translationally incorporated in at least one site in a protein^15,16^; however, it is unclear whether these ncMs can be incorporated at other sequence contexts. Other ncMs that can be acylated onto an orthogonal tRNA have not been incorporated into proteins in vivo. We hypothesized that the ability to directly discover orthogonal synthetases that acylate their cognate tRNAs with ncMs would enable the co-translational selection of other translational components that allow the incorporation of the ncM into proteins, and the incorporation of ncMs at diverse sequence contexts. These advances would enable the power of genetically encoded cellular synthesis to be generally leveraged for composing proteins, peptides and macrocycles containing ncMs.

Here we use tRNA display to discover *Methanosarcina mazei* pyrrolysyl synthetase (*Mm*PylRS, henceforth referred to as PylRS) variants that specifically and efficiently acylate their cognate tRNA (*Mm*tRNA^Pyl^_CUA_, henceforth referred to as tRNA^Pyl^) in vivo with ncMs from five chemical classes: β^2^-amino acids, non-canonical N-cyclic amino acids, malonic acids, carboxylic acids, as well as α,α-disubstituted amino acids. Using our newly discovered PylRS variants, we demonstrate, to our knowledge, the first site-specific incorporation of β^2^-amino acids and non-canonical N-cyclic amino acids into a protein by cellular translation. We then systematically investigate in vivo translation of ncMs with non-canonical backbone structures, namely β^3^-amino acids, β^2^-amino acids, N-cyclic amino acids, and α,α-disubstituted amino acids. We observe that the in vivo incorporation of these ncMs by the *E. coli* translational machinery is inefficient and strongly dependent on the codons immediately flanking the target codon. We develop a pipeline for directed evolution of tRNAs in vivo and discover tRNA variants which enable the incorporation of several monomers that could not be incorporated previously, and mediate up to 40-fold improved incorporation efficiencies for the four monomer classes. Moreover, we demonstrate that the optimised tRNA variants allow incorporation of ncMs at a wider range of sequence contexts. While wildtype tRNAs incorporated some ncMs at <1% sequence contexts, the evolved tRNAs enabled ncM incorporation at up to >95% sequence contexts. These advances directly enabled the genetically encoded cellular synthesis of macrocycles containing β^3^-amino acids, β^2^-amino acids, N-cyclic amino acids, and α,α-disubstituted amino acids and the synthesis of a wider range of ncM-containing macrocycles.

## Results

### Expanding the chemical diversity of noncanonical monomers by tRNA display

To further explore the scope of tRNA display for discovering orthogonal synthetases that direct the acylation of their cognate orthogonal tRNAs with diverse non-canonical monomers, we performed selections with a library of *Mm*PylRS variants and ncMs from three new monomer classes (**Fig. 1a, Supplementary Fig. 2**): β^2^-amino acids (**3**), N-cyclic amino acids (**4**), and malonic acids (**6** and **7**). Additionally, we performed tRNA display with an α,α-disubstituted amino acid (**8**). We performed two rounds of enrichment, in the presence and absence of each ncM, to select for synthetase variants that acylated tRNA^Pyl^ (**Supplementary Fig. 1**). By analysing next-generation sequencing (NGS) of the selected samples we identified synthetase variants (‘hits’) that were selectively enriched in the presence of the ncMs **3, 4** and **6-8** (**Supplementary Figs. 3-7**).

**Figure 1.**
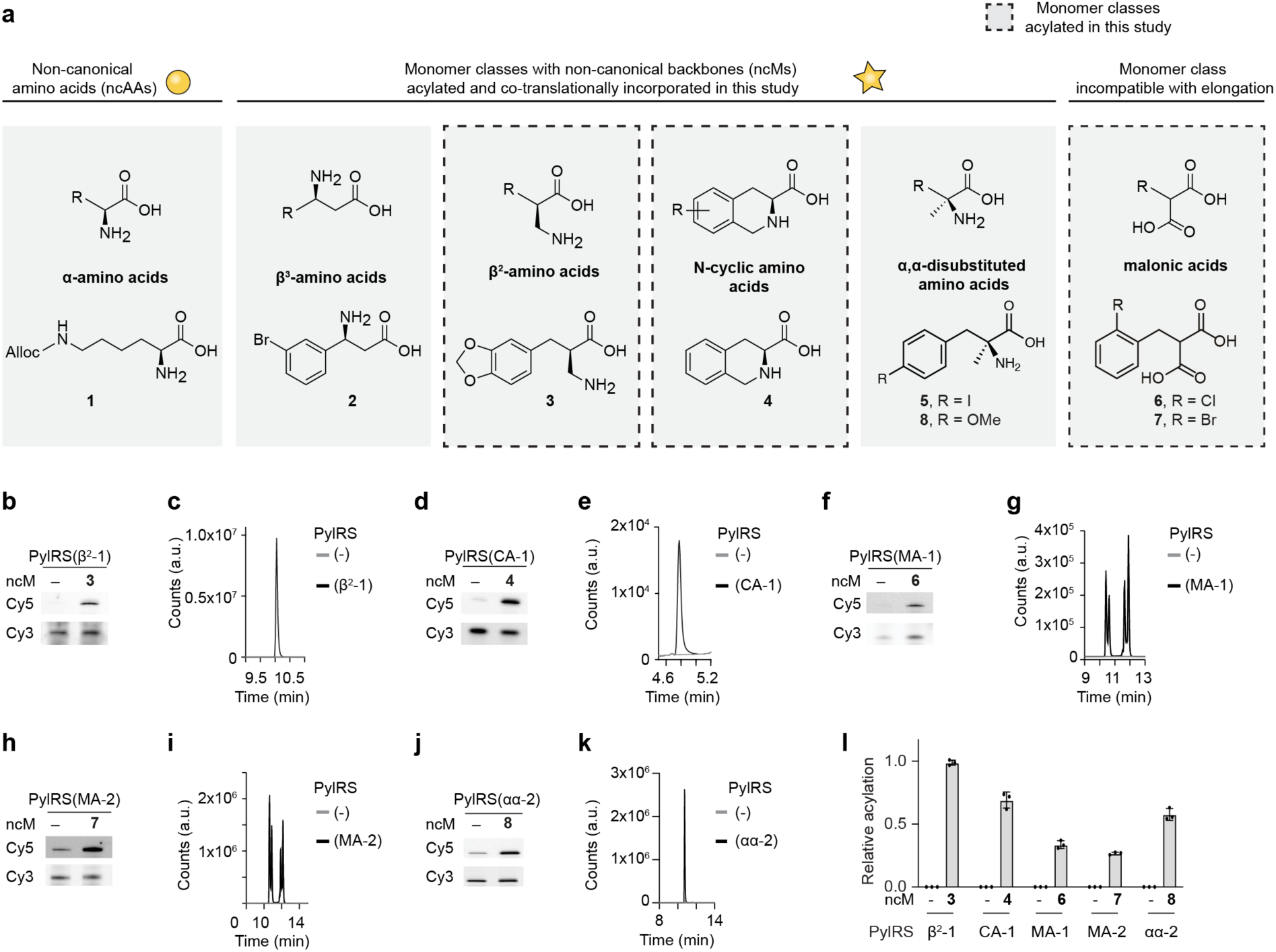
Acylation of diverse non-canonical monomers enabled by tRNA display. **a**, Structures of Nε-Alloc-lysine (**1**) and structures of ncMs used in this study, the β^3^-amino acid **2**, β^2^-amino acid **3**, the N-cyclic amino acid **4**, α,α-disubstituted amino acids **5** and **8**, and malonic acids **6** and **7**. **b**, ftREX detection of tRNA^Pyl^ acylation by PylRS(β^2^-1) in the absence and presence of ncM **3**. ftREX was performed in biological duplicates. **c**, Single ion monitoring LC-MS traces for the 6-aminoquinolyl-*N*-hydroxysuccinimidyl carbamate (AQC) adduct of compound **3** (SIM AQC-**3**, [M+H]^+^: *m/z* = 394.0) released from tRNA^Pyl^ after isolation and pulldown. Expression was conducted in the presence (black trace) or absence (grey trace) of PylRS(β^2^-1). **d**,**f**,**h**,**j**, As in **b**, but with the indicated PylRS variants and ncMs **4** (**d**), **6** (**f**), **7** (**h**), **8 (j**)**. e**,**g**,**i,k**, As in **c**, but with the indicated PylRS variants and for the AQC adduct of **4** (**e**, SIM AQC-**4**, [M+H]^+^: *m/z* = 348.0), for the adenosine adduct of **6** (**g**, EIC ad-**6**, [M+H]^+^: *m/z* = 478.0-478.2), for the adenosine adduct of **7** (**i**, EIC ad-**7**, [M+H]^+^: *m/z* = 522.0-522.2), and for the AQC adduct of **8** (**k**, EIC AQC-**8**, [M+H]^+^: *m/z* = 380.0-380.2). Experiments were performed in duplicates. **l**, tRNA^Pyl^ acylation relative to wt PylRS with the benchmark substrate **1** quantified by tREX. Acylation was measured for ncMs **3, 4, 6**-**8** and the respective PylRS variants. tRNA^Pyl^ was extracted from cells containing the specified PylRS variant grown in the presence or absence of monomer (2 mM for **1** and 4 mM for all other ncMs). All experiments were performed in triplicates. Bars indicate the mean, individual data points are shown as black dots and error bars indicate the standard deviation.

We evaluated the ncM-dependent acylation of tRNA^Pyl^ by PylRS hits by performing fluoro-tREX (ftREX)^15^ with cells grown in the presence and absence of the respective ncM (**Supplementary Figs. 3-8**). We identified at least one PylRS variant that led to a strong monomer-dependent increase in tRNA^Pyl^ acylation for ncMs **3** (PylRS(β^2^-1)), **4** (PylRS(CA-1)), **6** (PylRS(MA-1)), **7** (PylRS(MA-2)), and **8** (PylRS(αα-2)) (**Fig. 1b,d,f,h,j**).

We confirmed the identity and chirality of the acylated monomers by selective pulldown and mass spectrometry^15,18^ of the acylated tRNA^Pyl^ (**Fig. 1c,e,g,i,k, Supplementary Figs. 8-13**). We further observed that other PylRS hits (PylRS(AC-1) and PylRS(AC-2)) identified from the selections performed with malonic acids **6** and **7** did not acylate tRNA^Pyl^ with their respective ncMs; instead, they specifically acylated tRNA^Pyl^ with carboxylic acids **S1** and **S2**, which may be generated through the in vivo decarboxylation of a fraction of **6** and **7** (**Supplementary Figs. 11-12**). Additional mass spectrometry-based experiments further confirmed that the selected synthetases lead to the acylation of tRNA^Pyl^ with their cognate ncM and only minimally with canonical amino acids in vivo in presence of their cognate ncM (**Supplementary Figs. 14-16**).

Overall, these experiments demonstrated that the selected synthetases, identified by tRNA display, specifically acylate their cognate tRNA in the presence of their respective monomers and lead to minimal acylation of their tRNA with canonical amino acids, even in the absence of competing ncMs.

Next, we set out to quantify intracellular acylation levels of tRNA^Pyl^ achieved by ncM-specific synthetase variants for monomers **2-8**, using tREX. Unlike ftREX – which we used as a screen to discover synthetases that exhibit strong ncM-dependent acylation – tREX produces gel bands corresponding to the acylated and non-acylated form of a tRNA and thereby allows the fraction of acylated tRNA in cells to be directly quantified^20^. PylRS acylated 47.4 ± 1.2% of tRNA^Pyl^ when cells were provided with 2 mM of **1**, a benchmark substrate for this system. We report the acylation of tRNA^Pyl^ with ncMs **2-8** as a percentage of the acylation achieved with this benchmark (**Fig. 1l**, **Supplementary Figs. 17-18**).

For ncMs **3, 4** and **8** – the β^2^-amino acid, N-cyclic amino acid, and α,a-disubstituted amino acid – we observed comparable levels of acylation to α-amino acid **1** with their respective synthetase variants (102.1 ± 2.1% for **3**, 71.3 ± 6.6% for **4**, 59.5 ± 4.9% for **8**). For PylRS(MA-1) with **6** and PylRS(MA-2) with **7** we observed lower levels of acylation; 33.7 ± 3.1% and 26.7 ± 1.0%, respectively. For all PylRS variants acylation was strongly ncM-dependent (**Fig. 1l, Supplementary Fig. 18**), consistent with our observations made with ftREX and our LC-MS based pulldown assay (**Supplementary Figs. 9-16**).

We further tested whether PylRS(β^2^-1) and PylRS(CA-1) could acylate tRNA^Pyl^ with additional β^2^-amino acids and, non-canonical N-cyclic amino acids, respectively (**Supplementary Figs. 19-20)**. ftREX and LC-MS analysis demonstrated that PylRS(β^2^-1) charges the additional monomers **S3-S5** and PylRS(CA-1) charges ncM **S6** onto tRNA^Pyl^ (**Supplementary Figs. 19-20).**

Overall, these data substantially increase the range of monomers that can be specifically acylated onto orthogonal tRNAs in vivo and enable the selective in vivo acylation of orthogonal tRNAs with β^2^-amino acids, non-canonical N-cyclic amino acids and malonic acids.

### Incorporation of β^2^- and N-cyclic amino acids into proteins in vivo

We tested incorporation of ncMs **3**, **4,** and **8**, in response to a TAG codon at position 150 in sfGFP through expression of a sfGFP(150TAG)His_6_ reporter, with tRNA^Pyl^ and PylRS variants in the absence and presence of the corresponding ncMs (**Fig. 2a**). For the β^2^-amino acid **3**, we observed a low level of ncM-dependent increase in GFP fluorescence in the presence of PylRS(β^2^-1). For the N-cyclic amino acid **4** and the α,α-disubstituted amino acid **8,** we observed a strong ncM-dependent increase in GFP fluorescence in the presence of PylRS(CA-1) or PylRS(αα-2), respectively. We confirmed incorporation of ncMs **3**, **4**, and **8** in response to the amber codon by ESI-MS and MS/MS of purified sfGFP(150-**3**)His_6_, sfGFP(150-**4**)His_6_, sfGFP(150-**8**)His_6_, (**Fig. 2b-d, Supplementary Fig. 21**). We further tested incorporation of **6**, **7**, and **S3**-**S6** and observed incorporation of **S5** and **S6** with PylRS(β^2^-1) and PylRS(CA-1) respectively (**Supplementary Fig. 22**).

**Figure 2.**
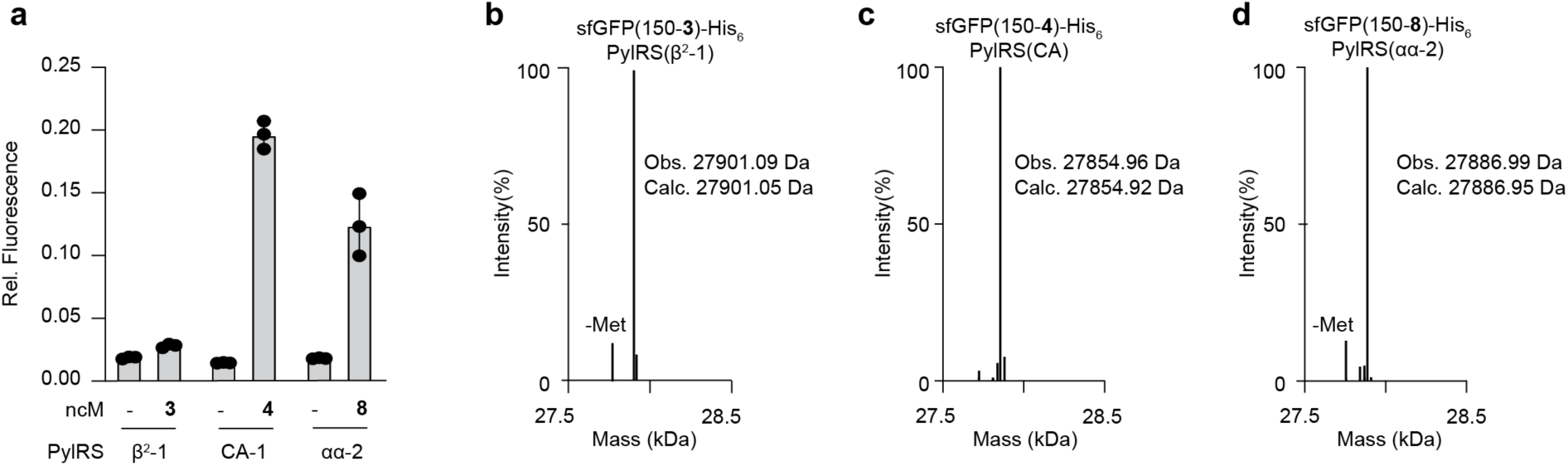
Site-specific incorporation of β^2^, N-cyclic, and α,α-disubstituted amino acids. **a**, GFP fluorescence from cells transformed with a sfGFP(150TAG)His_6_ reporter plasmid encoding for tRNA^Pyl^ and PylRS(β^2^-1), PylRS(CA-1), or PylRS(αα-2), expressed in the presence or absence of the respective substrates **3**, **4**, and **8** at 4 mM. Fluorescence is normalised to expression of sfGFP(150TAG)His_6_ with wt PylRS in the presence of **1** (2 mM). **b-d**, Deconvoluted mass spectra of sfGFP purified from cells harbouring a sfGFP(150TAG)His_6_ reporter, the indicated PylRS variant and grown in the presence of the corresponding ncM: **3** (**b**, calc. mass 27901.09 Da, found mass 27901.05 Da), **4** (**c**, calc. mass 27854.96 Da, found mass 27854.92 Da), and **8** (**d**, calc. mass 27886.99 Da, found mass 27886.95 Da).

Overall, these data demonstrate, to our knowledge, the first examples of the site-specific in vivo incorporation of non-canonical N-cyclic amino acids and β^2^ amino acids by cellular translation.

### Co-translational incorporation of ncMs is strongly sequence context dependent

We set out to investigate the extent of context dependence for ncM incorporation by the *E*. *coli* translational machinery for four different ncM classes (β^3^-amino acids, β^2^-amino acids, N-cyclic amino acids, and α,α−disubstituted amino acids), with four representative monomers (**2**, **3**, **4**, and **5**, using the evolved PylRS variants). While all of these monomers were incorporated at position 150 in GFP we had previously shown that monomers **2** and **5** were not incorporated in response to an amber codon at position 3 of GFP^15^. We tested the incorporation of **2**-**5** in response to the TAG codon at nine different positions, corresponding to nine different sequence contexts, in an sfGFP-His_6_ reporter gene (the positions of the amber codon in the sfGFP reporters 1-9 are: 39, 44, 50, 98, 109, 116, 150, 154, and 187)^6^. We compared ncM incorporation with incorporation of **1**, an α-amino acid which is efficiently acylated onto tRNA^Pyl^ by wt PylRS (**Fig. 3a-e**).

**Figure 3.**
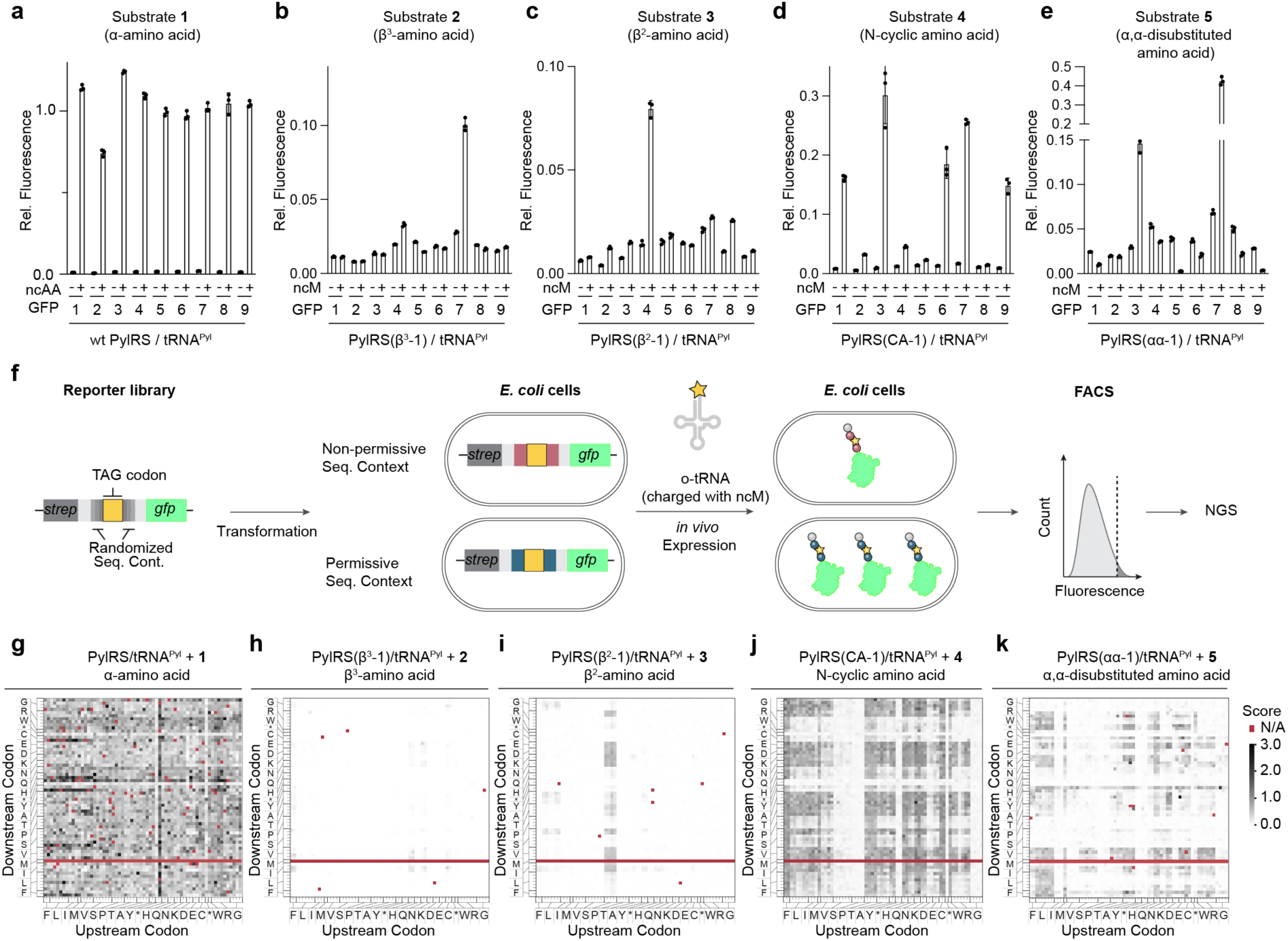
ncM incorporation efficiencies are strongly dependent on flanking codons. **a-e**, GFP fluorescence from cells transformed with sfGFP reporters containing a TAG codon at 9 different positions throughout the protein and a plasmid encoding for tRNA^Pyl^ and wt PylRS (**a**), PylRS(β^3^-1) (**b**), PylRS(β^2^-1) (**c**), PylRS(CA-1) (**d**), and PylRS(αα-1) (**e**) and expressed in the absence and presence of substrates **1** (**a**), **2** (**b**), **3** (**c**), **4** (**d**), or **5** (**e**). The positions of the TAG codon in the sfGFP reporters 1-9 are: 39, 44, 50, 98, 109, 116, 150, 154, and 187. **f**, Approach to exhaustively map the influence of codons directly before and after TAG codons for the incorporation efficiency of ncMs. A library in which the two codons adjacent to the TAG codon are varied to all possible codons (64×64=4096 library members) was transformed into cells harbouring an aaRS/tRNA pair which acylates the suppressor tRNA. After expression in the presence of the ncM, fluorescent cells were sorted by FACS and analysed by NGS, allowing identification of sequence contexts that likely support incorporation of the ncM. **g-k**, Incorporation efficiency heat maps for translation of monomers **1** with wt PylRS (**g**), **2** with PylRS(β^3^-1) (**h**), **3** with PylRS(β^2^-1) (**i**), **4** with PylRS(CA-1) (**j**), and **5** with PylRS(αα-1) mediated by tRNA^Pyl^. The sequence space is represented as a 64×64 matrix, in which the codon preceding TAG is represented on the x-axis and the codon following TAG on the y-axis. The 64 codons in each axis are arranged such that codons encoding the same amino acids are grouped. For the exact arrangement, see **Supplementary Fig. 24a**. Sequence contexts with an ATG codon following the TAG codon can lead to translation reinitiation and were excluded from analysis. Threshold values for selective incorporation of each ncM were determined to be 0.386 for (**g**), 0.125 for (**h**), 0.008 for (**i**), 0.216 for (**j**), 0.393 for (**k**).

Upon addition of **1,** we observed high, and comparable, GFP fluorescence levels with each of the 9 reporters; this confirmed that the corresponding TAG codon positions in GFP can support efficient and consistent levels of amber suppression in several sequence contexts (**Fig. 3a**). In contrast, for the β^3^-amino acid **2**, β^2^-amino acid **3**, N-cyclic amino acid **4**, and α,α-disubstituted amino acid **5** we observed predominantly no-or low-to-moderate-ncM-dependent increases in GFP fluorescence. A few positions supported much higher levels of incorporation than others; for each monomer the positions supporting higher levels of incorporation were distinct (**Fig. 3b-e**). In several cases, we observed a decrease in fluorescence upon addition of monomers. This was most notable for **5**, where 6/9 GFP reporters led to a decrease in GFP fluorescence upon addition of the ncM. These observations are consistent with the acylation of tRNA^Pyl^ with **5** outcompeting any background acylation with canonical amino acids and **5-**tRNA^Pyl^ being dominant negative in translational readthrough at these positions (eg: **5**-tRNA^Pyl^ might act as a chain terminator in translational readthrough). Overall, our results suggested that the in vivo incorporation of ncMs is very sensitive to the sequence context of the codon at which the ncM is incorporated.

Previously, sequence context has been observed to affect amber suppression efficiency in vivo^21–23^. As the co-translational incorporation of a monomer requires direct bond formation to the amino acid before and after it in the resulting polymer, the largest effects on backbone-modified monomer incorporation may be localized to the codons directly adjacent to the codon used to encode the monomer^24,25^. Therefore, we decided to directly map the influence of codons before and after the amber codon on the incorporation of each ncM using a library of codon context reporters.

We generated a library of codon context reporters in which a TAG codon was placed in a ten-amino acid stretch between the coding sequence of an N-terminal Strep-tag and the coding sequence of sfGFP; the codons immediately flanking the TAG codon were varied to all possible combinations (64×64; 4096 members) (**Fig. 3f**).

We first picked 58 random and distinct clones from this library and measured the GFP fluorescence resulting from each reporter in the presence of the PylRS tRNA^Pyl^ pair with or without **1.** All 58 codon contexts generated strong monomer dependent increases in fluorescence (**Supplementary Fig. 23a**), further supporting the permissivity of translation to this α-L amino acid across diverse sequence contexts. We used the 58 codon context reporters to measure GFP fluorescence with and without ncMs **2**, **3**, **4** and **5** in the presence of their respective PylRS variants and tRNA^Pyl^. We observed monomer dependent increases in GFP fluorescence for 8/58 contexts for **2**, 28/58 contexts for **3**, 45/58 contexts for **4** and, 8/58 codon context reporters for **5** (**Supplementary Fig. 23b-e**). In all tested sequence contexts, with all synthetase variants and ncMs the GFP signal in the absence of ncM was low, consistent with the observed specificity of the synthetases for acylating their tRNAs with ncMs. However, we note that for 45/58 sequence contexts the α,α-disubstituted amino acid **5** led to a decrease in GFP fluorescence upon addition of the ncM, consistent with our observations with the 9 GFP reporters with this ncM. Overall, our data further support the strong sequence context dependence for the incorporation of these ncMs.

Next, we expressed the sequence context reporter library in the presence of aaRS/tRNA pairs with or without the cognate ncM and analysed GFP fluorescence by flow cytometry. For the +ncM samples we also collected fluorescent cells by FACS (**Methods**) and sequenced the sorted population, as well as the input library, and used this data to calculate an incorporation efficiency score (abundance of a sequence in the sorted sample normalised by its input abundance) for each sequence context (**Supplementary Fig. 24**). We used this data to generate heatmaps of incorporation efficiency scores as a function of sequence context (**Fig. 3g-k**) For the α-amino acid **1**, 0.4% of cells passed a fixed FACS gate in the absence of **1**, while 83.37 ± 0.8% of cells passed the gate in the presence of **1 (Supplementary Fig. 25a)**. We saw an even distribution of incorporation efficiency scores across most sequence contexts (**Fig. 3g**, **Supplementary Fig. 25b**). 97.7% of sequence contexts were above a ‘threshold value’, an incorporation efficiency score value above that for sequences, within the 58 clones, for which we have directly measured a ncM-dependent GFP signal above the highest background level with the same synthetases/tRNA pair (**Methods)**. Sequence contexts with scores above the threshold value are likely to exhibit ncM-dependent readthrough. These observations suggested that, as expected, most sequence contexts support the incorporation of **1**. In contrast, for the β^3^-amino acid the percentage of cells that passed the gate for fluorescence was comparable and low without and with addition of ncM **2** (**Supplementary Fig. 25c,** 0.7 ± 0.1% and 0.6% respectively) and only 0.6% of sequence contexts were above the threshold value (**Fig. 3h**, **Supplementary Fig. 25d**). These observations suggested that very few sequence contexts support the incorporation of **2**. For the β^2^-amino acid 0.4% of cells passed the gate in the absence of **3** and 2.9 ± 0.1% of cells passed the gate in the presence of **3**, 26.3% of sequence context were above the threshold value (**Fig. 3i**, **Supplementary Fig. 25e-f**); these data suggested most sequence contexts do not support incorporation of **3**. For the PylRS(CA-1)/tRNA^Pyl^ pair, 0.3% of cells passed the gate in the absence of **4** and 36.9 ± 1.2% of cells passed the gate in presence of **4** (**Supplementary Fig. 25g**). 52.0% of sequence contexts were above the threshold value (**Fig. 3j**, **Supplementary Fig. 25h**); these data suggest that several sequence contexts support the incorporation of **4**. For the PylRS(αα-1)/ tRNA^Pyl^ pair, 30% of cells passed the gate in the absence of **5** and 18 ± 0.6% of cells passed the gate in the presence of **5**. The decrease in the fraction of cells passing the FACS gate in the presence of ncM **5** likely results from the dominant negative effect of tRNAs acylated with **5** on translational readthrough at many positions, as directly observed at 6/9 positions tested within GFP and 45/58 sequence context reporters (**Fig. 3e**, **Supplementary Fig. 23e**). 14.5% of sequence contexts were above the threshold value (**Fig. 3k, Supplementary Fig. 25i-j**).

Overall, our observations show that incorporation of the ncMs is much more strongly dependent on local sequence context than an alpha-L-amino acid control. The individual classes of ncMs display different sequence context preferences, and the incorporation efficiencies of some ncMs are strongly influenced by the identity of the preceding and subsequent amino acids. For example, for the β^2^-amino acid, **3**, we observed a strong preference for proline codons upstream of the TAG codon (**Fig. 3i**); Monomer **4** was incorporated at a wider range of sequence contexts than **2** and **3**, and was poorly incorporated when the upstream codons encoded glycine, serine or proline or when the first base of the downstream codon was uracil (**Fig. 3j**). Monomer **5** was incorporated at a limited number of diverse sequence contexts and was incorporated especially poorly when codons for small amino acids (glycine or serine) were upstream or downstream (**Fig. 3k**).

### Engineered tRNAs direct efficient ncM incorporation

Next, we aimed to improve the *in vivo* incorporation of ncMs and the diversity of accessible sequence contexts by engineering the translational machinery. We focused on engineering the orthogonal tRNA (**Fig. 4a**) as this is a central element of translation that is directly acylated with the ncM and interacts with several translational components (including: aaRS, EF-TU, ribosome, and mRNA) and determines the positioning of the monomer at the peptidyl transfer-center (PTC) of the ribosome. Since the orthogonal tRNA is involved in every step of translation, we reasoned that mutations in the orthogonal tRNA could potentially compensate for defects in translation with ncMs at any steps that were limiting incorporation^26^. Guided by the functions of distinct regions of the tRNA in translation, we constructed eight tRNA^Pyl^ saturation libraries, targeting positions known to interact with EF-TU (acceptor and the T-stem), as well as regions of the D-arm and the anticodon arms (**Supplementary Fig. 26**)^27–29^.

**Figure 4.**
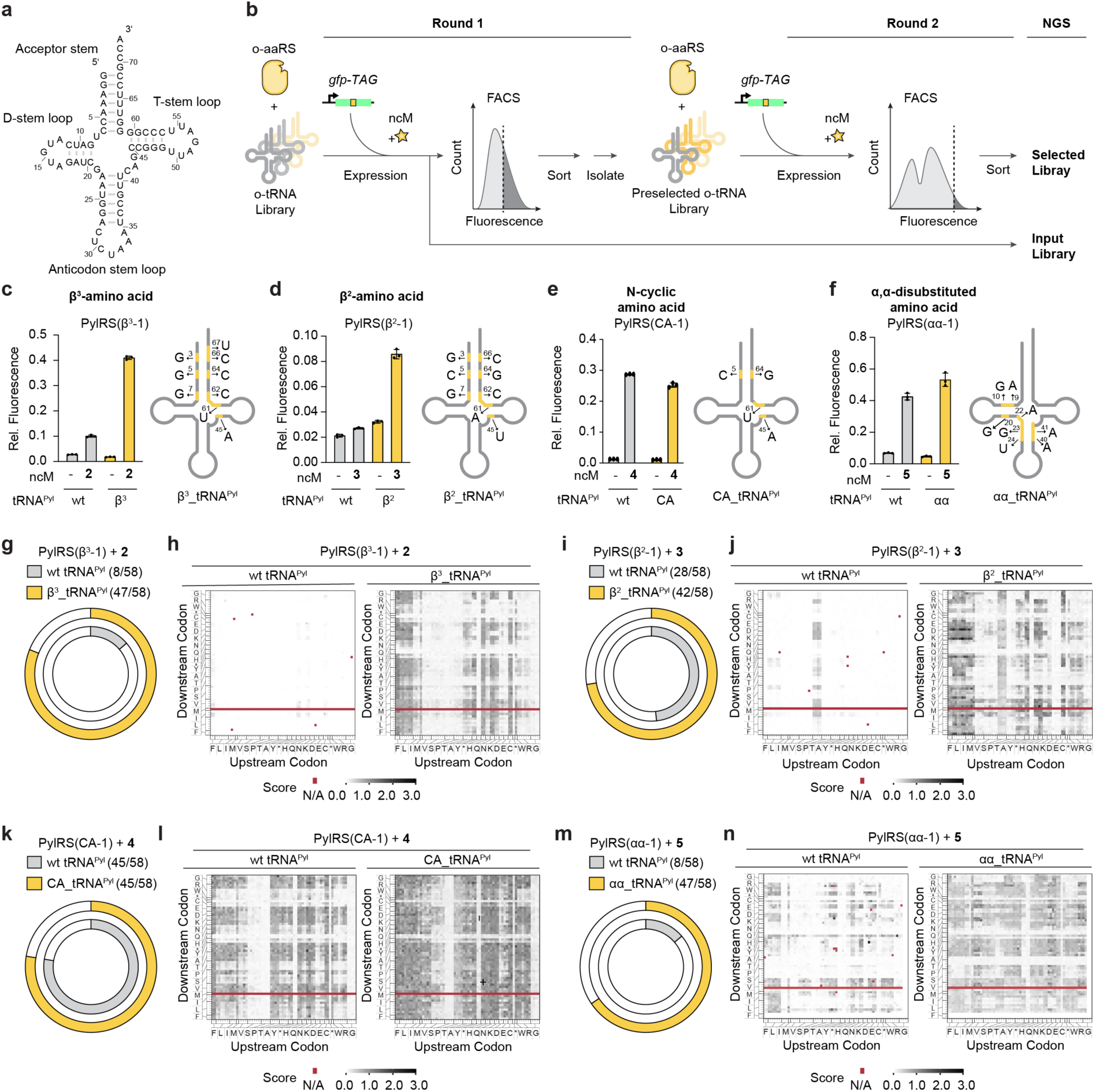
tRNA evolution increases in vivo incorporation of ncMs at diverse sequence contexts. **a**, Secondary structure of tRNA^Pyl^. **b**, FACS-based strategy for directed evolution of tRNAs. A tRNA library is expressed with its cognate aaRS and a fluorescence reporter in the presence of the corresponding ncM. Active sequences are sorted using FACS. The preselected library is isolated, retransformed into cells containing a new fluorescence reporter, and expressed in the presence of ncM. In a second selection step the most active sequences are sorted. The resulting populations and the input library are then analysed by next-generation sequencing in order to discover tRNA variants with the highest activity. **c-f** GFP fluorescence of cells transformed with wildtype and evolved tRNAs expressing a sfGFP(150TAG)His_6_ reporter. Cells were transformed with a sfGFP(150TAG)His_6_ reporter and a plasmid encoding for PylRS(β^3^-1) (**c**), PylRS(β^2^-1) (**d**), PylRS(CA-1) (**e**), or PylRS(αα-1) (**f**) and the indicated tRNAs. Protein expression was induced in the presence and absence of the respective ncMs **2-5**. Fluorescence was normalised to expression of sfGFP(150TAG)His_6_ with wt PylRS in the presence of **1** (2 mM). For each ncM-tRNA^Pyl^ pair, the mutations in the evolved tRNA^Pyl^ variant are shown. **g**, **i**, **k**, **m** Number of ncM-dependent sequence context reporters (out of 58 tested) with PylRS(β^3^-1) (**g**), PylRS(β^2^-1) (**i**), PylRS(CA-1) (**k**), and PylRS(αα-1) (**m**) expressed in the presence and absence of ncMs **2** (**g**), **3** (**i**), **4** (**k**), or **5** (**m**). For each PylRS variant expression was conducted both with wt tRNA^Pyl^ and the indicated tRNA^Pyl^ variant evolved for the target ncM. A sequence context reporter was deemed ncM-dependent if fluorescence values in the presence of the respective ncM were at least 1.2 times higher than the fluorescence values in the absence of the respective ncM. **h**, **j**, **l**, **n**, Incorporation efficiency heat maps for translation of substrates **2** with PylRS(β^3^-1) (**h**), **3** with PylRS(β^2^-1) (**j**), **4** with PylRS(CA-1) (**l**), and **5** with or PylRS(αα-1) (**n**). for each ncM-PylRS combination heat maps are shown for the wt tRNA^Pyl^ (left) or the corresponding evolved tRNA^Pyl^ (right). Threshold values for selective incorporation of each ncM with evolved tRNAs were determined to be 0.109 for (**h**), 0.137 for (**j**), 0.216 for (**l**), 0.401 for (**n**). Heat maps for wt tRNA^Pyl^ experiments correspond to data shown in Fig. 3g-k.

We first focused on discovering orthogonal tRNA^Pyl^ variants that enhance the incorporation of the β^3^-amino acid, **2**. To select tRNAs that promote efficient incorporation of **2** we performed eight parallel selections in cells harbouring PylRS(β^3^-1), a sfGFP(150TAG)His_6_ reporter and, each tRNA library. After inducing sfGFP expression in the presence of **2**, we selected for fluorescent cells by FACS. After a low stringency round of sorting, to enrich active library members, we performed a second – more stringent – round, in which we sorted the most fluorescent cells (**Fig. 4b)**.

After the first round of sorting, we decided to focus on library 1 and library 3, which had the largest population of strongly fluorescent cells in the presence of **2** and very few fluorescent cells in the absence of **2** (**Supplementary Fig. 27**). Following the second round of selection, we performed NGS on the +ncM sample and the input from the first round and, used this data to calculate the enrichment of each tRNA. For both libraries (1 and 3) the most enriched tRNA sequences contained clear sequence motifs (**Fig. 4c**, **Supplementary Fig. 28a-b**); these sequences preserved the topology of the tRNA cloverleaf in secondary structure predictions.

The most active tRNA, β^3^_tRNA^Pyl^, derived from library 3, led to 4-fold greater fluorescence than wildtype tRNA^Pyl^ when cells were provided with an sfGFP(150TAG)His_6_ reporter, PylRS(β^3^-1) and **2**. All selected tRNAs maintained strong monomer-dependent fluorescence (**Supplementary Fig. 28c**); indeed, several evolved tRNAs showed reduced GFP fluorescence in the absence of **2**, when compared to their parental tRNA. These results are consistent with the evolved tRNAs increasing the efficiency of incorporation of **2** (**Fig. 4c**). We confirmed high fidelity, site-specific incorporation of monomer **2** by mass spectrometry (**Supplementary Fig. 28d**).

Next, we investigated whether β^3^_tRNA^Pyl^ could increase the efficiency or fidelity of incorporation for other β^3^ amino acids (**S21-S25**) for which we have previously discovered aminoacyl-tRNA synthetases. We combined PylRS(β^3^-2)–PylRS(β^3^-6) with (or without) their cognate β^3^ amino acids (**S21-S25, Supplementary Fig. 29a**), with the sfGFP(150TAG)His_6_ reporter and either wt tRNA^Pyl^ or β^3^_tRNA^Pyl^. ncMs **S22** and **S25** have previously been incorporated into proteins, while **S21**, **S23** and **S24** can by acylated on to wt tRNA^Pyl^ but have not been incorporated into proteins. We observed higher, and more ncM-dependent, fluorescence with β^3^_tRNA^Pyl^ for ncMs **S21-S23** and **S25** (up to 7-fold higher than with wt tRNA^Pyl^, in the case of **S25**) (**Supplementary Fig. 29b**). Intact protein mass via ESI-MS. demonstrated the incorporation of **S21**, **S22**, **S23**, and **S25** at position 150 of sfGFP (**Supplementary Fig. 29c-f**). Our data demonstrates that β^3^_tRNA^Pyl^ functions with additional PylRS variants to enable the incorporation of β^3^ amino acids that could not previously be incorporated.

Encouraged by our results, we set out to evolve tRNA^Pyl^ for increased incorporation of the ncMs **3**, **4**, and **5**. For the β^2^-amino acid **3**, we identified β^2^_tRNA^Pyl^, from library 3, which led to a four-fold increase in fluorescence with respect to wildtype tRNA^Pyl^ when cells were provided with an sfGFP(150TAG)His_6_ reporter, PylRS(β^2^-1) and **3 (Fig. 4d**, **Supplementary Fig. 30a-d).** We confirmed incorporation of **3,** using this new tRNA, by mass spectrometry (**Supplementary Fig. 30e**). β^2^_tRNA^Pyl^ increased the fidelity of incorporation for additional β^2^-amino acids (**Supplementary Fig. 31**). For the N-cyclic amino acid **4**, we did not find tRNA^Pyl^ variants that increased monomer-dependent fluorescence levels with the sfGFP(150TAG)His_6_ reporter, when compared to wt tRNA^Pyl^ (**Fig. 4e**, **Supplementary Fig. 32a-d**). However, we identified CA_tRNA^Pyl^, from library 3, which increased ncM-dependent fluorescence levels in the presence of **4** when challenged with a more stringent sfGFP reporter (**Supplementary Fig. 32e**). We confirmed incorporation of **4,** using this new tRNA, by mass spectrometry. CA_tRNA^Pyl^ also increased the efficiency of incorporation for the additional N-cyclic amino acid **S6 (Supplementary Fig. 33)**. For ncM **5**, the α,α-disubstituted amino acid, we identified αα_tRNA^Pyl^, from library 6 **(Supplementary Fig. 34a-h)**. This tRNA increased monomer dependent fluorescence, and also decreased background fluorescence, with the sfGFP(150TAG)His_6_ reporter, when compared to wt tRNA^Pyl^ (**Fig. 4f**). We confirmed the site-specific incorporation of **5,** using this tRNA, by mass spectrometry (**Supplementary Fig. 34i**). Overall, the evolved tRNA^Pyl^ variants mediate higher efficiency incorporation for monomers of the corresponding ncM class and in some cases the evolved tRNAs enabled the first incorporation of particular monomers into proteins.

### Evolved tRNAs substantially expand the sequence contexts that support ncM incorporation

Next, we investigated whether the evolved tRNAs expanded the sequence contexts that support the incorporation of ncMs. To do this we investigated the incorporation of each ncM using: 1) GFP expression experiments with the 9 GFP reporters containing internal amber codons, 2) GFP expression experiments with the 58 selected codon context reporters and, 3) FACS-based sequence context incorporation mapping using the complete codon context reporter library.

β^3^_tRNA^Pyl^ increased the number of GFP reporters with internal amber codons that exhibited ncM-dependent increases in fluorescence from 2/9, with wt tRNA^Pyl^, to 7/9 (**Supplementary Fig. 35a**). Similarly, β^3^_tRNA^Pyl^ increased the fraction of the 58 codon context reporters that exhibited ncM-dependent fluorescence from 8/58, with wt tRNA^Pyl^, to 47/58 (**Fig. 4g**). The evolved tRNA also substantially increased the level of GFP produced in the sequence contexts that previously exhibited ncM-dependent fluorescence with wt tRNA^Pyl^ (2/2 internal codons up to 4.9 fold, and 8/8 codon context reporters up to 7.2 fold, **Supplementary Fig. 35a-c**). In FACS-based mapping experiments, the percentage of cells bearing the PylRS(β^3^-1)/β^3^_tRNA^Pyl^ pair, that passed the gate for fluorescence in the presence of **2** increased (from 0.7% ± 0.1 for wt tRNA^Pyl^) to 24.1 ± 0.3% of cells; while only 0.9% (compared to 0.4% for wt tRNA^Pyl^) of cells passed the gate in the absence of **2** (**Supplementary Fig. 35d**). The percentage of sequence contexts with an incorporation efficiency score above the threshold for background was 55.6% (compared to 0.6% for wt tRNA^Pyl^) (**Fig. 4h**, **Supplementary Fig. 35e**). We observed a strong influence of the upstream codon on incorporation efficiency scores for **2** with β^3^_tRNA^Pyl^; the incorporation efficiency score was higher when the upstream codon encodes large, hydrophobic amino acids (phenylalanine, leucine, isoleucine, methionine, tyrosine, tryptophan) and certain polar amino acids (glutamine and cysteine). This suggested that these sequence contexts support the incorporation of **2** with β^3^_tRNA^Pyl^.

β^2^_tRNA^Pyl^ did not increase the number of GFP reporters with internal amber codons that exhibited ncM-dependent increases in fluorescence (8/9 as with wt tRNA^Pyl^) (**Supplementary Fig. 36a**). However, β^2^_tRNA^Pyl^ increased the fraction of the 58 codon context reporters that exhibited ncM-dependent fluorescence from 28/58, with wt tRNA^Pyl^, to 42/58 (**Fig. 4i**). The evolved tRNA also substantially increased the level of GFP produced in the sequence contexts that previously exhibited ncM-dependent fluorescence with wt tRNA^Pyl^ (8/8 internal codons up to 8.1 fold, and 28/28 codon context reporters up to 10.8 fold, **Supplementary Fig. 36a-c**). In FACS-based mapping experiments, the percentage of cells bearing the PylRS(β^2^-1)/β^2^_tRNA^Pyl^ pair, that passed the gate for fluorescence in the presence of **3** increased (from 2.9 ± 0.1% for wt tRNA^Pyl^) to 31.1 ± 0.3% of cells; while 10.8% (compared to 0.4% for wt tRNA^Pyl^) of cells passed the gate in the absence of **3** (**Supplementary Fig. 36d**). The percentage of sequence contexts with an incorporation efficiency score above the threshold for background was 54.6% (compared to 26.3% for wt tRNA^Pyl^) (**Fig. 4j**, **Supplementary Fig. 36e**). The incorporation efficiency scores suggested than incorporation of **3** with β^2^_tRNA^Pyl^, was favoured when the upstream codon encoded a hydrophobic amino acid or cysteine, and incorporation was disfavoured when the downstream codon encoded tryptophan or threonine.

CA_tRNA^Pyl^ showed ncM-dependent increase in fluorescence for all nine GFP reporters with internal amber codons, the same as wt tRNA^Pyl^. (**Supplementary Fig. 37a**). Similarly, CA_tRNA^Pyl^ did not increase the fraction of the 58 codon context reporters that exhibited ncM-dependent fluorescence from 45/58, with wt tRNA^Pyl^, (**Fig. 4k**). However, the evolved tRNA substantially increased the level of GFP produced upon addition of **4** in the sequence contexts that previously exhibited ncM-dependent fluorescence with wt tRNA^Pyl^ (4/9 internal codons up to 1.6 fold, and 45/45 codon context reporters up to 2.5 fold, **Supplementary Fig. 37a-c**). In FACS experiments, the percentage of cells bearing the PylRS(CA-1)/CA_tRNA^Pyl^ pair, that passed the gate for fluorescence in the presence of **4** increased (from 36.1 ± 1.2% for wt tRNA^Pyl^) to 56.0 ± 0.5% of cells; while only 0.4% (compared to 0.3% for wt tRNA^Pyl^) of cells passed the gate in the absence of **3** (**Supplementary Fig. 37d**). The percentage of sequence contexts with an incorporation efficiency score above the threshold for background was 78.8% (compared to 52.0% for wt tRNA^Pyl^) (**Fig. 4l**, **Supplementary Fig. 37e**). The incorporation efficiency scores for CA_tRNA^Pyl^ with **4** were low when the upstream codon encoded glycine or proline; this suggested that these sequence contexts disfavour incorporation of **4** with CA_tRNA^Pyl^.

αα_tRNA^Pyl^ increased the number of GFP reporters with internal amber codons that exhibited ncM-dependent increases in fluorescence from 2/9, with wt tRNA^Pyl^, to 6/9 (**Supplementary Fig. 38a**). Similarly, αα_tRNA^Pyl^ increased the fraction of the 58 codon context reporters that exhibited ncM-dependent fluorescence from 8/58, with wt tRNA^Pyl^, to 38/58 (**Fig. 4m**).

αα_tRNA^Pyl^ also substantially increased the level of GFP produced in the sequence contexts that previously exhibited ncM-dependent fluorescence with wt tRNA^Pyl^ (2/2 internal codons up to 2.1 fold, and 8/8 codon context reporters up to 4.8 fold, **Supplementary Fig. 38a-c**). In FACS experiments, the percentage of cells bearing the PylRS(αα-1)/αα_tRNA^Pyl^ pair, that passed the gate for fluorescence in the presence of **5** increased (from 18.0% ± 0.6 for wt tRNA^Pyl^) to 45.9 ± 2.6% of cells; while 10.2% (compared to 30.0% for wt tRNA^Pyl^) of cells passed the gate in the absence of **5** (**Supplementary Fig. 38d**). αα_tRNA^Pyl^ decreases background incorporation in the absence of ncM for a range of sequence context reporters (**Supplementary Fig. 38a-b**). This is reflected in the decrease in the fraction of cells bearing the sequence context library that pass the fluorescence gate in the absence of **5** compared to wt tRNA^Pyl^. For αα_tRNA^Pyl^ the percentage of sequence contexts with an incorporation efficiency score above the threshold for background was 48.7% (compared to 14.5% for wt tRNA^Pyl^) (**Fig. 4n**, **Supplementary Fig. 38e**). The evolved αα_tRNA^Pyl^ showed more uniform incorporation efficiency scores across sequence contexts than wildtype tRNA^Pyl^ for incorporation of ncM **5**. However, our data suggest that sequence contexts with downstream proline and tryptophan codons disfavour the incorporation of **5** with αα_tRNA^Pyl^.

### Further expanding the sequence contexts that support ncM incorporation

While the incorporation of ncMs at diverse sequence context was enabled by the evolved tRNAs, certain sequence contexts remained recalcitrant to efficient ncM incorporation. We hypothesized that selecting tRNA variants on additional sequence context reporters might enable incorporation of ncMs at sequence contexts that were otherwise recalcitrant to ncM incorporation. We investigated this hypothesis for the incorporation of the β^3^-amino acid **2.**

We chose four N-terminal sfGFP variants s.c.1 (CAG-TAG-TGC, Gln-X-Cys), s.c.2 (GCC-TAG-GCC, Ala-X-Ala), s.c.3 (AAC-TAG-CAA, Asn-X-Gln) s.c.4 (CAC-TAG-TCG, His-X-Ser), which lead to variable incorporation of the β^3^-amino acid **2** with the PylRS(β^3^-1)/ β^3^_tRNA^Pyl^ pair, as sequence context reporters (**Fig. 5a**). We constructed four libraries on top of β^3^_tRNA^Pyl^ targeting complementary parts of the tRNA (Lib β^3^_1-4) (**Supplementary Fig. 39**). We performed FACS based selections with each reporter, PylRS(β^3^-1) and the tRNA libraries (**Supplementary Figs. 40-42**).

**Figure 5.**
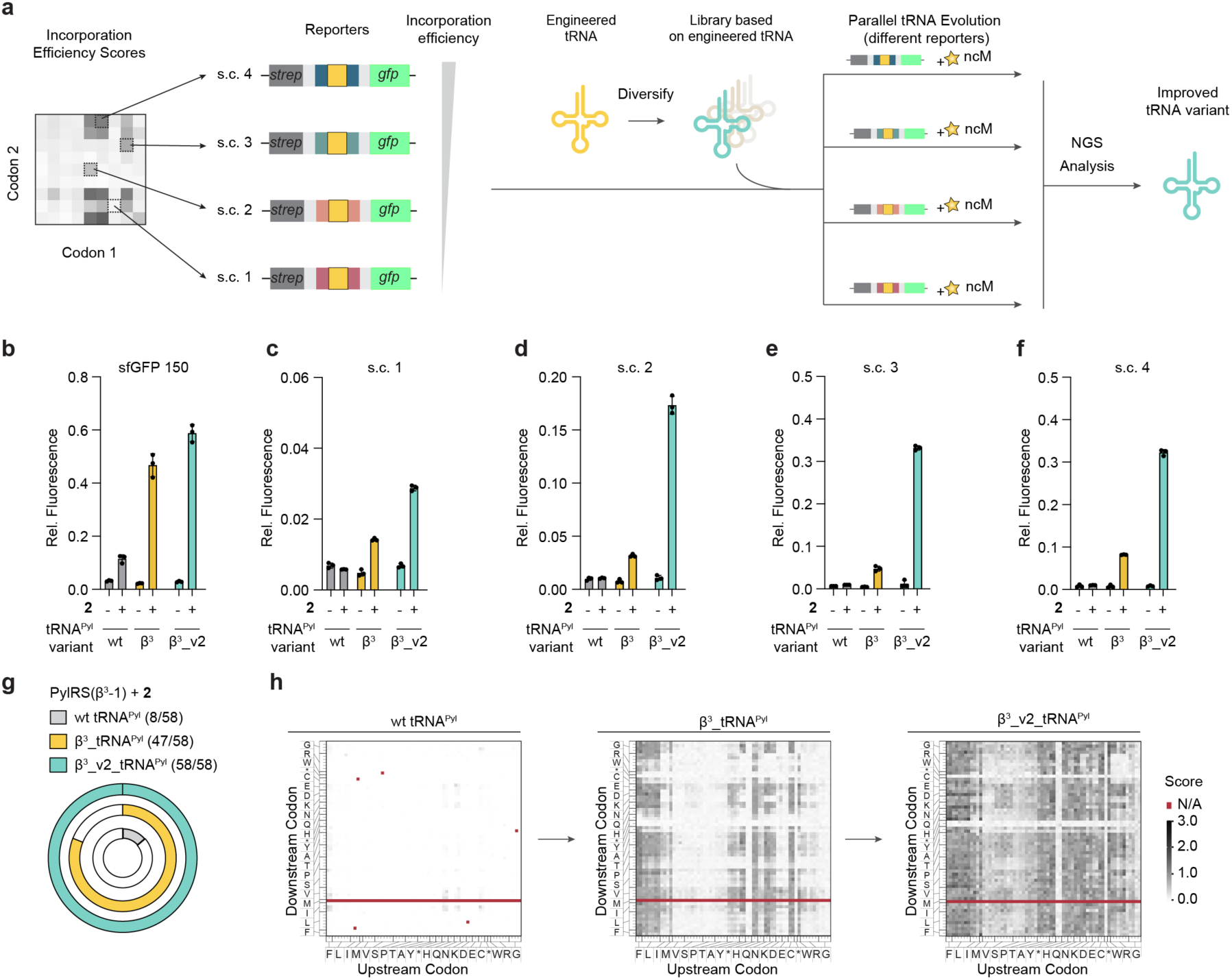
tRNAs selected on additional sequence contexts expand the scope of ncM incorporation. **a**, Sequence contexts with varying degrees of incorporation efficiency for a specific ncM can be derived from sequence context mapping data or fluorescence values measured directly from clones of a sequence context collection. Different sequence contexts can be used for the evolution of tRNAs in parallel to tune selection stringency. Selections were run with substrate **2**, PylRS(β^3^-1) and four sequence context reporters with varying degrees of incorporation efficiencies with β^3^_tRNA^Pyl^. Libraries were built on top of β^3^_tRNA^Pyl^. **b-f**, GFP fluorescence values of the most active tRNA sequence from parallel selections run with libraries β^3^_2 and β^3^_3 using sequence context reporters s.c. 1, s.c. 2, s.c. 3, and s.c.4. Plasmids encoding for PylRS(β^3^-1) and the indicted tRNA variants were transformed into cells harbouring reporter plasmids encoding for: sfGFP(150TAG)His_6_ (**b**), s.c. 1 (**c**), s.c. 2 (**d**), s.c. 3 (**e**), or s.c. 4 (**f**). Expression was conducted in the presence or absence of **2**. Fluorescence was normalised to expression of the sfGFP(150TAG)His_6_ reporter with wt PylRS and **1** (2 mM). **g**, Fraction of ncM-dependent sequence context reporters (out of 58 tested) with PylRS(β^3^-1) and wt tRNA^Pyl^, β^3^_tRNA^Pyl^ and β^3^_v2_tRNA^Pyl^ expressed in the presence and absence of **2**. A sequence context reporter was deemed ncM-dependent if fluorescence values in the presence of the respective ncM were at least 1.2 times higher than the fluorescence values in the absence of the respective ncM. The data for wt tRNA^Pyl^ and β^3^_tRNA^Pyl^ are duplicated from Fig. 4g. **h**, Incorporation efficiency heat maps for translation of substrate **2** mediated by PylRS(β^3^-1) and a tRNA^Pyl^ variant: wt tRNA^Pyl^ (left), β^3^_tRNA^Pyl^ (middle), and β^3^_v2_tRNA^Pyl^ (right). The threshold value for selective incorporation with β^3^_v2_tRNA^Pyl^ was determined to be 0.373. The data for wt tRNA^Pyl^ and β^3^_tRNA^Pyl^ are duplicated from Fig. 4h.

We screened enriched hits from these selections and ultimately identified β^3^_v2_tRNA^Pyl^ (**Supplementary Figs. 43-48, Methods**). This tRNA enabled a strong increase in fluorescence for the sequence context reporters (s.c.1-4) in the presence of **2**. For example, for s.c.4, β^3^_v2_tRNA^Pyl^ increased the ncM-dependent fluorescence 5-fold with respect to β^3^_tRNA^Pyl^ and 40-fold with respect to wt tRNA^Pyl^ (**Fig. 5b-f**). We confirmed the incorporation of **2** into s.c.2-s.c.4 reporters with the PylRS(β^3^-1)/β^3^_v2_tRNA^Pyl^ pair by ESI-MS (**Supplementary Fig. 47**).

Next, we tested the influence of β^3^_v2_tRNA^Pyl^ and other hits on the incorporation of **2** in the nine different sfGFP reporters with internal amber codons. We observed substantially increased fluorescence levels (up to 12-fold) in samples expressed in the presence of β^3^_v2_tRNA^Pyl^ compared to samples expressed in the presence of β^3^_tRNA^Pyl^ for six of the nine positions (**Supplementary Fig. 48b**). All 58 codon context reporters exhibited ncM-dependent fluorescence with β^3^_v2_tRNA^Pyl^; up from 47, with β^3^_tRNA^Pyl^ and 8 for wt tRNA^Pyl^ (**Fig. 5g**). β^3^_v2_tRNA^Pyl^ increased fluorescence signal for all 47 experimentally tested sequence context reporters that were previously accessible with β^3^_tRNA^Pyl^; the increase in fluorescence was up to 11.4-fold with respect to β^3^_tRNA^Pyl^, and up to 28-fold with respect to wt tRNA^Pyl^ (**Supplementary Fig. 49b**). In FACS-based mapping experiments, the percentage of cells bearing the PylRS(β^3^-1)/β^3^_v2_tRNA^Pyl^ pair, that passed the gate for fluorescence in the presence of **2** increased to 60.9 ± 0.5% of cells (from 24.1 ± 0.3% for β^3^_tRNA^Pyl^ and 0.7 ± 0.1% for wt tRNA^Pyl^); while only 5.3% (compared to 0.9% for β^3^_tRNA^Pyl^ and 0.4% for wt tRNA^Pyl^) of cells passed the gate in the absence of **2** (**Supplementary Fig. 49c**). The percentage of sequence contexts with an incorporation efficiency score above the threshold for background was 95.5% (compared to 55.6% for β^3^_tRNA^Pyl^ and 0.6% for wt tRNA^Pyl^) (**Fig. 5h**, **Supplementary Fig. 49e**). We conclude that β^3^_v2_tRNA^Pyl^, enables the incorporation of **2** at a wider range of sequence contexts.

In additional experiments, we demonstrated the generality of our approach by selecting αα_v2_tRNA^Pyl^ on an additional sequence context reporter (s.c.50, CAA-TAG-AAA, Gln-X-Lys); this evolved tRNA further increased the range of sequence contexts at which the α,α-disubstituted ncM **5** can be efficiently and specifically incorporated (**Supplementary Figs. 50-51**). Overall, we conclude that by selecting tRNAs, from libraries of variants, on distinct sequence contexts we can further expand the range of sequence contexts that support efficient and specific ncM incorporation.

### Genetically programmed cell-based synthesis of macrocycles with non-canonical backbones

To demonstrate the utility of our ncM incorporation systems we realized the genetically encoded cell-based synthesis of macrocycles^5,6^ bearing β^3^-amino acids, β^2^-amino acids, non-canonical N-cyclic amino acids and α,α-disubstituted amino acids. We detected formation of macrocycles CLSL-**2**-V (calc. *m/z* 741.2640, found *m/z* 741.2657), CLSL-**3**-V (calc. *m/z* 721.3589, found *m/z* 721.3567), CLSL-**4**-V (calc. *m/z* 675.3534, found *m/z* 675.3545), and CLSL-**5**-V (calc. *m/z* 803.2657, found *m/z* 803.2659) by ESI-MS (**Fig. 6a-d**). Incorporation of ncMs in the macrocycles was supported by MS/MS fragmentation data matching all permutations of linearized sequences (**Supplementary Fig. 52)**. These observations demonstrated that we can directly generate genetically encoded peptides with a diversity of altered backbones in living cells.

**Figure 6.**
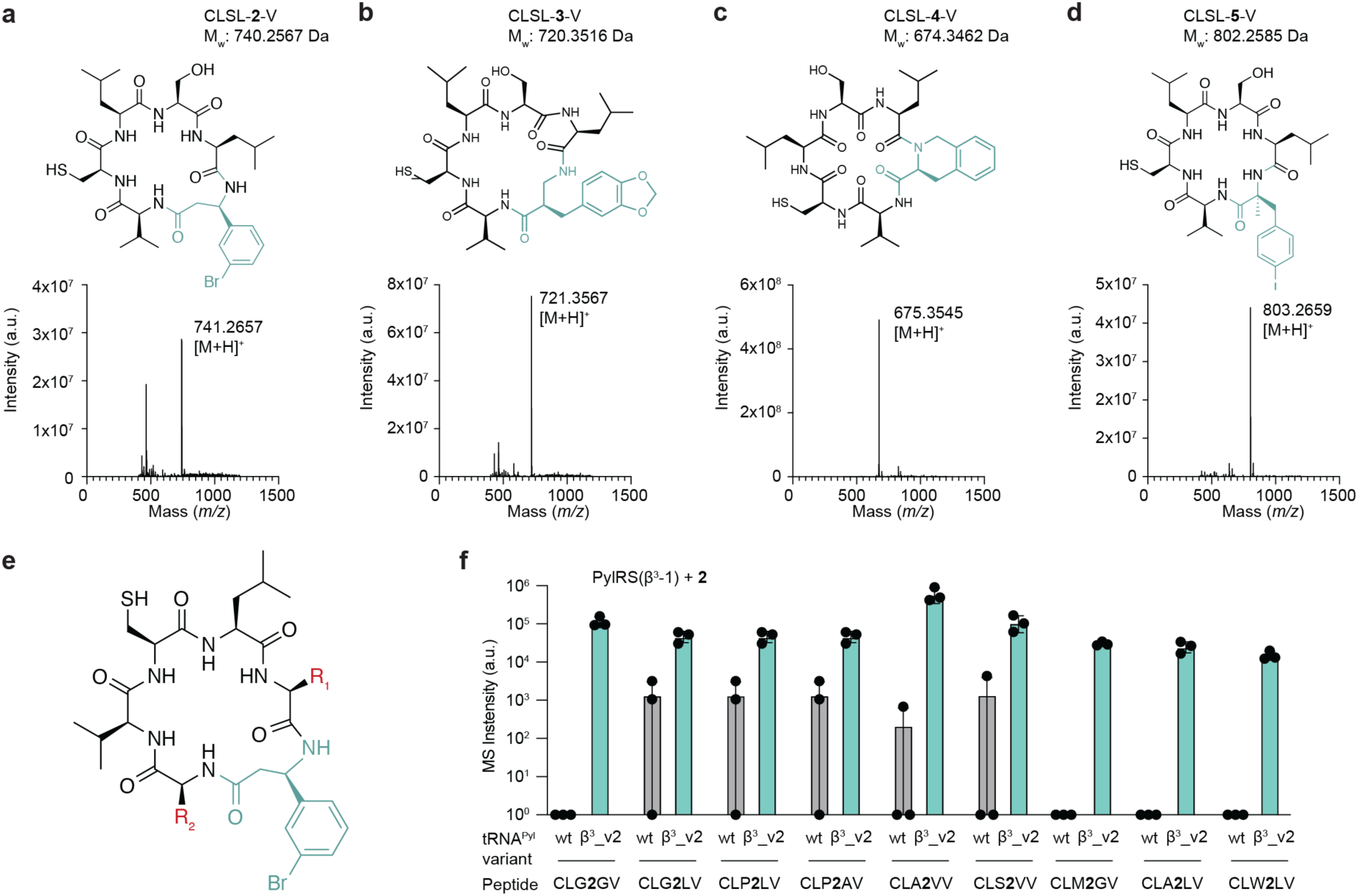
Genetically programmed cell-based production of macrocycles containing ncMs. **a**-**d**, Structures (above) and MS1 spectra (below) of cyclic CLSL-ncM-V peptides where the ncM corresponds to **2** (**a**), **3** (**b**), **4** (**c**), or **5** (**d**). The respective ncM is highlighted in green in the structure. **e**, Chemical structure of cyclic peptide variants. 32 distinct variants with different side chains R_1_ and R_2_ were expressed in the presence of PylRS(β^3^-1), **2** and wt tRNA^Pyl^ or β^3^_v2_tRNA^Pyl^. For a list of the 32 variants tested, see **Supplementary Data 1**. **f**, Area under the curve of MS intensities for masses matching indicated distinct peptides expressed in the presence of PylRS(β^3^-1), **2** and wt tRNA^Pyl^ or β^3^_v2_tRNA^Pyl^.

To directly demonstrate the ability of the evolved tRNAs to enhance access to genetically encoded macrocycles containing ncMs in distinct sequence contexts we picked 32 random independent clones from a library encoding macrocyclic peptide variants CLX_1_-ncM-X_2_V (where X_1_ and X_2_ are encoded by NNN and the ncM by TAG, **Fig. 6e**).

For PylRS(β^3^-1) and β^3^_v2_tRNA^Pyl^ grown in the presence of **2** we could purify and identify purified peptides from 9 of the library clones by mass spectrometry (**Fig. 6f,g**; **Supplementary Fig. 53**). For PylRS(β^3^-1) and wt tRNA^Pyl^ grown in the presence of **2** we could purify and identify peptides from 5 clones. The five clones identified with wt tRNA^Pyl^ were a subset of the 9 clones identified with β^3^_v2_tRNA^Pyl^ and these peptides were detected at lower abundances when produced using wt tRNA^Pyl^. Four of the nine library clones, which generated the corresponding peptides with β^3^_v2_tRNA^Pyl^, did not measurably generate the same peptides with wt tRNA^Pyl^. We note that we did not expect to purify all 32 sequences (our purification conditions favour the purification of hydrophobic peptides). The experiment allows us to compare the relative abundance of each peptide that is produced and can be purified under the conditions used. These experiments demonstrated that the evolved tRNAs enabled incorporation of ncMs into macrocycles at a wider range of sequence contexts and with greater efficiency.

## Discussion

Our work explicitly demonstrates that the discovery of orthogonal synthetases that acylate their cognate tRNAs with ncMs enables the co-translational selection of translational components that facilitate the incorporation of ncMs. We demonstrate the utility of this paradigm through the evolution of tRNAs that enable: 1) the incorporation of previously unincorporated ncMs, 2) the incorporation of ncMs with much higher efficiency, 3) the incorporation of ncMs across many more sequence contexts, and 3) direct access to cellular synthesis of ncM containing macrocyclic library members that could not be accessed without evolved tRNAs. Orthogonal tRNA evolution is a powerful approach for enhancing ncM incorporation because orthogonal tRNAs participate in essentially every step in translation and their evolution ensures they are optimized for the process of translation, rather than particular sub-steps of the overall process.

Future work may expand upon the paradigm reported herein to engineer other translational components, including orthogonal ribosomes^30–32^ and EF-Tu^33,34^, for ncM incorporation and polymerization.

The ability to efficiently incorporate diverse ncMs at diverse sequence contexts expands the chemical space for innovation in living organisms. These advances will allow the free composition, design^35^ and selection of peptides, proteins and polymers with new functions and properties, including protease resistance, new catalytic and binding activities and alternative structural motifs and folds^36–38^.

## Methods

### Buffers

Buffer Res: 100 mM sodium acetate, 50 mM NaCl, 1 mM EDTA, pH 5.0; 10x deacylation buffer: 500 mM bicine, 1 mM EDTA, pH 9.6; 10x annealing buffer: 10 mM Tris-HCl, 25 mM NaCl, pH 7.4; tREX loading dye: 8 M urea, orange G; Base Buffer: 1.6 M Na_2_CO_3_; Buffer W-T: 10 mM Tris-HCl, 150 mM LiCl, 1 mM EDTA, 0.05% (v/v) Tween-20, pH 7.5; binding buffer: 20 mM Tris-HCl, 1 M LiCl, 2 mM EDTA, pH 7.5; buffer B-T: 25 mM NaOH, 4 mM EDTA, 0.05% (v/v) Tween-20; RT-Hybridisation buffer: 1 μL DNA primer (2 μM), 1 μL 10 mM dNTPs, 1 μL 10x hybridization buffer, 10 μL water; RT-MM: 4 μL SSIV buffer, 1 μL RNAse Out (Invitrogen), 1 μL SSIV reverse transcriptase (Invitrogen), 1 μL 0.1 M dithiothreitol (DTT); buffer D: 50 mM sodium acetate pH 5, 150 mM NaCl, 10 mM MgCl_2_, 0.1 mM EDTA; RNaseA elution buffer: 20 mM NH_4_OAc, pH 5, 1.5 U/μL RNAseA (Thermoscientific); acW1-T: 100 mM ammonium acetate, pH 5.0, 0.05% (v/v) Tween-20; acW1: 100 mM ammonium acetate, pH 5.0; acW2: 20 mM ammonium acetate, pH 5.0; lysis buffer: 1x PBS, 1x BugBuster, 20 mM imidazole, 100 μg/mL lysozyme, 50 μg/mL DNAse; peptide lysis buffer: 50 mM HEPES, 150 mM NaCl, 10 mM imidazole, pH 8.0, containing 0.1 mg ml^-1^ lysozyme; NHI-30: (50 mM HEPES, 150 mM NaCl, 30 mM imidazole, pH 8.0); NHI-250: (50 mM HEPES, 150 mM NaCl, 250 mM imidazole, pH 8.0); MOPS reaction buffer: 20 mM MOPS, 150 mM NaCL, pH 6.9.

### Media

SOC: Super Optimal Broth + 20 mM glucose; 2xYT-s: 2xYT medium + 75 µg/mL spectinomycin sulphate; 2xYT-c: 2xYT medium + 100 µg/mL carbenicillin disodium salt; 2xYT-s-ap: 2xYT medium + 50-75 µg/mL spectinomycin sulphate + 50 µg/mL apramycin

### Plasmid construction

Plamids were assembled from PCR products using NEBuilder HiFi DNA Assembly Master Mix (NEB) according to manufacturer’s guidelines. Libraries were generated amplifying a template plasmid by PCR using two primers (see primer list) containing degenerate codons at desired mutagenesis sites and a type IIS restriction site. In the case of custom mixes, primers containing different codons were manually mixed and used for PCR reactions. Samples were purified, ligated using T4 DNA Ligase, and transformed into electrocompetent *E. coli* DH10β cells ensuring a minimal transformation efficiency of 10^9^. Plasmid DNA was prepared from the resulting culture, sequenced as a bulk using Sanger sequencing and used for subsequent experiments.

### tRNA display selections

The previously described library 14 encoded as a stmRNA construct on a pColE1 plasmid^15^ was transformed into electrocompetent Bl21 derived cells. Between 3-5 µg of plasmid was transformed in 3-5 separate electroporations each of which was recovered in 1 mL SOC for 1 h. After recovery, cells were diluted into 1 L of 2xYT-c and grown overnight at 37 °C, 220 rpm.

A fraction of the library was plated to determine transformation efficiency (minimum transformation efficiency >10^9^).

Cells were diluted 1:50 in 2xYT-c and grown to OD_600_ = 0.3. Then substrate was added to 2.5 mL of culture to a final concentration of 2 mM. This was conducted in 4 independent replicates. Cultures were grown for 40 min at 37 °C, 220 rpm in the presence of amino acid. Then cells were induced by addition of Isopropyl β-D-1-thiogalactopyranoside (IPTG) to a final concentration of 1 mM and incubated for 20 min at 220 rpm at 37 °C. Cells were harvested by centrifugation at 4200 rcf at 4 °C for 10 min. Cells were resuspended in 800 µL buffer RES and centrifuged again at the same speed and temperature. RNA isolation was conducted with an Agencourt RNAdvance Cell v2 RNA Isolation kit (Beckman Coulter). Cells were lysed by resuspension in 220 µL LBE buffer and 10 µL Proteinase K and incubated for 1 h at room temperature. Then 244 µL BBC beads and 266 µL isopropanol were added to each sample followed by incubation at room temperature for 10 min. Beads were washed 3 times with 70% Ethanol, then dried and eluted in 80 µL of water. 70 µL of RNA containing solution were added to 40 µL of Buffer RES and 7.5 µL of 100 mM NaIO_4_. The reaction was carried out on ice for 1 h and quenched with 10.5 µL of 100 mM DTT. Then 1.5 µL of Base Buffer and 16.5 µL DNAse I buffer (ThermoFisher) were added. This was followed by addition of 18 µL of Ambion Dnase I, RNase free (ThermoFisher) and incubation at 37 °C for 30 min. Finally, RNA was cleaned-up with Agencourt RNAClean XP beads (Beckman) following manufacturer’s guidelines. RNA was eluted in 25 µL of water. RNA concentration was measured with a Nanodrop2000 spectrophotometer and normalised across all replicates from the same selection.

Annealing was conducted with 6-12 µg of RNA, 1 µL of DNA probe (2 µM), 2.5 µL of 10x annealing buffer in a final volume of 25 µL. Annealing was conducted at 65°C for 5 min. Extension was conducted by adding 17 µL water, 5 µL NEBuffer 2.0 (NEB), 1 µL dNTPs without dCTP (10 mM), 1 µL Btn-11-dCTP (20 µM) (Jena Bioscience) and 1 µL Klenow fragment exo(-) (NEB) and incubating samples at 37 °C for 6 min. For each sample 10 µL of Dynabeads MyOne C1 streptavidin beads (Invitrogen) were washed twice with buffer W-T and resuspended in 50 µL of binding buffer. The beads were added to each sample and incubated with head-over-tail rotation at 4 °C for 1 h. Following binding, beads were washed three times with buffer W-T, twice with buffer B-T, once with buffer W-T and once with water. Each wash was performed with 200 µL of the respective buffer and samples were transferred to a fresh tube between each wash. Beads were then resuspended in 13 µL of RT-Hybridisation buffer and incubated at 65 °C for 5 min. 7 µL of RT-MM were added to each sample and reverse transcription was performed at 50 °C for 10 min. Then 1 µL of RNaseH (NEB) was added and samples were incubated at 37 °C for 20 min and 98 °C for 5 min to elute cDNA. cDNA was then recovered from the beads and diluted 1:2 with water.

Half of the recovered cDNA was used to clone stmRNA libraries for a second round of selection, as previously described^15^. PCRs obtained from replicates of selections with the same substrate were combined before Golden Gate assembly.

A second round of selection was then performed. In this case for each selection three replicate samples were grown in the presence of the substrate and three in the absence of the substrate. For each replicate, expression, RNA isolation and pulldown were performed as described above. Recovered cDNA was amplified by PCR using Q5 High-Fidelity 2x Master Mix (NEB), using oligos NGS_selections_1(1-8) and NGS_selections_2(1-8), containing Illumina adapters and indexes (Nextera XT-derived) for Illumina NGS. Libraries were purified using AMPureXP Beads (Beckman Coulter), pooled in equimolar amounts and sequenced using a NextSeq2000 system (Illumina) with P2 kits (600 cycle, XLEAP or SBS) + ∼20% PhiX spike-in.

### NGS Data analysis

The paired end reads obtained from NGS sequencing runs were first paired using PEAR^39^ and aligned to a PylRS reference sequence using Bowtie2^40^. In a second step, the targeted library positions were extracted, their sequence translated to a string of amino acids and occurrence of each variant counted using previously described R-scripts^15^. Subsequently, the frequency of each variant in the sequenced library as well as its enrichment (frequency in the selected vs the input sample) and selectivity score (frequency in the sample grown in the presence of ncM vs the absence of ncM) were calculated using previously described R-scripts^15^. Finally, empirical thresholds for enrichment, selectivity and normalized standard deviation values were determined and only variants passing these thresholds were selected for further testing.

### Fluoro-tREX characterisation of PylRS variants

A pMB1 plasmid encoding both tRNA^Pyl^ and a PylRS variant was transformed into chemically competent DH10B cells. Cells were rescued in 500 µL SOC for 10-30 min. Cells were diluted 1:50 in 2xYT-s (0.5-5 mL) and grown overnight. Overnight cultures were diluted into fresh 2xYT-s (1.5-5 mL) at a ratio between 1:20 and 1:50. Cells were grown to OD_600_ of 0.6-0.8. Cells were centrifuged at >3000 rcf at 4 °C for 10 min. Cells were resuspended in 600 µL buffer RES and centrifuged again as above. Cells were resuspended in 135 µL buffer RES and 15 µL of liquified phenol was added. Cells were lysed by head-over-tail rotation at room temperature for 20 min and were then spun at 4 °C, 4000 rcf for 20-30 min. The supernatant was added to 40 µL of chloroform and mixed thoroughly by pipetting. The mixture was centrifuged 4 °C, 4000 rcf for 10 min. 100 µL of the aqueous phase were transferred into a new vessel and 6 µL of 100 mM NaIO_4_ were added. The oxidation reaction was carried out on ice for 1 h. tRNAs were purified using the ZR-96 Oligo Clean & Concentrator kit from Zymo Research and eluted in 50 µL of water. 6 µL of 10x deacylation buffer was added to purified tRNAs and deacylation was carried out at 42 °C for 1 h. The reaction was quenched by adding 12 µL 3 M sodium acetate and purified using the ZR-96 Oligo Clean & Concentrator. tRNAs were eluted in 16 µL of water. RNA concentration was measured on a Nanodrop2000 spectrophotometer and normalised across all samples that were going to be compared.

0.5 µL 10x annealing buffer and 0.5 µL Cy3-labelled probe (2 µM) were added to 6 µL of RNA containing solution (1-5 µg). Hybridisation was carried out at 65 °C for 5 min. Then 3.1 µL water, 1.2 µL NEBuffer2.0, 0.25 µL dNTPs without dCTP (10 mM), 0.25 µL Btn11-dCTP (20 µM), 0.2 µL Klenow fragment exo(-) (NEB) were added and extension was carried out at 37 °C for 6 min. 12 µL of tREX loading dye were added. Samples were loaded onto a Novex TBE-Urea 15% PAGE gel (Invitrogen). Gel electrophoresis was run at 250 V for 1 h. Gels were imaged with an Amersham Typhoon Biomolecular Imager (GE) with Cy5 and Cy3 emission filters.

### Determining the identity of the acylating monomer by tRNA pulldown and LC-MS

A pMB1 plasmid encoding both tRNA^Pyl^ and a PylRS variant or a control only encoding tRNA^Pyl^ were transformed into chemically competent DH10B cells. Cells were rescued in 500 µL SOC for 10-30 min. Cells were diluted 1:50 in 2xYT-s (0.5-5 mL) and grown overnight. Overnight cultures were diluted into fresh 2xYT-s (5-10 mL) at a ratio between 1:20 and 1:50. Cells were grown to OD_600_ of 0.6-1.0. Cells were centrifuged at >3000 rcf at 4°C for 10 min. Cells were resuspended in 600 µL buffer D and centrifuged again as above. Cells were resuspended in 225 µL buffer D and 25 µL of liquified phenol was added. Cells were lysed by head-over-tail rotation at room temperature for 20 min and were then spun at 4 °C, 4000 rcf for 20-30 min. The supernatant was added to 250 µL of chloroform and mixed thoroughly by pipetting. The mixture was centrifuged 4 °C, 4000 rcf for 10 min. 200 µL of the aqueous phase were transferred into a new Eppendorf tube and RNA was purified with a Monarch RNA Cleanup Kit (50 µg) (NEB). RNA was eluted in 50 µL water into an Eppendorf tube containing 50 µL 2x Buffer D. Ninety microlitres of sample were hybridised with 0.5 µL of biotinylated DNA probe (100 µM) and annealing was carried out at 65 °C for 5 min. For each sample 40 µL of Dynabeads MyOne C1 streptavidin beads (Invitrogen) were washed twice with Buffer D +0.05% (v/v) Tween 20 and resuspended in 10 µL Buffer D. Beads were added to each sample and binding was carried out at 4°C with head-over-tail rotation for 1 h. Samples were washed with 3× acW1-t, 2× acW1, 3× acW2 and 1× water. All washes were carried out with 200 µL of the respective buffer. Elution was carried out with one of two protocols:

A) Base elution: 25 µL of deacylation buffer were added to the beads followed by incubation at 42 °C for 1 h. Twenty microlitres of supernatant were removed from the beads and 5 µL of 6-aminoquinolyl-*N*-hydroxysuccinimidyl carbamate (AQC, 3 mg ml^−1^ in acetonitrile) were added. The reaction was incubated at 55°C for 15 min. Samples were analysed either by single ion monitoring on an Agilent Technologies 6130 Quadrupole LC-MS or by a full-scan as described below.

B) RnaseA elution (adapted from ref^41^): 20 µL of RNaseA elution buffer were added to the beads and incubated at room temperature for 5 min. The supernatant was removed from the beads and 2.2 µL of 50% trichloroacetic acid (final concentration of 5%) were added. The mixture was cooled at-80 °C for 30 min, thawed and then spun at 21,130 rcf, 4°C, 30 min. The supernatant was transferred to a fresh vial and analysed as described below.

For full scans samples were analysed by LC-MS using a Vanquish HPLC (Thermoscientific, USA) with mobile phase A – 99.9% water with 0.1% formic acid and mobile phase B – 80% acetonitrile, 19.9% water, and 0.1% formic acid. Delivered flow was 150 μL min^-1^ and the gradient from 10% to 45% A over 17 minutes. Separation was carried out using a Kinetex 1.7 μm, EVO C18, 100 Å, 100 x 1.0 mm, LC column. Analytes were eluted and ionized with a HESI ion source onto an orbitrap mass spectrometry (Exploris MX, Thermoscientific USA). MS data were acquired in a broadband detection mode (R = 120,000, *m/z* 100 – 700) in positive ion mode (AGC = 200%, μscans = 5).

### Quantification of tRNA^Pyl^ acylation with ncMs and canonical amino acids

tRNA^Pyl^ isolation and pulldown was performed as described above using 2.5 mL cell culture. Triplicates were run for each sample. Controls were run in which tRNA^Pyl^ was expressed in the absence of PylRS and cells were grown in the presence of the relevant ncM/BocK. The concentration of isolated total RNA was normalised to 200 ng/µL and 90 µL of this were used for probe hybridisation and pulldown. Elution was carried out using protocol A for all samples and samples were diluted two-fold with deacylation buffer. For AQC derivatisation 10 µL of sample were treated with 2.5 µL of 5 mg/mL AQC in acetonitrile and incubated at 55 °C for 15 min. For dansyl chloride derivatization 10 µL of sample were treated with 2.5 µL of 10 mM dansyl chloride in acetonitrile and incubated at room temperature for 2 h.

Two sets of standards were used to generate standard curves. The first was generated by mixing all 20 natural amino acids each at the same concentration and making serial dilutions of the following concentration: 200 nM, 40 nM and 8 nM. The second set was generated by mixing **BocK**, **3**, **4**, **8**, **S3**, **S4**, Phe and Trp and making serial dilutions at the same concentrations as above. Derivatizations were carried out with 10 µL of standard following the same protocol as the samples. Triplicates were run for each standard curve.

LC-MS analysis was performed as described above and MS data was subsequently analysed using a custom python script based on the pyOpenMS interface^42^. In brief, for each relevant ncM and natural amino acid EICs were extracted for each sample. Integrated signals for each peak were generated and standard curves were used to calculate concentrations. The specific retention times for all ncMs and natural amino acids are given in **Supplementary Data 2**.

### tREX

This protocol was adapted from^15^. Total tRNA was extracted using the protocol described under “Fluoro-tREX characterisation of PylRS variants” from 4 mL cell culture grown in presence or absence of 4 mM of ncM, except for monomers **1** and **5** (2mM). The deacylated tRNA samples were normalised to 300 ng/μL and 6 μL thereof (1.8 μg RNA) were mixed with 0.5 μL 10x annealing buffer and 0.5 µL Cy5-labelled probe (2 µM). Primer hybridisation was carried out at 65 °C for 5 min. Then 3.35 µL water, 1.2 µL NEBuffer2.0, 0.25 µL dNTPs (10 mM), 0.1 µL Klenow fragment exo(-) (NEB) were added and extension was carried out at 37 °C for 6 min. Samples were mixed with 12 µL of tREX loading dye and loaded onto a 10% acrylamide, 1x TBE gel. Gel electrophoresis was performed at 180 V for 100 min. Gels were imaged using an Amersham Typhoon Biomolecular Imager (GE) with Cy5 emission filter.

### GFP Fluorescence

A pMB1 plasmid encoding both tRNA^Pyl^ and PylRS (or evolved variants thereof) and a p15A plasmid encoding a TAG containing sfGFP reporter were co-transformed into 20 μL chemically competent DH10B cells. Cells were rescued in 500 μL SOC at 37 °C for 1 h and 30 μL thereof were diluted into 470 μL 2xYT-s-ap and grown overnight. GFP expression was carried out in 96-well plates in presence and absence of 4 mM ncM by diluting 10 μL of the overnight culture into 500 μL 2xYT containing 0.2% L-arabinose. Plates were incubated at 37 °C, 750 rpm for 24 h and subsequently centrifuged at 4 °C, 4,000 rcf for 10 min. Cell pellets were resuspended in 150 μL PBS and 100 μL thereof were transferred to a Greiner 96-well flat bottom plate. OD_600_ and GFP fluorescence measurements were performed on a PHERAstar FS plate reader with focal height and gain adjustment set to 0. All values were normalized to expression with wt tRNA^Pyl^ and wt PylRS in the presence of 2 mM of **1**.

### GFP-His_6_ Purification and Mass spectrometry analysis

GFP-His_6_ expression was carried out as described above and samples were collected after 24 h. Three replicates of each protein sample were combined in a 1.5 mL Eppendorf tube and centrifuged at 4,000 rcf for 5 min. The cell pellet was resuspended in 500 μL lysis buffer and incubated at 4 °C for 1 h with head-over-tail rotation. The lysed samples were cleared by centrifugation at 4 °C, 20,000 rcf for 30 min and the cell lysate was incubated with 30 μL Ni-NTA beads for 1 h at 4 °C with head-over-tail rotation. The beads were first washed 3x with 500 μL 40 mM imidazole in PBS and then GFP-His_6_ was eluted 1-2x with 50 μL 300 mM imidazole in PBS. Samples were analysed by LC-MS using a Vanquish HPLC (Thermoscientific, USA) with mobile phase A – 90% water, 9.9% acetonitrile with 0.1% formic acid and mobile phase B – 80% acetonitrile, 19.9% water, and 0.1% formic acid. Delivered flow was 300 μL min^-1^ proteins were trapped on a MAbPac RP 2.1 x10 mm trapping column for two minutes before separation using a MAbPac RP 2.1 x 50 mm LC column. The gradient was from 10% to 30% B over 2 minutes before increasing to 55% B over the next 8 minutes at 80 °C. Proteins were eluted and ionized with a HESI ion source onto an orbitrap mass spectrometer (Exploris MX, Thermoscientific USA). MS data were acquired in a broadband detection mode (R = 120,000, *m/z* 600 – 2000, Intact protein mode) in positive ion mode (AGC = 100%, μscans = 10, in source energy = 10 eV). Protein deconvolution was carried out using a sliding window Xtract (isotopically resolved) deconvolution to generate monoisotopic protein masses.

### MS/MS analysis of purified GFP(150X)His_6_

Modified GFP protein samples were reduced with 5 mM DTT and alkylated with 20 mM iodoacetamide. SP3 protocol as described in Refs.^43,44^ was used to clean-up and buffer exchange the reduced and alkylated protein, shortly; proteins are washed with ethanol using magnetic beads for protein capture and binding. The proteins were resuspended in 100 mM NH_4_HCO_3_ and were digested with trypsin (Promega, UK) at an enzyme-to-substrate ratio of 1:50, and protease max 0.1% (Promega, UK). Digestion was carried out overnight at 37 °C. Clean-up of peptide digests was carried out with HyperSep SpinTip P-20 (ThermoScientific, USA) C18 columns, using 60% Acetonitrile as the elution solvent. Peptides were then evaporated to dryness *via* Speed Vac Plus (Savant, USA). Dried peptides were resuspended in 0.1% formic acid and run on a Vanquish Neo (ThermoSceintific, USA) to deliver a flow of 200 nL/min.

Peptides were trapped on a C18 Acclaim PepMap100 5 μm, 0.3 μm x 5 mm cartridge (ThermoScientific, USA) before separation on Aurora Ultimate C18, 1.7 μm, 75 μm x 25 cm (Ionopticks, Melbourne). Peptides were eluted on optimised gradients of 98 min and interfaced to a tribrid quadrupole Orbitrap mass spectrometer (Orbitrap Ascend, ThermoScientific, USA). MS data were acquired in data dependent mode with a Top-S method with 3 second cycle time, high resolution full mass scans were carried out (R = 120,000, *m/z* 400 – 1400) followed by higher energy collision dissociation (HCD) with stepped collision energy range 25, 30, 35 % normalised collision energy. The tandem mass spectra were recorded (R=60,000, isolation window *m/z* 1, dynamic exclusion 50 s, AGC 400%). Alternatively, peptides were analysed as describe in ‘Peptide macrocycle mass spectrometry’ below.

### Flow-cytometry-based analysis of sequence context libraries

Electrocompetent cells bearing a pMB1 plasmid encoding the target PylRS/tRNA^Pyl^ pair were transformed with the N-terminal GFP sequence context library (transformation efficiency >10^8^) and recovered in 2xTY-s-ap overnight. Cultures were diluted 1:50 in 3-5 mL of 2xYT-s-ap supplemented with 0.2% arabinose with and without 4 mM of the target ncM. The +ncM condition was set up in triplicate. Expression was conducted at 37 °C for 16-20 h with shaking at 220 rpm. 500 µL of culture was washed twice with 1 mL of PBS, resuspended in 1 mL of PBS and diluted 1:15-1:60 in PBS. All washing and dilution steps were conducted on ice. Cells were strained and sorted by FACS. Cell sorting was performed on a ThermoFisher Bigfoot Spectral cell sorter using a 70 µm nozzle at 60 psi. Cells were gated on Forwards Scatter Polar and Side Scatter Polar. Excitation was performed with a 488 nm laser set to 125 mW, emission was detected with a 549/15 bandpass filter, a 505 nm laser with a 455/14 bandpass filter was used as an empty channel. The sorting gate was set to include the top 0.5% of GFP fluorescence intensity from the library transformed into a negative control expressing only tRNA^Pyl^ and no PylRS. The same gate was applied to all samples. For each sample at least 20’000 cells were sorted into 500 µL of PBS and kept on ice until all samples in a batch were sorted. Cells were recovered by addition of 1 mL of 2xYT and incubation at 37 °C without shaking for 45 min and with shaking for 3 h. 3 mL of 2xYT supplemented with spectinomycin and apramycin at 1.5x working concentration were added and cultures grown overnight at 37 °C with shaking. Cultures were harvested and plasmids were extracted by miniprep (QIAprep Spin Miniprep Kit, Qiagen). Minipreps of the input cultures (harvested on the previous day) were also carried out.

### NGS and data analysis of sorted sequence context libraries

50 ng of prepped plasmid was used to set up a PCR reaction (50 µl scale, Q5 High-Fidelity 2x Master Mix (NEB), 200 µM of each primer) to amplify the sequence context library region and append Illumina adapters and indexes (Nextera XT-derived). Amplification was conducted for 18 cycles (98 °C for 10 s, 60 °C for 10 s, 72 °C for 30 s).). Amplicons were checked by gel electrophoresis and purified using AMPure XP (Beckman) according to manufacturer’s guidelines. Samples were pooled and PhiX control v3 (Illumina) was added at a molar ration of 15-30%. This library was sequenced at 0.8 nM on a NextSeq2000 system (Illumina) according to the manufacturer’s guidelines using the P2 XLEAP-SBS kit (100 Cycles).

Reads were paired with PEAR^45^ and aligned to a reference of the reporter library using Bowtie2^40^. Custom R-scripts^15^ were used for downstream analysis. To normalise for different sequencing depth of different samples each sequence in the library was normalised by the total counts in its respective sample to generate a normalised frequency. Sequences with mutations outside the target region, mutations to the target UAG codon as well as sequences containing an AUG codon following the target UAG codon were removed for downstream analysis. Frequencies were normalised by the percentage of cells that passed the gate in their respective sample. Only sequences present in all three replicates of the same sample (and in three replicates of the input) were considered for downstream analysis. The mean frequency of each sequence in the sorted sequence was divided by the mean in the input library to calculate an incorporation efficiency score. Sequences with a relative standard deviation greater than 1.6 in the sorted sample were filtered out. Heat maps were generated following the scheme in **Supplementary Fig. 24**.

For each sample an empirical threshold value for the NGS incorporation efficiency score was calculated. Sequences with a score above the threshold are likely to be permissive to ncM incorporation. The threshold was set as the lowest NGS score for a sequence for which 1) we measured an ncM-dependent increase in GFP fluorescence in the 58 experimentally validated clones (Fluorescence +ncM/ Fluorescence–ncM >1.2) and 2) for which the NGS score was above the highest NGS score for a clone for which we have directly measured background GFP signal independent of ncM addition. We then calculated the percentage of sequences in the library that had a score above the threshold.

### Selection of engineered tRNAs

Electrocompetent cells bearing a sfGFP(150TAG)-His_6_ reporter were transformed with a PylRS variant and a tRNA library (PylRS and the tRNA are encoded on the same plasmid) and recovered in 2xTY-s-ap overnight. Cultures were diluted 1:50 in 3-5 mL of 2xYT-s-ap supplemented with 0.2% arabinose with and without 4 mM of the target ncM. Expression was conducted at 37 °C for 16-20 h with shaking at 220 rpm. Samples were prepared and used in the same manner and with the same instrument settings for FACS as described in ‘FACS-based analysis of sequence context libraries’ (except for the selection with **3** for which we used a 525/35 bandpass filter). We set the gate to capture the top 0.5-2% of the fluorescent cell population, ensuring minimal overlap with the population of cells grown in the absence of ncM. Plasmids were extracted from input samples with a QIAprep Spin Miniprep Kit (Qiagen).

The output reporter plasmids were digested by treatment with DraI (NEB) and T5 exonuclease (NEB) following manufacturer guidelines. The sorted tRNA library was transformed into fresh electrocompetent cells with a sfGFP(150TAG)-His_6_ reporter. Recovery, expression and sorting were conducted in triplicate for each sample following the same procedure as in the first round. The gate for FACS was set to capture the top 0.5-1% fluorescent cells. A minimum of 10,000 cells were sorted per replicate. Recovery and plasmid prep were conducted in the same way as described for FACS-based analysis of sequence context libraries

### NGS and data analysis of tRNA selections

NGS sample preparation was carried out following the same procedure as describe in ‘NGS and data analysis of FACS sorted sequence context libraries. Specific primers containing Illumina adapters and indexes and amplifying the tRNA were used for PCR.

Paired-end reads were paired with PEAR and aligned to a tRNA^Pyl^ reference with Bowtie2. subsequent analysis was carried out with custom R-scripts. To normalise for different sequencing depth of different samples each sequence in the library was normalised by the total counts in its respective sample to generate a normalised frequency. Only sequences present in all three replicates of the sorted sample were considered for downstream analysis. Where no counts were detected in the input sample a placeholder value of 0.9999 counts was attributed. Sequences with a relative standard deviation above 0.5 in the three replicates of the sorted samples were excluded from analysis. For each tRNA an enrichment score was calculated by dividing the mean frequency in the sorted sample by the mean frequency in the input sample. We grouped tRNA variants by targeted mutations and consolidated mean frequencies of all variants with non-targeted mutations into the most common variant for further analysis. tRNAs were ranked by enrichment and the most enriched variants were cloned into pMB1 plasmids with the corresponding PylRS variant.

### Flow-cytometry-based screening of tRNA libraries

Electrocompetent cells bearing GFP reporters (encoding for s.c.1, s.c.2, s.c.3, s.c.4, s.c.50, or sfGFP(150TAG)-His_6_) were transformed with a PylRS variant and a tRNA library and recovered in 2xTY-s-ap overnight. Cultures were diluted 1:50 in 3-5 mL of 2xYT-s-ap supplemented with 0.2% arabinose with and without 4 mM of the target ncM. Expression was conducted at 37 °C for 16-20 h with shaking at 220 rpm. 500 µL of culture was washed twice with 1 mL of PBS, resuspended in 1 mL of PBS and diluted 1:15-1:600 in PBS. All washing and dilution steps were conducted on ice. Cells were analysed on a BD LSRFortessa. Cells were identified based on forward and side scatter relative to a buffer-only control. GFP fluorescence was measured using 488 nm excitation and a 530/25 nm bandpass filter.

### Parallel selection of engineered tRNAs on four sequence contexts

Electrocompetent cells bearing GFP reporters (encoding for s.c.1, s.c.2, s.c.3, s.c.4) were transformed with PylRS(β^3^-1) and tRNA libraries β^3^_2 or β^3^_3. Cells were sorted and enrichment of tRNA variants was analysed as described in the sections ‘Selection of engineered tRNAs’ and ‘NGS and data analysis of tRNA selections’. We then selected the ten most enriched tRNA variants from the β^3^_2 or β^3^_3 libraries enriched in selections on s.c.1, s.c.2, s.c.3, and s.c.4 reporters for characterisation. Additionally, we selected the 10 tRNA variants with the highest geometric mean of enrichment across 3 reporter selections for β^3^_2 or β^3^_3 libraries, and the 10 tRNA variants with the highest geometric mean of enrichment across all reporter selections for the β^3^_3 library.

We expressed all variants in the presence of PylRS(β^3^-1), **2** and GFP reporters s.c.1, s.c.2, s.c.3, s.c.4. We then chose the ten hits with the highest fluorescence averaged over all four reporters for further characterisation. We measured GFP fluorescence of cells grown with PylRS(β^3^-1) GFP reporters s.c.1, s.c.2, s.c.3, s.c.4, and sfGFP(150TAG)-His_6_ grown in the absence or presence of **2**. We identified β^3^_v2_tRNA^Pyl^ as the variant with the highest fluorescence in the presence and minimal fluorescence in the absence of **2** across reporters.

### Peptide macrocycle purification

Peptides were largely purified as described previously^5^. Briefly, 25 μL of chemically competent DH10B cells were transformed with a pMB1 plasmid encoding both tRNA^Pyl^ and PylRS (or evolved variants thereof) and a p15A plasmid encoding for a SUMO-GyrA-CBD(110TAG) fusion. Cells were rescued in 500 μL SOC at 37 °C for 1 h and 50 μL thereof were diluted into 450 μL 2xYT-s-ap and grown overnight. 250 μL of the overnight culture was diluted into 0.5-5 mL terrific broth (TB) containing 4 mM monomer. Flasks were incubated at 37 °C, 220 rpm for 3.5 h. L-arabinose (0.2% final concentration) was added to the culture, and the culture was incubated for 16 h at 37 °C, 220 rpm and subsequently centrifuged at 4 °C, 4,000 rcf for 10 min. Cell pellets were resuspended in 1 mL peptide lysis buffer. The resuspended pellet was lysed by sonication in an ice bath (2 mm tip, 10 × 10 s pulses every 10 s, 60% amplitude), or by the addition of 10% Bug buster and incubation at 4 °C for 1 h, and the cell lysate was cleared from debris by centrifugation (20,000 rcf, 45 min, 4 °C). The supernatant was transferred to a new tube and 125 μL Ni-NTA slurry were added. The suspension was incubated at 4 °C for 1 h under gentle agitation. The slurry was added to a Poly-Prep column (BioRad) and collected through vacuum filtration. Beads were washed twice with 20 mL NHI-30 and the sample was eluted with 500 μL NHI-250. The eluted protein was concentrated and buffer-exchanged on Amicon filters (3 kDa cut-off) to MOPS reaction buffer. Alternatively, cleared lysates were purified using Thermo Scientific™ HisPur™ Ni-NTA Spin Plates following manufactures guidelines, eluting with 400 μL NHI-250. Samples were then stored at 4 °C for further analysis.

For cyclization 20-300 μL protein elution was diluted in 100-180 μL MOPS reaction buffer and cyclisation was initiated through the addition of 0.005 mg ml^-1^ Ulp1, 100 mM DTT, and 2 mM MESNA. The reaction was incubated at 37 °C for 20 h. The resulting cyclic peptides were extracted through the addition of 10 μL 10% acetic acid and 100 μL 3:1 chloroform:isopropanol. The bottom phase was transferred to a new tube and dried under a continuous air flow at room temperature. The purified peptides were then resuspended in 25 μL 1% acetic acid.

### Peptide macrocycle mass spectrometry

Peptides were analysed by LC-MS/MS using a Vanquish Neo UHPLC (Thermo Fisher Scientific, USA) Orbitrap™ Excedion™ Pro mass spectrometer. A “Trap and Elute” system with a PepMap Neo trap column (300 µm x 5 mm, Thermo Scientific) and an Easy-Spray PepMap Neo, C18, 75 µm x 150 mm, 2 µm, 100 Å separation column was used at 350 nL/min^-1^ and 45 °C with the following gradient: 0 – 1 min 50% B;1– 8.5 min 50 - 90% B; 8.5 – 8.6 min 90 - 99% B and 8.6 – 14. min 99% B with a final run time of 14.5 min (A: H_2_O with 0.1 % formic acid, B: 80 % CH_3_OH with 0.1% formic acid).

The mass spectrometer was operated at positive polarity, and the ionization conditions were 280 °C for capillary temperature and 1900 V for spray voltage. Peptides were isolated using data dependent acquisition (top 20 scans from a full MS scan selected for fragmentation). MS scans were acquired within 400–1200 scan range (R = 120,000, AGC target 2 x e5). Isolation was followed by higher energy collision dissociation (HCD) with stepped collision (28, 30, 32% normalised collision energy) to acquire tandem mass spectra (R = 60,000, isolation window *m/z* 4, dynamic exclusion 10 s, AGC target 5 x e4).

## Statistics

GraphPad Prism version 10 was used to generate graphs. Sequence alignments and further processing of NGS data was caried out with custom R scripts.

## Acknowledgements

This work was supported by the Medical Research Council (MRC), UK (MC_U105181009 and MC_UP_A024_1008) to J.W.C. For the purpose of Open Access, the MRC Laboratory of Molecular Biology has applied a CC BY public copyright license to any Author Accepted Manuscript (AAM) version arising from this submission. A.D. and C.W. were supported by the Boehringer Ingelheim Fonds. D.R. was supported by a SNSF Postdoc.Mobility fellowship (grant number 235287) and an EMBO postdoctoral fellowship (ALTF 648-2025). We thank M. Spinck for advice on peptide macrocycle purification, J. McKenzie, S. Tapp, and P. Andree Penttilä for help with flow cytometry, and I. Liko, F. Goncalves de Almeida and C. Franco for help with mass spectrometry.

## Author contributions

J.W.C. supervised research. C.P., A.D., D.R., C.W., T.S.E., and Z.L. designed and performed experiments and data analysis. D.L.D. provided insights on tRNA display. T.M. contributed to mass spectrometry measurements. C.P., F.Z., and Y.L. designed FACS-based experiments.

K.C.L. advised on next-generation sequencing design and implemented HPC bioinformatics pipelines. C.P., A.D., D.R., C.W., T.S.E. and J.W.C. wrote the paper with input from all authors.

