## Supplementary Information for "Expanding the genetic code with diverse backbone structures across diverse sequence contexts"

**for**

### Supplementary Figures

**a**

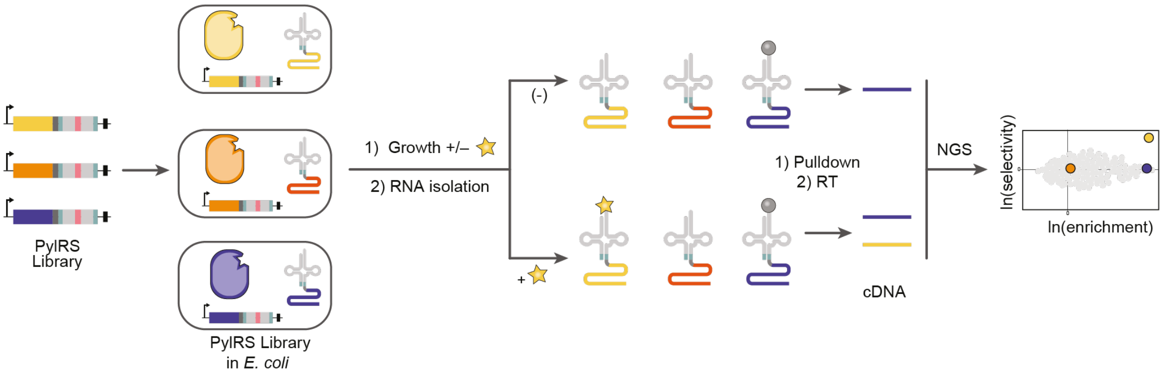


**Supplementary Figure 1. tRNA display enables discovery of PylRS variants that selectively acylate tRNA$^{Pyl}$ with ncMs**

**a,** Schematic of the tRNA display selection used to discover PylRS variants that selectively acylate their cognate tRNA with ncMs. *E. coli* cells are transformed with a library of genes encoding PylRS variants fused to a circularly permuted tRNA$^{Pyl}$. The cells produce a stmRNA from each gene. Each stmRNA contains a split version of tRNA$^{Pyl}$ fused to the mRNA for PylRS, which codes for the production of the corresponding PylRS protein. Active synthetases acylate the stmRNA with their substrates. Cells are grown in the presence and absence of the ncM of interest (yellow star). After RNA isolation, acylated tRNAs are selectively enriched using a biotinylation based pulldown (bio-mREX) and subsequently reverse transcribed (RT) to cDNA. cDNA is analysed by Next-generation-sequencing (NGS) and data is visualised as a spindle plot to identify synthetase variants that are active (x-axis, blue and yellow variants) and selective for the added ncM (y-axis, yellow variant).

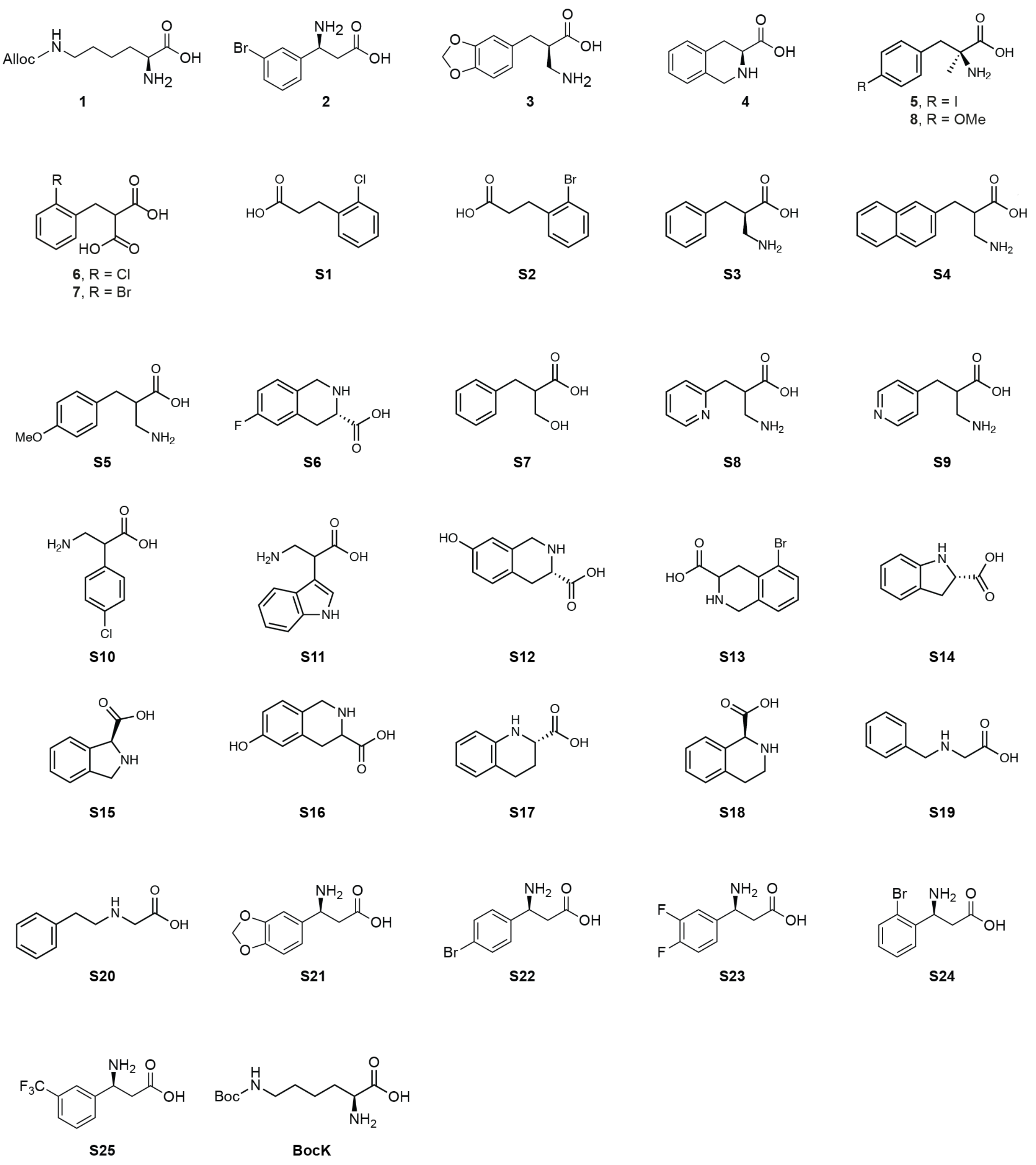


**Supplementary Figure 2. Full list of monomers used in this study.**

*N*[6]-(allyoxycarbony*l*)-*L*-lysine (**AllocK**, **1**), (*S*)-3-amino-3-(3-bromophenyl)propanoic acid (**2**), (*R*)-3-amino-2-(benzo[*d*][1,3]dioxol-5-ylmethyl)propanoic acid (**3**), (*S*)-1,2,3,4-tetrahydroisoquinoline-3-carboxylic acid (**4**), (*S*)-2-amino-3-(4-iodophenyl)-2-methylpropanoic acid (**5**), 2-(2-chlorobenzyl)malonic acid (**6**), 2-(2-bromobenzyl)malonic acid (**7**), *S*)-2-amino-3-(4-methoxyphenyl)-2-methylpropanoic acid (**8**), 3-(2-chlorophenyl)propanoic acid (**S1**), 3-(2-bromophenyl)propanoic acid (**S2**), 3-amino-2-benzylpropanoic acid (**S3**), 3-amino-2-(naphthalen-2-ylmethyl)propanoic acid (**S4**), 3-amino-2-(4-methoxybenzyl)propanoic acid (**S5**), 3-(((benzyloxy)carbonyl)amino)-2-(hydroxymethyl)propanoic acid (**S6**), 2-benzyl-3-hydroxypropanoic acid (**S7**), 3-amino-2-(pyridin-4-

ylmethyl)propanoic acid (**S8**), 3-amino-2-(pyridin-4-ylmethyl)propanoic acid (**S9**), (*S*)-indoline-2-carboxylic acid (**S10**), 2-(4-chlorophenyl)-3-hydroxypropanoic acid (**S10**), 3-amino-2-(4-chlorophenyl)propanoic acid (**S11**), (*S*)-7-hydroxy-1,2,3,4-tetrahydroisoquinoline-3-carboxylic acid (**S12**), 5-bromo-1,2,3,4-tetrahydroisoquinoline-3-carboxylic acid (**S13**), (*S*)-indoline-2-carboxylic acid (**S14**), (*S*)-isoindoline-1-carboxylic acid (**S15**), 6-hydroxy-1,2,3,4-tetrahydroisoquinoline-3-carboxylic acid (**S16**), (*S*)-1,2,3,4-tetrahydroquinoline-2-carboxylic acid (**S17**), (*S*)-1,2,3,4-tetrahydroisoquinoline-1-carboxylic acid (**S18**), Benzylglycine (**S19**), Phenethylglycine (**S20**), (*S*)-3-amino-3-(benzo[*d*][1,3]dioxol-5-yl)propanoic acid (**S21**), (*S*)-3-amino-3-(4-bromophenyl)propanoic acid (**S22**), *S*)-3-amino-3-(3,4-difluorophenyl)propanoic acid (**S23**), (*S*)-3-amino-3-(2-bromophenyl)propanoic acid (**S24**), (*S*)-3-amino-3-(3-(trifluoromethyl)phenyl)propanoic acid (**S25**), $N^6$-(*tert*-butoxycarbonyl)-*L*-lysine (BocK, **S26**)

**a**

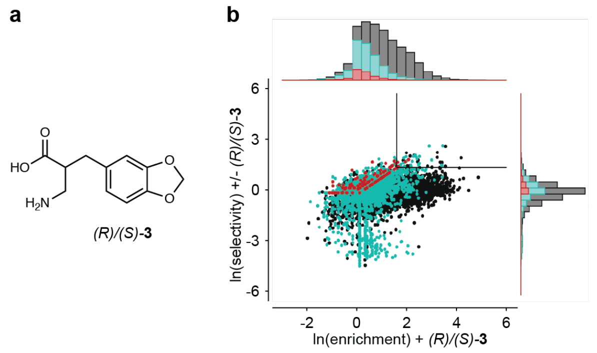


**c**

| Name | Ranking | 300 | 302 | 305 | 306 | 309 | 344 | 346 | 348 | 401 | 6 Selectivity | 6 Enrichment |
|---|---|---|---|---|---|---|---|---|---|---|---|---|
| | | *Mm*PylRS residue | | | | | | | | | | |
| wt | | M | A | L | Y | L | M | N | C | V | | |
| 21.24 | 1 | C | H | L | Y | L | M | A | S | L | 18.23 | 31.41 |
| 21.25 | 2 | C | H | N | Y | N | M | A | S | V | 18.05 | 18.38 |
| 21.26 | 3 | D | H | L | Y | N | M | A | S | A | 17.63 | 35.24 |
| 21.27 | 4 | D | H | L | Y | F | M | A | C | V | 12.20 | 25.30 |
| | 5 | C | G | C | C | G | M | V | S | V | 11.95 | 7.33 |
| 21.28 | 6 | D | H | L | D | L | M | A | L | V | 10.28 | 10.47 |
| 21.29 | 7 | D | H | L | N | L | M | T | F | V | 10.22 | 10.04 |
| | 8 | M | A | L | Y | L | Q | C | F | V | 9.85 | 8.51 |
| | 9 | M | H | V | L | L | M | T | S | V | 9.61 | 9.76 |
| 21.33 | 10 | D | H | C | Y | L | M | A | G | V | 9.57 | 12.39 |
| 21.31 | 11 | S | C | F | F | L | M | A | G | V | 9.49 | 7.26 |
| | 12 | A | G | R | G | G | M | G | S | C | 9.42 | 7.19 |
| | 13 | M | H | L | C | L | Q | C | L | K | 9.24 | 10.89 |
| 21.34 | 14 | D | H | L | L | L | M | A | S | C | 9.06 | 25.16 |
| | 15 | C | H | I | Y | N | M | A | N | V | 8.81 | 65.73 |
| 21.35 | 16 | C | H | L | N | L | M | A | G | V | 8.76 | 6.61 |
| | 17 | D | H | L | I | L | M | G | S | V | 8.22 | 11.21 |
| | 18 | C | D | H | F | L | M | E | C | F | 7.87 | 8.02 |
| | 19 | A | H | H | Y | C | M | S | V | V | 7.71 | 11.78 |
| | 20 | C | H | L | Y | C | M | A | N | C | 7.71 | 20.16 |
| | 21 | D | H | L | Y | L | M | A | N | S | 7.55 | 6.50 |
| | 22 | C | H | L | Y | F | M | A | L | V | 7.43 | 11.67 |
| | 23 | D | H | V | Y | F | M | A | G | V | 7.35 | 7.45 |
| 21.30 | 24 | D | C | C | F | V | M | A | I | T | 7.15 | 9.26 |
| | 25 | D | H | L | Y | I | M | G | V | C | 7.01 | 11.39 |
| | 26 | D | H | L | F | G | M | G | L | C | 6.94 | 7.06 |
| | 27 | C | H | L | Y | L | M | A | G | C | 6.75 | 28.52 |
| | 28 | D | C | C | F | C | M | G | I | V | 6.75 | 5.15 |
| | 29 | D | H | R | C | H | M | N | Y | C | 6.70 | 5.18 |
| | 30 | M | H | C | Y | L | Q | A | C | V | 6.70 | 30.49 |
| | 31 | M | H | L | Y | C | M | A | F | K | 6.68 | 6.80 |
| | 32 | C | H | H | H | H | Q | V | G | C | 6.61 | 6.73 |
| | 33 | M | H | I | Y | C | M | A | C | V | 6.51 | 6.60 |
| | 34 | D | H | L | N | L | M | V | F | V | 6.28 | 12.32 |
| | 35 | D | H | I | Y | V | M | A | C | T | 6.12 | 30.06 |
| 21.32 | 36 | S | C | C | F | V | M | A | V | V | 6.03 | 5.93 |

**d**

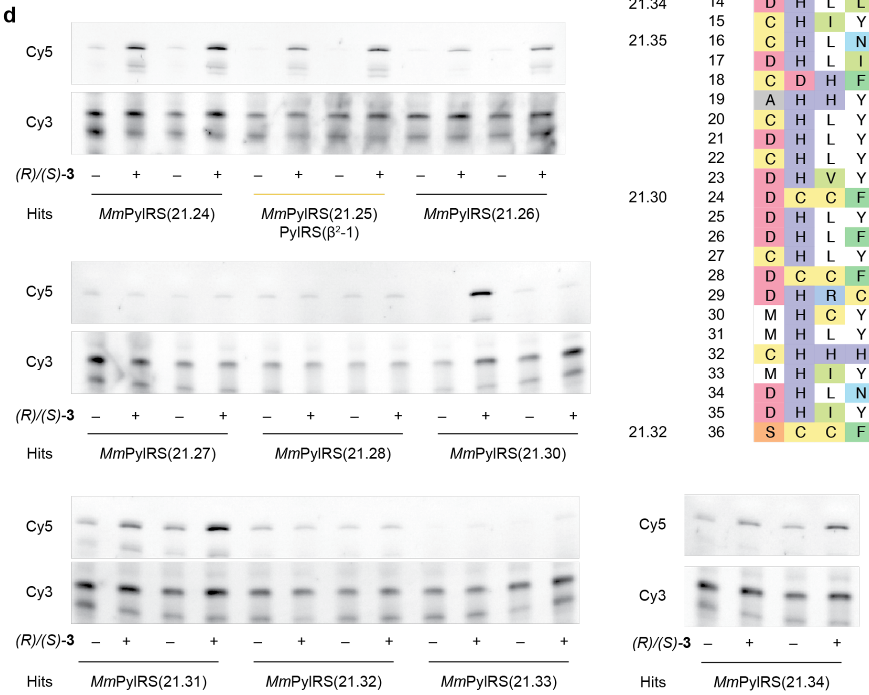


**Supplementary Figure 3. Selection of PylRS variants for 3 by tRNA display.**

**a**, Chemical structure of compound **3**. **b**, Spindle plot obtained from selection of PylRS with compound **3**. The X-axis shows the logarithm of enrichment, calculated from the ratio of the abundances of a given sequence in the positive sample (+**3**) versus the input library. The y-axis shows the logarithm of selectivity, calculated from the ratio of the abundances of a given sequence in the positive sample versus the negative sample (-**3**). Data points are divided into three categories indicated by colour. Black data points represent sequences observed in all replicates of the positive and negative sample. Blue data points are sequences observed in all replicates of the positive sample and at least one replicate of the negative sample. Red data points represent sequences observed in all replicates of the positive sample and not observed in any replicate of the negative sample. Marginal histograms depict the number of data points for a bin along each axis. Histogram bin size was determined using the Freedman-Diaconis Rule. The number of data points of each category is recorded in each histogram with the category's corresponding colour. The lines in the upper right quadrant of the plot delineate the region of the spindle plot where selectivity and enrichment is ≥ 5 and selectivity is ≥5. The plot was run with an error threshold of 0.5. **c**, Sequence of hits found in the delineated region of the spindle plot. Calculated selectivity and enrichment are shown for each hit as well as its ranking by selectivity amongst all gated hits. Hits that were chosen for cloning and testing are given a name with which they are described throughout the text and in subsequent panels and figures. **d**, ftREX of PylRS variants selected in the presence of **3**. Experiments were performed using tRNA extracted from cells harbouring a pMB1 plasmid encoding each PylRS and $tRNA^{Pyl}$ in presence and absence of **3** (4 mM). Experiments were performed in duplicate.

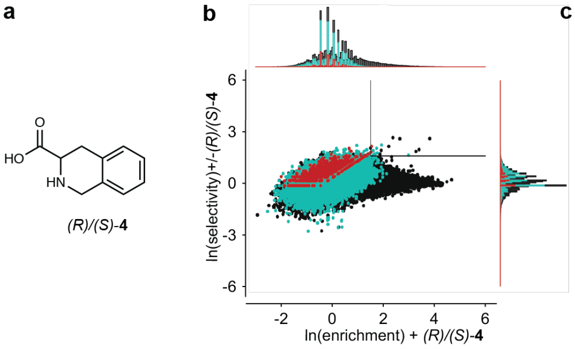


c

| Name | Ranking | 300 | 302 | 305 | 306 | 309 | 344 | 346 | 348 | 401 | 10 Selectivity | 10 Enrichment |
|---|---|---|---|---|---|---|---|---|---|---|---|---|
| | | *Mm*PylRS residue | | | | | | | | | | |
| wt | | M | A | L | Y | L | M | N | C | V | | |
| 18.11 | 1 | D | S | L | F | L | M | C | C | F | 14.31 | 10.76 |
| 18.12 | 2 | S | G | L | Y | G | M | A | V | V | 13.61 | 45.37 |
| 18.13 | 3 | S | G | L | Y | G | M | S | V | V | 13.48 | 14.16 |
| 18.14 | 4 | S | G | C | Y | G | M | C | V | C | 9.39 | 6.90 |
| 18.15 | 5 | D | S | V | F | C | M | C | I | F | 8.66 | 4.66 |
| 18.16 | 6 | D | G | L | G | L | M | A | V | F | 8.64 | 9.19 |
| | 7 | D | Y | L | C | C | M | V | F | C | 7.74 | 5.69 |
| 18.17 | 8 | D | G | F | Y | G | M | S | V | V | 7.21 | 30.00 |
| 18.18 | 9 | S | G | I | Y | S | M | C | L | V | 6.88 | 6.82 |
| | 10 | N | G | Y | L | F | M | C | N | C | 6.76 | 4.97 |
| | 11 | M | A | Q | Y | L | M | A | L | V | 6.55 | 4.82 |
| 18.19 | 12 | S | G | F | Y | G | M | S | V | V | 6.43 | 26.53 |
| | 13 | M | S | N | G | V | M | N | R | L | 6.23 | 4.58 |
| 18.20 | 14 | D | G | H | Y | G | M | S | V | V | 5.92 | 5.86 |
| | 15 | D | H | C | Y | V | M | C | V | C | 5.87 | 5.82 |
| | 16 | M | H | I | H | L | M | T | G | V | 5.60 | 5.37 |
| | 17 | D | H | F | Y | F | M | C | C | C | 5.49 | 6.49 |
| | 18 | M | H | L | I | L | Q | N | F | C | 5.34 | 5.12 |
| 18.21 | 19 | S | G | G | Y | L | M | C | Y | V | 5.22 | 19.85 |

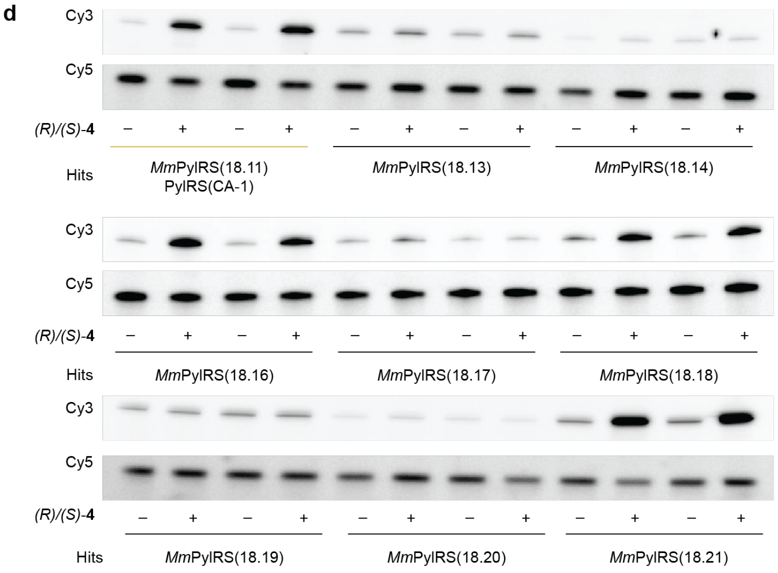


**Supplementary Figure 4. Selection of PylRS variants for** 4 **by tRNA display.**

**a**, Chemical structure of compound **4**. **b**, Spindle plot obtained from selection of PylRS with compound **4**. The x-axis shows the logarithm of enrichment, calculated from the ratio of the abundances of a given sequence in the positive sample (+**4**) versus the input library. The y-axis shows the logarithm of selectivity, calculated from the ratio of the abundances of a given sequence in the positive sample versus the negative sample (-**4**). Data points are divided into three categories indicated by colour. Black data points represent sequences observed in all replicates of the positive and negative sample. Blue data points are sequences observed in all replicates of the positive sample and at least one replicate of the negative sample. Red data points represent sequences observed in all replicates of the positive sample and not observed in any replicate of the negative sample. Marginal histograms depict the number of data points for a bin along each axis. Histogram bin size was determined using the Freedman-Diaconis Rule. The number of data points of each category is recorded in each histogram with the category's corresponding colour. The lines in the upper right quadrant of the plot delineate the region of the spindle plot where selectivity and enrichment is ≥ 4.5 and selectivity is ≥ 5. The plot was run with an error threshold of 1. **c**, Sequence of hits found in the delineated region of the spindle plot. Calculated selectivity and enrichment are shown for each hit as well as its ranking by selectivity amongst all gated hits. Hits that were chosen for cloning and testing are given a name with which they are described throughout the text and in subsequent panels and figures. **d**, ftREX of PylRS variants selected in the presence of **4**. Experiments were performed using tRNA extracted from cells harbouring a pMB1 plasmid encoding each PylRS and $tRNA^{Pyl}$ in presence and absence of **4** (4 mM). Experiments were performed in duplicates.

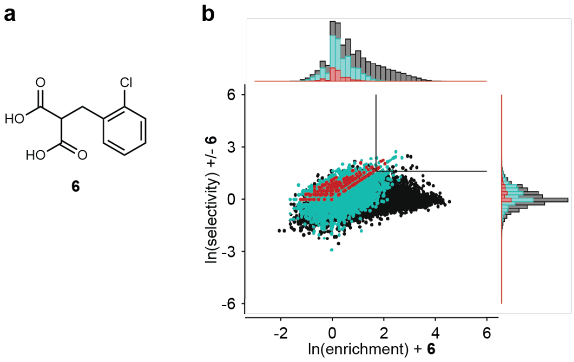


c

| Name | Ranking | 300 | 302 | 305 | 306 | 309 | 344 | 346 | 348 | 401 | 2 Selectivity | 2 Enrichment |
|---|---|---|---|---|---|---|---|---|---|---|---|---|
| | | *Mm*PylRS residue | | | | | | | | | | |
| wt | | M | A | L | Y | L | M | N | C | V | | |
| 22.38 | 1 | S | H | L | Y | L | Q | A | S | L | 15.94 | 11.85 |
| 22.39 | 2 | N | H | L | Y | L | M | G | L | L | 12.42 | 18.75 |
| 22.40 | 3 | S | H | L | Y | I | Q | G | C | V | 11.91 | 21.30 |
| 22.41 | 4 | S | S | L | R | L | Q | N | H | A | 11.84 | 11.79 |
| 22.42 | 5 | S | G | L | Y | L | M | V | H | V | 11.29 | 15.05 |
| 22.43 | 6 | N | H | L | Y | L | Q | G | V | V | 11.29 | 10.96 |
| | 7 | E | R | N | F | C | Q | C | V | L | 11.11 | 5.62 |
| | 8 | N | H | L | Y | L | Q | A | I | L | 10.57 | 10.53 |
| 22.44 | 9 | S | S | H | Y | S | M | A | R | V | 9.74 | 7.33 |
| | 10 | E | S | G | G | V | M | V | N | S | 9.64 | 9.60 |
| | 11 | N | H | C | Y | V | M | A | L | V | 9.39 | 6.98 |
| 22.45 | 12 | N | H | L | Y | V | M | C | I | V | 9.32 | 9.29 |
| | 13 | S | S | L | R | L | Q | N | H | V | 9.09 | 6.87 |
| 22.46 | 14 | S | H | N | Y | L | Q | A | F | L | 9.24 | 9.05 |
| | 15 | N | H | H | Y | L | M | V | I | L | 9.05 | 6.73 |
| 22.47 | 16 | S | S | S | R | L | M | V | H | V | 8.97 | 8.94 |
| 22.48 | 17 | S | H | S | Y | L | Q | C | I | L | 8.62 | 8.59 |

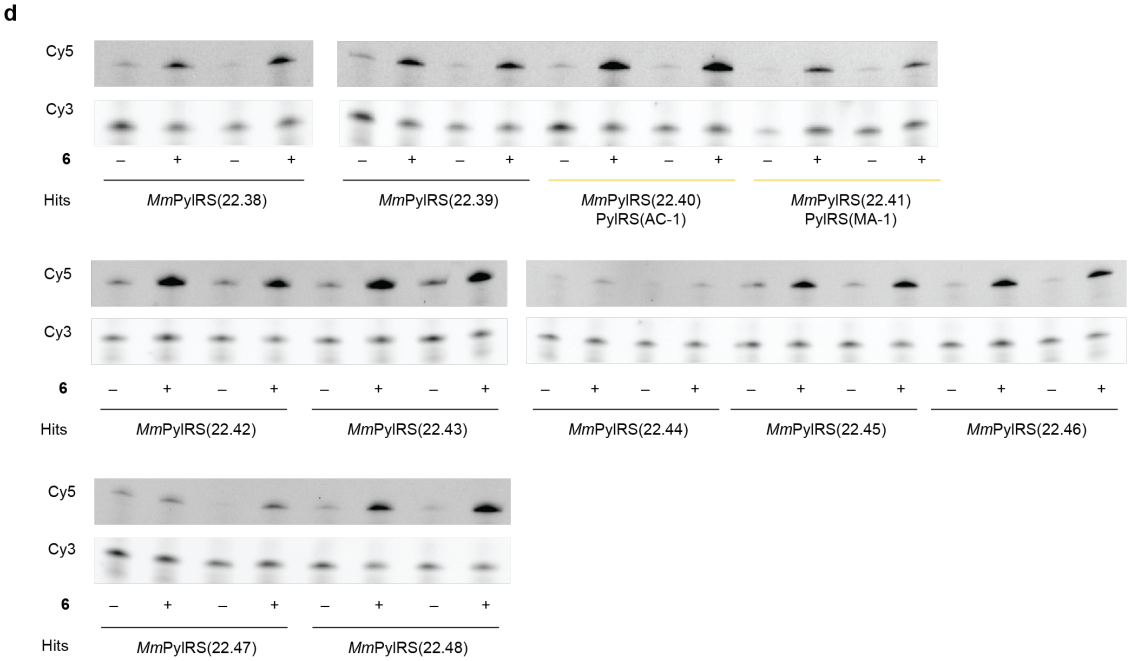


**Supplementary Figure 5. Selection of PylRS variants for** 6 **by tRNA display.**

**a**, Chemical structure of compound **6**. **b**, Spindle plot obtained from selection of PylRS with compound **2**. The x-axis shows the logarithm of enrichment, calculated from the ratio of the abundances of a given sequence in the positive sample (+**6**) versus the input library. The y-axis shows the logarithm of selectivity, calculated from the ratio of the abundance of a given sequence in the positive versus the negative condition (-**6**). Data points are divided into three categories indicated by colour. Black data points represent sequences observed in all replicates of the positive and negative samples. Blue data points are sequences observed in all replicates of the positive sample and at least one replicate of the negative sample. Red data points represent sequences observed in all replicates of the positive sample and not observed in any replicate of the negative sample. Marginal histograms depict the number of data points for a bin along each axis. Histogram bin size was determined using the Freedman-Diaconis Rule. The number of data points of each category is recorded in each histogram with the category's corresponding colour. The lines in the upper right quadrant of the plot delineate the region of the spindle plot where selectivity and enrichment is ≥ 5.5 and selectivity is ≥ 5. The plot was run with an error threshold of 0.7. **c**, Sequence of hits found in the delineated region of the spindle plot. Calculated selectivity and enrichment are shown for each hit as well as its ranking by selectivity amongst all gated hits. Hits that were chosen for cloning and testing are given a name with which they are described throughout the text and in subsequent panels and figures. **d**, ftREX of PylRS variants selected in the presence of **6**. Experiments were performed using tRNA extracted from cells harbouring a pMB1 plasmid encoding each PylRS and $tRNA^{Pyl}$ in presence and absence of **6** (4 mM). Experiments were performed in duplicate.

**a**

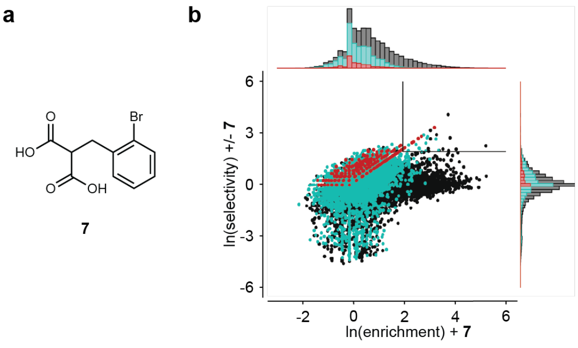


**c**

| Name | Ranking | *Mm*PylRS residue 300 | 302 | 305 | 306 | 309 | 344 | 346 | 348 | 401 | 3 Selectivity | Enrichment |
|---|---|---|---|---|---|---|---|---|---|---|---|---|
| wt | | M | A | L | Y | L | M | N | C | V | | |
| 20.01 | 1 | S | S | S | R | L | Q | V | Y | V | 59.34 | 41.75 |
| 20.02 | 2 | S | D | L | R | N | Q | G | I | V | 27.92 | 24.38 |
| 20.03 | 3 | S | H | L | R | L | M | S | V | K | 23.8 | 20.78 |
| 20.04 | 4 | N | H | V | Y | L | M | C | I | V | 18.45 | 26.51 |
| 20.05 | 5 | S | N | L | Y | L | Q | G | C | V | 18.25 | 15.94 |
| 20.06 | 6 | S | S | S | R | L | M | C | Y | K | 16.70 | 14.59 |
| 20.07 | 7 | N | H | L | Y | V | M | A | V | V | 16.51 | 18.60 |
| 20.08 | 8 | S | H | S | Y | L | Q | G | F | V | 15.40 | 45.13 |
| 20.09 | 9 | N | H | L | Y | V | M | G | L | V | 15.30 | 18.12 |
| | 10 | N | H | L | Y | L | Q | G | C | C | 14.77 | 12.9 |
| | 11 | S | H | L | Y | L | Q | G | I | L | 14.50 | 20.83 |
| | 12 | N | H | L | Y | L | Q | G | V | L | 14.32 | 25.82 |
| | 13 | S | S | S | R | L | Q | N | H | C | 12.21 | 10.66 |
| | 13 | S | H | V | Y | L | Q | A | I | V | 12.20 | 10.66 |
| 20.10 | 14 | S | S | S | R | L | M | A | H | V | 11.57 | 10.10 |
| | 15 | S | H | L | Y | N | Q | G | V | V | 11.31 | 17.42 |
| | 16 | A | H | I | Y | L | Q | G | I | V | 10.48 | 11.80 |
| | 17 | D | S | L | Y | L | M | E | V | V | 10.11 | 8.83 |
| 20.11 | 18 | N | H | N | Y | L | Q | A | F | V | 9.70 | 183.34 |

**d**

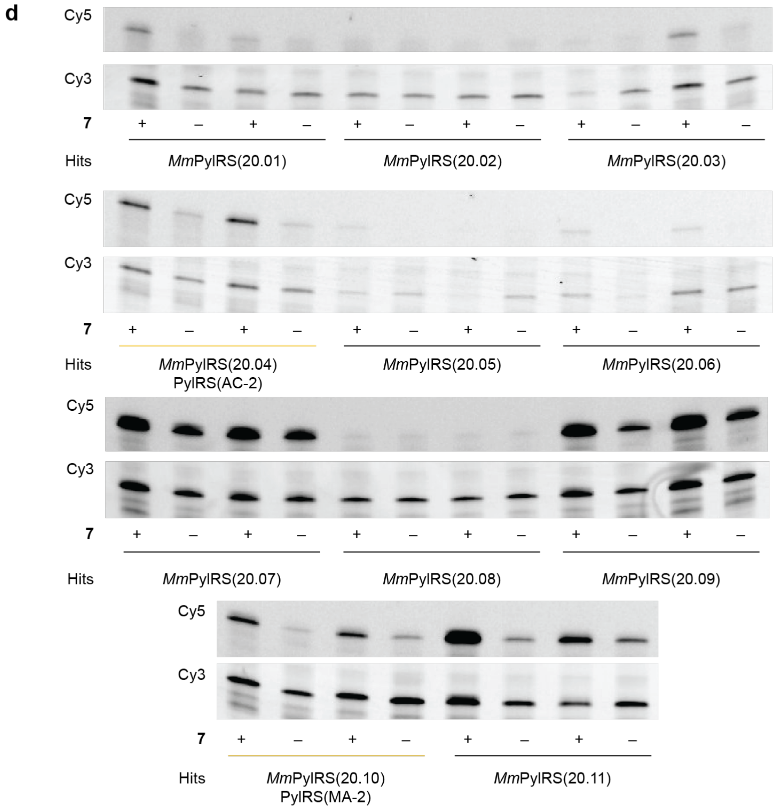


**Supplementary Figure 6. Selection of PylRS variants for 7 by tRNA display.**

**a**, Chemical structure of compound **7**. **b**, Spindle plot obtained from selection of PylRS with compound **3**. The x-axis shows the logarithm of enrichment, calculated from the ratio of the abundances of a given sequence in the positive sample (+**7**) versus the input library. The y-axis shows the logarithm of selectivity, calculated from the ratio of the abundances of a given sequence in the positive sample versus the negative sample (-**7**). Data points are divided into three categories indicated by colour. Black data points represent sequences observed in all replicates of the positive and negative sample. Blue data points are sequences observed in all replicates of the positive sample and at least one replicate of the negative sample. Red data points represent sequences observed in all replicates of the positive sample and not observed in any replicate of the negative sample. Marginal histograms depict the number of data points for a bin along each axis. Histogram bin size was determined using the Freedman-Diaconis Rule. The number of data points of each category is recorded in each histogram with the category's corresponding colour. The lines in the upper right quadrant of the plot delineate the region of the spindle plot where selectivity and enrichment is ≥ 7 and selectivity is ≥ 7. The plot was run with an error threshold of 0.5. **c**, Sequence of hits found in the delineated region of the spindle plot. Calculated selectivity and enrichment are shown for each hit as well as its ranking by selectivity amongst all gated hits. Hits that were chosen for cloning and testing are given are name with which they are described throughout the text and subsequent panels and figures. **d**, ftREX of PylRS variants selected in the presence of **7**. Experiments were performed using tRNA extracted from cells harbouring a pMB1 plasmid encoding each PylRS and $tRNA^{Pyl}$ in presence and absence of **7** (4 mM). Experiments were performed in duplicate.

**a**

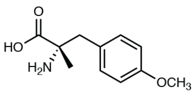


**b**

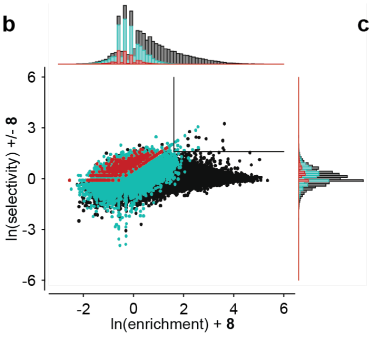


**c**

| | | *Mm*PylRS residue | | | | | | | | | 1 | |
|---|---|---|---|---|---|---|---|---|---|---|---|---|
| Name | Ranking | 300 | 302 | 305 | 306 | 309 | 344 | 346 | 348 | 401 | Selectivity | Enrichment |
| wt | | M | A | L | Y | L | M | N | C | V | | |
| **20.35** | **1** | **D** | **H** | **C** | **F** | **F** | **M** | **A** | **G** | **V** | **25.91** | **37.50** |
| 20.36 | 2 | D | G | C | F | F | M | V | C | L | 21.58 | 13.13 |
| 20.37 | 3 | D | G | C | F | F | M | V | S | V | 17.67 | 11.99 |
| | 4 | D | G | C | F | L | M | A | S | L | 16.18 | 9.37 |
| 20.39 | 5 | D | G | F | F | F | M | V | S | V | 12.83 | 7.81 |
| 20.40 | 6 | D | H | C | Y | F | M | C | G | L | 11.73 | 12.12 |
| | 7 | D | G | C | F | F | M | A | G | L | 11.51 | 13.64 |
| | 8 | D | G | C | Y | F | M | A | G | L | 10.72 | 8.80 |
| | 9 | D | G | I | F | F | M | V | G | L | 10.71 | 18.15 |
| | 10 | D | G | I | F | F | M | V | G | V | 9.62 | 5.85 |
| 20.41 | 11 | D | H | C | F | F | M | A | G | C | 9.58 | 35.15 |

**d**

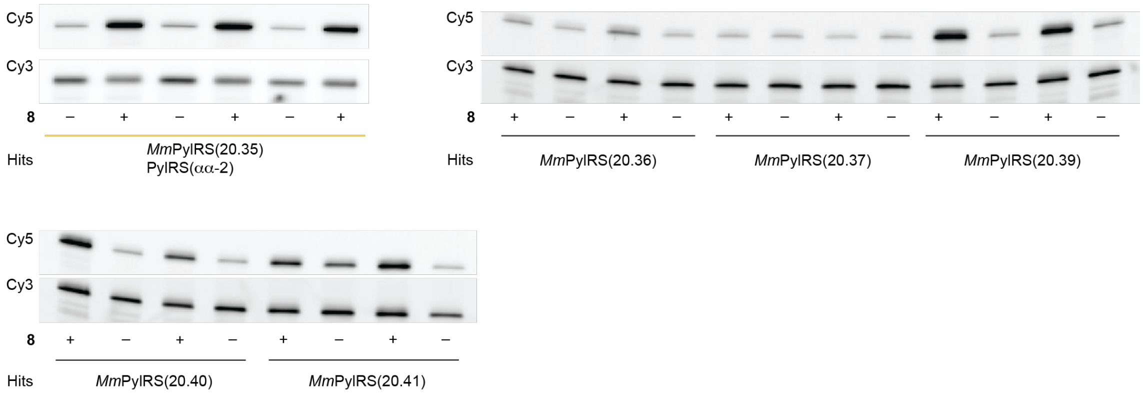


**Supplementary Figure 7. Selection of PylRS variants for** 8 **by tRNA display.**

**a**, Chemical structure of compound **8**. **b**, Spindle plot obtained from selection of PylRS with compound **1**. The x-axis shows the logarithm of enrichment, calculated from the ratio of the abundances of a given sequence in the positive samples (+**8**) versus the input library. The y-axis shows the logarithm of selectivity, calculated from the ratio of the abundances of a given sequence in the positive samples versus the negative samples (-**8**). Data points are divided into three categories indicated by colour. Black data points represent sequences observed in all replicates of the positive and negative samples. Blue data points are sequences observed in all replicates of the positive sample and at least one replicate of the negative sample. Red data points represent sequences observed in all replicates of the positive sample and not observed in any replicate of the negative sample. Marginal histograms depict the number of data points for a bin along each axis. Histogram bin size was determined using the Freedman-Diaconis Rule. The number of data points of each category is recorded in each histogram with the category's corresponding colour. The lines in the upper right quadrant of the plot delineate the region of the spindle plot where selectivity and enrichment are ≥ 5. The plot was run with an error threshold of 0.8. **c**, Sequence of hits found in the delineated region of the spindle plot. Calculated selectivity and enrichment are shown for each hit as well as its ranking by selectivity amongst all gated hits. Hits that were chosen for cloning and testing are given a name with which they are described throughout the text and in subsequent panels and figures. **d**, ftREX of PylRS variants selected in the presence of **8**. Experiments were performed using tRNA extracted from cells harbouring a pMB1 plasmid encoding each PylRS and $tRNA^{Pyl}$ in presence and absence of 4 mM of **8**. Experiments were performed in duplicate for all hits and in triplicate for PylRS(20.35).

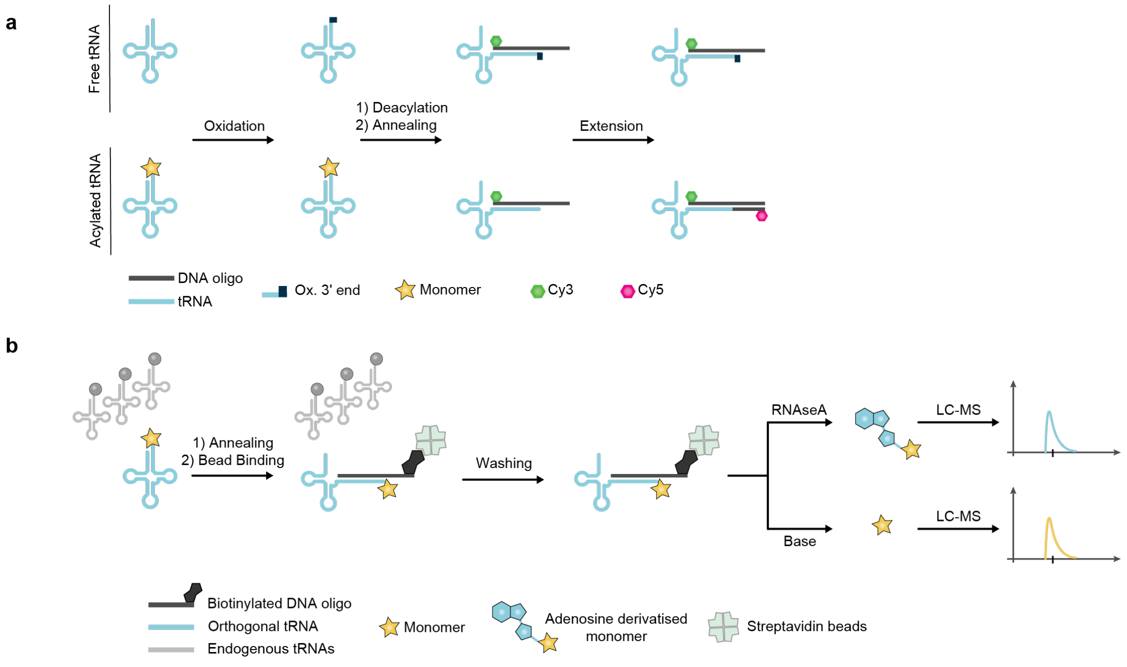


**Supplementary Figure 8. Determining acylation status by ftREX and mass spectrometry.**

**a**, Determining the acylation status of a target tRNA by ftREX. tRNAs are isolated from *E. coli* and treated with $NaIO_4$. Free tRNAs are oxidised by $NaIO_4$, which cleaves the vicinal diol of the 3' adenosine ribose ring. In contrast acylated tRNAs are protected from oxidation. tRNAs are then deacylated by base treatment and a Cy3 labelled DNA oligo is annealed to the 3' end of the target tRNA. Klenow fragment exo(-) is added to extend previously acylated target tRNAs, while the 3' of oxidised tRNAs cannot be extended. Cy5 labelled nucleotides are added to the extension reaction. Following polyacrylamide gel electrophoresis the Cy5 signal is visualized and reports on the extent to which the target tRNA was acylated. The Cy3 signal from the DNA probe acts as a loading control for the total amount of target tRNA.

**b**, Schematic of the tRNA pulldown assay used to identify monomers acylated onto target tRNAs. First, total tRNA is isolated from *E. coli*. Then, a biotinylated DNA oligo is annealed to a target tRNA. This allows selective binding to streptavidin beads. Other tRNAs and residual amino acids are then washed off. Two methods can be used to elute the target monomer. The first relies on treatment with RNaseA which releases the adenosine derivatised monomer. The second relies on treatment with base to elute the monomer which is then derivatised with AQC (6-aminoquinolyl-*N*-hydroxysuccinimidyl carbamate). Derivatised monomers are analysed by liquid chromatography-mass spectrometry (LC-MS).

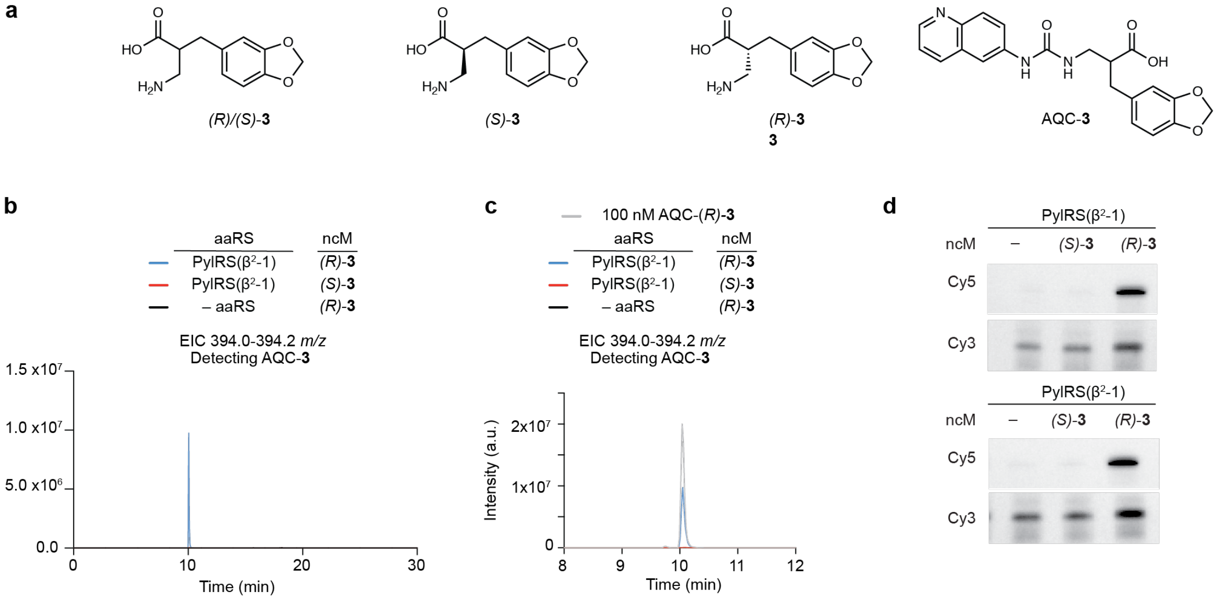


**Supplementary Figure 9. PylRS(β$^2$-1) acylates tRNA$^{Pyl}$ with the *S*-enantiomer of** 3**.**

**a**, Chemical structures of racemic compound **3**, its enantiomerically pure forms *S*-**3** and *R*-**3** and its AQC derivative AQC-**3**. **b**, Extracted Ion Chromatogram (EIC) for AQC-**3**. RNA$^{Pyl}$ was expressed with or without the synthetase PylRS(β$^2$-1) and cells were grown in the presence of *(R)*-**3** (blue traces) or (*S)*-**3** (red traces) (4 mM). A pulldown of tRNA$^{Pyl}$ was performed and the acylated ncM was eluted by treatment with base and AQC derivatised and analysed by LC-MS (M+H 394.0-394 *m/z*). Black traces show samples in which tRNA$^{Pyl}$ was expressed in the absence of PylRS. Maroon traces show samples in which tRNA$^{Pyl}$ was expressed in the presence of PylRS(β$^2$-1). **c**, Zoom in on (**b**). A standard containing 100 nM of *(R)*-**3** (grey trace) was derivatised in parallel to the samples and analysed by LC-MS. It is shown for reference (grey trace). **d**, ftREX of PylRS(β$^2$-1) with the enantiomers of compound **3**. Experiments were performed using tRNA extracted from cells harbouring a pMB1 plasmid encoding tRNA$^{Pyl}$ and PylRS(β$^2$-1). Cells were grown in absence of any ncM or in the presence of the shown monomers (4 mM). The experiment was performed in duplicates.

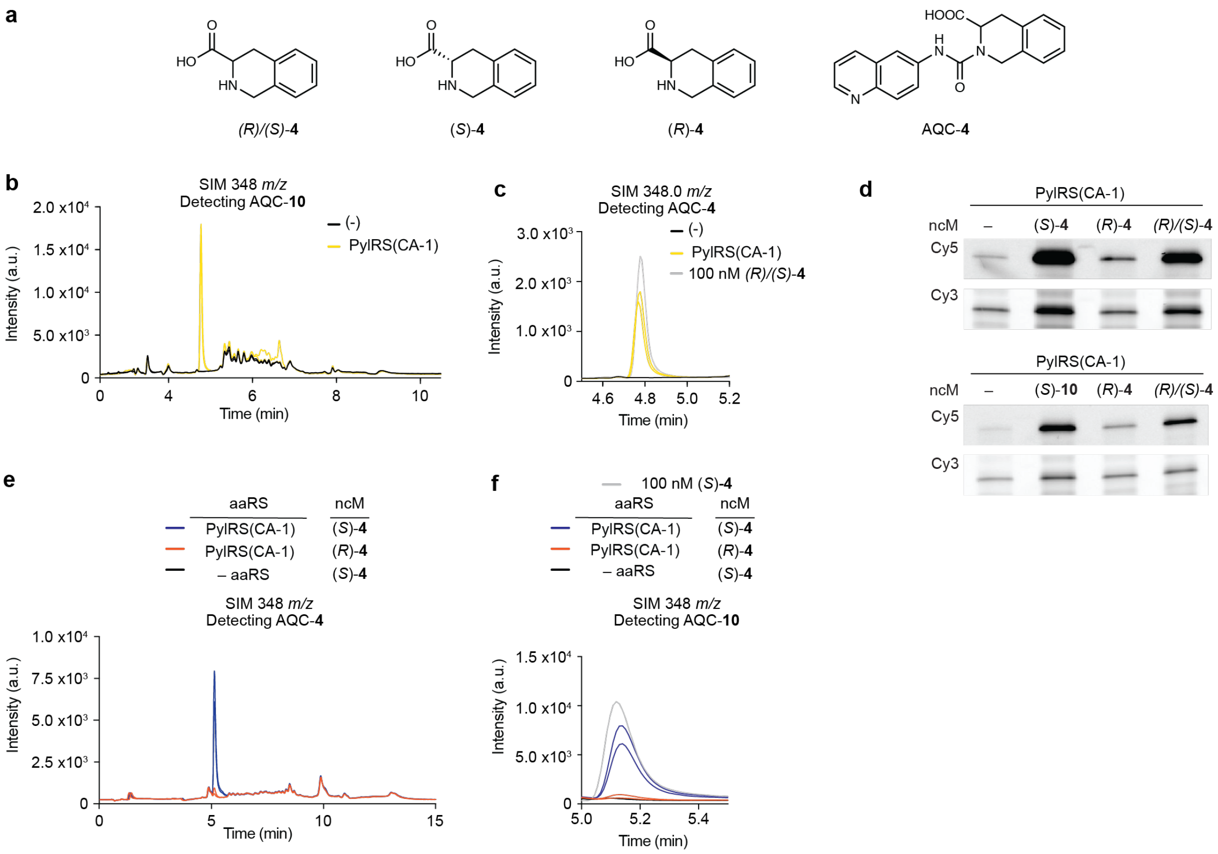


**Supplementary Figure 10. PylRS(CA-1) acylates tRNA$^{Pyl}$ with the *L*-enantiomer of** 4**.**

**a**, Chemical structures of racemic compound **4**, its enantiomerically pure forms *L*-**4** and *D*-**4** and its AQC derivative AQC-**4**. **b**, Single ion monitoring (SIM) LC-MS traces for AQC-**4**. tRNA$^{Pyl}$ was expressed with or without the synthetase PylRS(CA-1) and cells were grown in the presence of 4mM of **4**. A pulldown of RNA$^{Pyl}$ was performed and the acylated ncM was eluted by base treatment, AQC derivatised and analysed by single ion monitoring (M+H 348 *m/z*). Black traces show samples in which tRNA$^{Pyl}$ was expressed in the absence of PylRS. Yellow traces show samples in which tRNA$^{Pyl}$ was expressed in the presence of PylRS(CA-1). **c**, Zoom in on (**b**). A standard containing 100 nM of **4** was derivatised in parallel to the samples and analysed by LC-MS. It is shown for reference (grey trace). **d**, ftREX screen of PylRS(CA-1) with the enantiomerically pure forms of compound **4**. Experiments were performed using tRNA extracted from cells harbouring a pMB1 plasmid encoding tRNA$^{Pyl}$ and PylRS(CA-1). Cells were grown in absence of any ncM or in the presence of 4 mM of the shown ncM. The experiment was performed in duplicate. **e**, Analogous experiment to panel (**b**), but the experiment was performed with *L*-**4** (blue traces) or *D*-**4** (orange traces) instead of the racemic mixture of **4**. **f**, Zoom in on (**e**). A standard containing 100 nM of **4** was derivatised in parallel to the samples and analysed by LC-MS. It is shown for reference (grey trace). Graphs showing LC-MS traces show both replicates of each sample in the same graph.

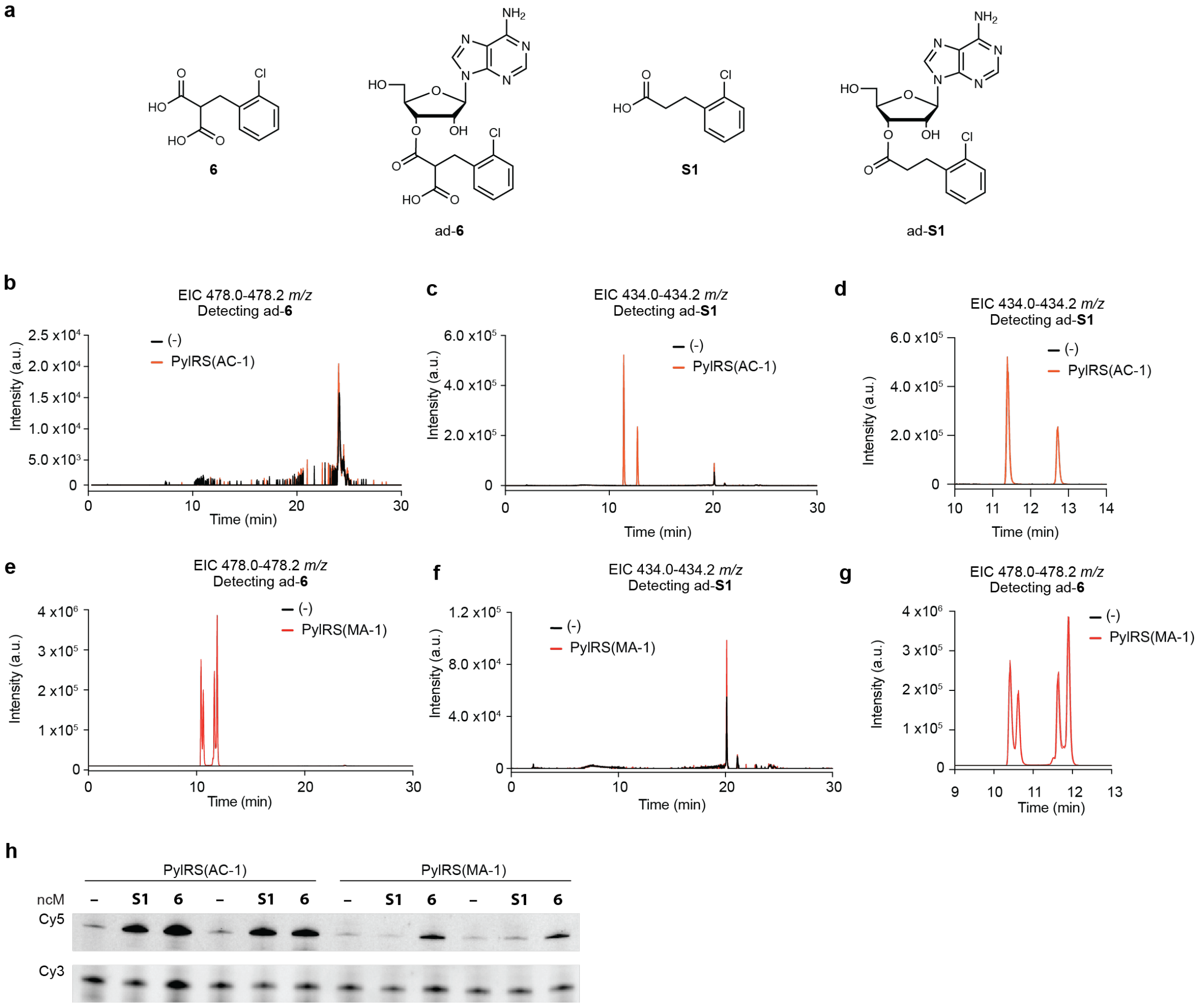


**Supplementary Figure 11. PylRS(MA-1) and PylRS(AC-1) acylate tRNA$^{Pyl}$ with** 6 **and** S1 **respectively.**

**a**, Chemical structures of compounds **6** and **S1** and their respective adenosine derivatives ad-**6** and ad-**S1**. **b**, **c** Full extracted ion chromatograms (EIC) for adenosine derivatives of **6** and **S1**. tRNA$^{Pyl}$ was expressed with or without the synthetase PylRS(AC-1) and cells were grown in the presence of **6** (4 mM). A pulldown of tRNA$^{Pyl}$ was performed and the acylated adenosine was eluted by RNAseA treatment and analysed by full scan LC-MS. Red traces show samples in which RNA$^{Pyl}$ was expressed in the presence of PylRS(AC-1). Black traces show samples in which tRNA$^{Pyl}$ was expressed in the absence of PylRS. **b**, shows an EIC (478.0-478.2 *m/z*) for the adenosine derivative of **6** (ad-**6**), which cannot be detected. **c**, shows an EIC (434.0-434.2 *m/z*) for the adenosine derivative of **S1** (ad-**S1**). The double peak is consistent with the presence of two regio-isomers derived from acylation at the 2' and 3' hydroxyls of adenosine. HR-MS of the observed EIC peaks in the presence of PylRS(AC-1) is consistent with ad-**S1** (calculated M+H = 434.1232 Da, observed *m/z* = 434.1226). **d,** zoom in of (**c**).

**e**, **f**, Analogous to (**b**), (**c**), but with synthetase variant PylRS(22.41), (**e**) EIC (478.0-478.2 m/z) for the adenosine derivative of **6** (ad-**6**). HR-MS of the observed EIC peaks in the presence of PylRS(MA-1) is consistent with ad-**6** (calculated M+H = 478.1130 Da, observed *m/z* = 478.1120). The quadruplet peak is consistent with the presence of two regio-isomers derived from acylation at the 2' or 3' hydroxyls of adenosine, each of which has two diastereomers derived from acylation with either the proR or the proS carboxylate. **f**, shows an EIC (434.0-434.2 *m/z*) for the adenosine derivative of **S1** (ad-**S1**) which cannot be detected. **g**, zoom in on (**e**). **h**, ftREX of PylRS variants. Experiments were performed using tRNA extracted from cells harbouring a pMB1 plasmid encoding $tRNA^{Pyl}$ and either PylRS(AC-1) or PylRS(MA-1). Cells were grown in absence of any ncM or in the presence of either **6** or **S1** (4 mM) as denoted in the figure. Experiments were performed in duplicates. Graphs showing LC-MS traces show both replicates of each sample in the same graph.

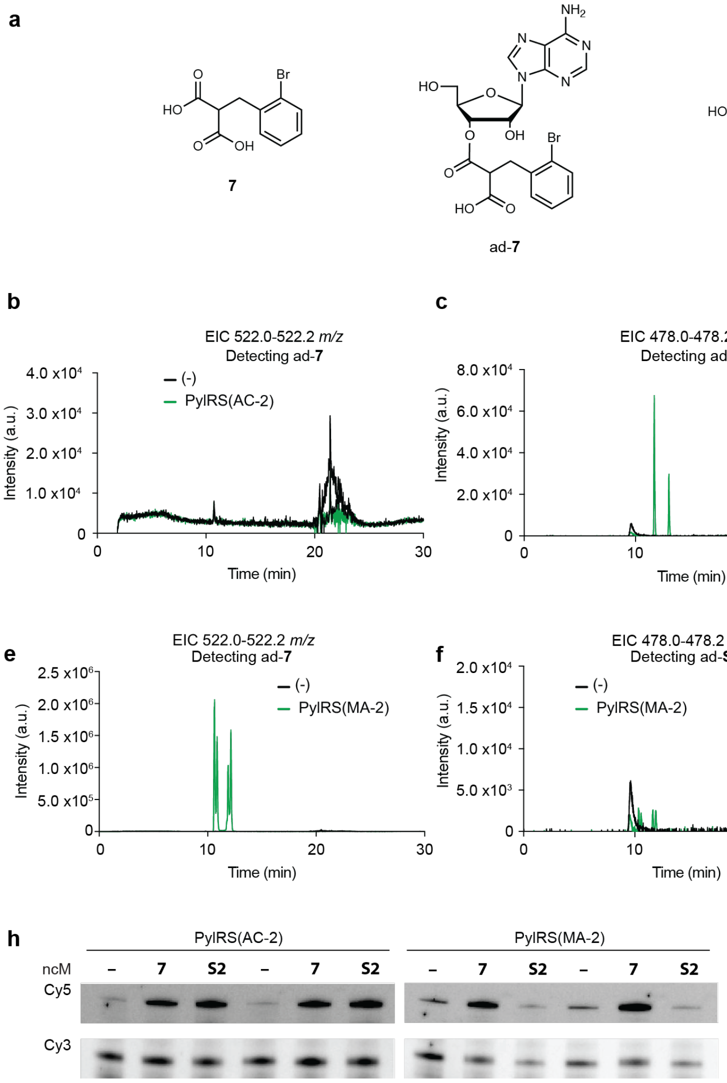


**Supplementary Figure 12. PylRS(MA-2) and PylRS(AC-2) acylate tRNA$^{Pyl}$ with** 7 **and** S2 **respectively.**

**a**, Chemical structures of compounds **7** and **S2** and their respective adenosine derivatives ad-**7** and ad-**S2**. **b**, **c** Characterisation of acylation by PylRS(AC-1) showing full extracted ion chromatograms (EIC) for adenosine derivatives of **7** and **S2**. tRNA$^{Pyl}$ was expressed with or without the synthetase PylRS(AC-2) and cells were grown in the presence of **7** (4 mM). A pulldown of RNA$^{Pyl}$ was performed and the acylated-adenosine was eluted by RNAseA treatment and analysed by full scan LC-MS. Green traces show samples in which tRNA$^{Pyl}$ was expressed in the presence of PylRS(AC-2). Black traces show samples in which tRNA$^{Pyl}$ was expressed in the absence of PylRS. **b**, shows an EIC (522.0-522.2 *m/z*) for the adenosine derivative of **7** (ad-**7**), which cannot be detected. **c**, shows an EIC (478.0-478.2 *m/z*) for the adenosine derivative of **S2** (ad-**S2**). The double peak is consistent with the presence of two regio-isomers derived from acylation at the 2' and 3' hydroxyls of adenosine. HR-MS of the observed EIC peaks in the presence of PylRS(AC-2) is consistent with ad-**S2** (calculated $^{79}$Br M+H = 478.0726 Da, $^{81}$Br M+H = 480.0706 Da, observed *m/z* = 478.0723 and 480.0703). **d,** zoom in on (**c**).

**e**, **f**, Analogous to (**b**), (**c**), but with synthetase variant PylRS(MA-2), (**e**) EIC (522.0-522.2 *m/z*) for the adenosine derivative of **7** (ad-**7**). The quadruplet peak is consistent with the presence of two regio-isomers derived from acylation at the 2' or 3' hydroxyls of adenosine, each of which has two diastereomers derived from acylation with either the proR or the proS carboxylate. HR-MS of the observed EIC peaks in the presence of PylRS(MA-2) is consistent with ad-**7** (calculated $^{79}$Br M+H = 522.0624 Da, $^{81}$Br M+H = 524.0604 Da, observed m/z = 522.0615 and 524.0594)). **f**, shows an EIC (478.0-478.2 m/z) for the adenosine derivative of **S2** (ad-**S2**) whch cannot be detected. **g**, Zoom in on (**e**). **h**, Fluoro-tREX of PylRS variants. Experiments were performed using tRNA extracted from cells harbouring a pMB1 plasmid encoding RNA$^{Pyl}$ and either PylRS(AC-2) or PylRS(MA-2). Cells were grown in absence of any ncM or in the presence of either **7** or **S2** (4 mM) as denoted in the figure. Experiments were performed in duplicates. Graphs showing LC-MS traces show both replicates of each sample in the same graph.

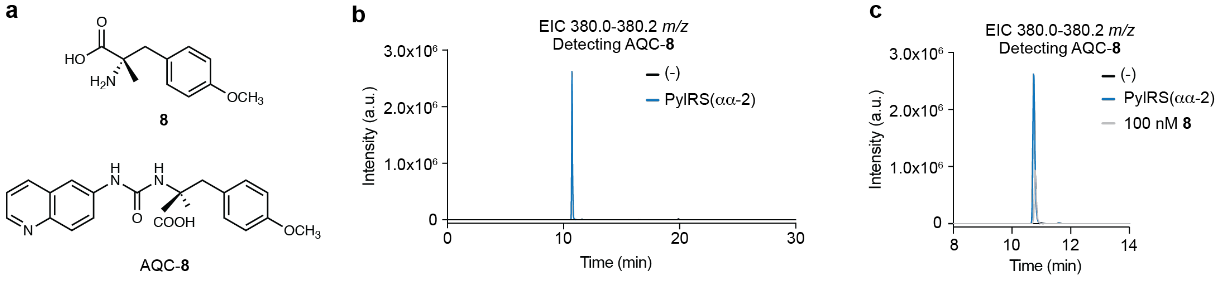


**Supplementary Figure 13. PylRS(αα-2) acylates tRNA$^{Pyl}$ with** 8**.**

**a**, Chemical structures of compound **8** and its AQC derivative AQC-**8**. **b**, Full extracted ion chromatogram (EIC 380.0-380.2 *m/z*) for AQC-**8**. tRNA$^{Pyl}$ was expressed with or without the synthetase PylRS(αα-2) and cells were grown in the presence of **8** (4 mM). A pulldown of tRNA$^{Pyl}$ was performed and the acylated ncM was eluted by treatment with base and AQC derivatised. Samples were analysed by full scan LC-MS. Blue traces show samples in which tRNA$^{Pyl}$ was expressed in the presence of PylRS(αα-2). Black traces show samples in which tRNA$^{Pyl}$ was expressed in the absence of PylRS. HR-MS of the observed EIC peaks in the presence of PylRS(αα-2) is consistent with AQC-**8** (calculated mass M+H = 380.1610 Da, observed *m/z* = 380.1602). **c,** zoom in of (**b**). A standard containing 100 nM of **8** was derivatised in parallel to the samples and analysed by LC-MS. It is shown for reference (grey trace). Two replicates of each sample are shown in each graph

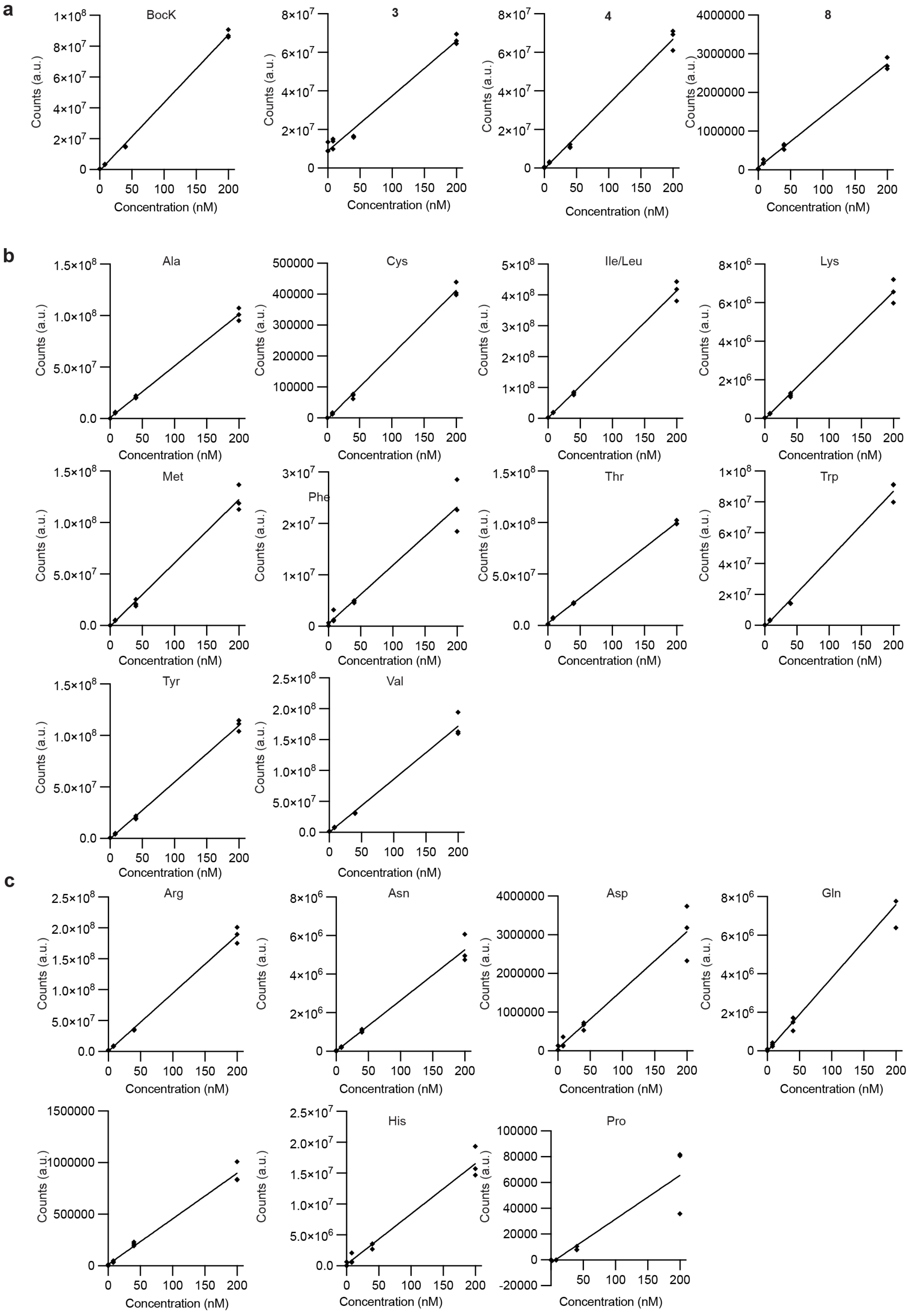


**Supplementary Figure 14. Standard curves used to quantify acylation of tRNA$^{Pyl}$ with ncMs and natural amino acids.**

**a**, Standard curves for **BocK**, compounds **3**, **4**, and **8.** Concentrations of 0 nM, 8 nM, 40 nM and 200 nM substrate were derivatised with AQC and analysed by LC-MS. Standard curves were generated by plotting the area under the curve of the isolated peak in the EIC for each compound against the concentration (EIC masses and retention times are specified in the Methods section). All experiments were performed in triplicates and individual data points are shown. **b**, Standard curves for 11 natural amino acids (Ala, Cys, Leu/Ile, Lys, Met, Phe, Trp, Tyr, Val). Concentrations of 0 nM, 8 nM, 40 nM and 200 nM amino acid were derivatised with AQC and analysed by LC-MS. Standard curves were generated by plotting the area under the curve of the isolated peak in the EIC for each compound against the concentration. All experiments were performed in triplicates. AQC derivatisation did not yield good standard curves at the low concentrations necessary to quantify acylation so we performed derivatisation with dansyl chloride to detect these amino acids. **c**, Standard curves for 7 natural amino acids (Arg, Asp, Asn, Gln, Glu, His, Pro). Concentrations of 0 nM, 8 nM, 40 nM and 200 nM amino acid were derivatised with dansyl chloride and analysed by LC-MS. Standard curves were generated by plotting the area under the curve of the isolated peak in the EIC for each compound against the concentration. Two sets of standard curves were used for experiments conducted on different days, only one set is shown here. All experiments were performed in triplicates. Ser and Gly could not be detected at the concentrations necessary to quantify acylation with ether of the two derivatisation methods.

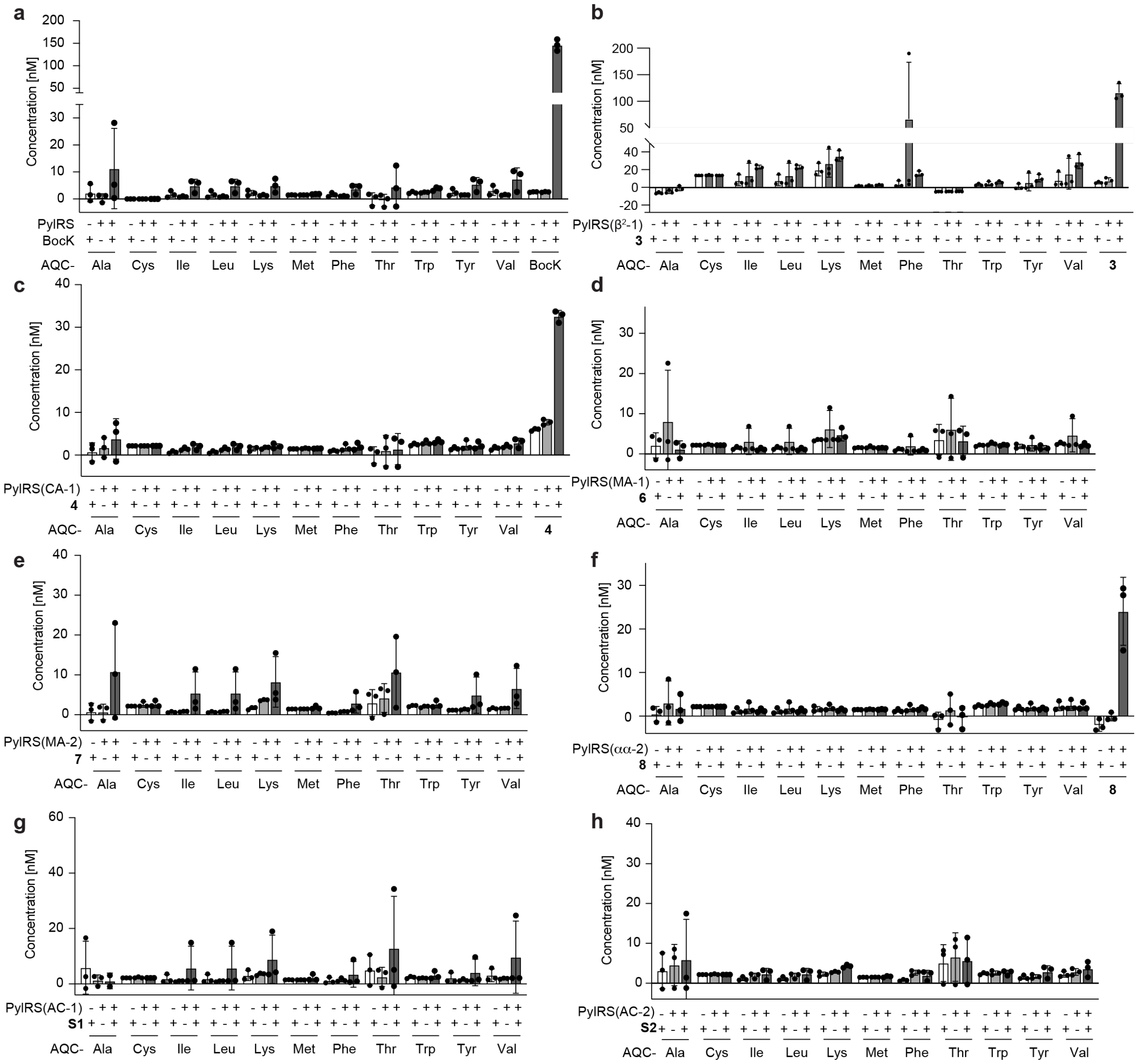


**Supplementary Figure 15. Quantification of tRNA$^{Pyl}$ acylation with ncMs and 11 natural amino acids for all PylRS variants by AQC derivatisation and LC-MS.**

To quantify tRNA$^{Pyl}$ acylation, *E. coli* cells containing a plasmid encoding both tRNA$^{Pyl}$ and a PylRS variant (or wildtype PylRS) were grown in presence or absence of 4 mM ncM or BocK. A pulldown of tRNA$^{Pyl}$ was performed and the monomers/amino acids acylated onto tRNA$^{Pyl}$ were eluted by base treatment and AQC derivatised. For each compound, the respective peak in the EIC was identified and integrated. The integrated signal was used to determine the concentration of monomer using standard curves (see **Supplementary Fig. 14**). Abundance of 11 natural amino acids (Ala, Cys, Leu/Ile, Lys, Met, Phe, Trp, Tyr, Val) was quantified for every sample in the same way. Background levels of amino acids were determined by expressing tRNA$^{Pyl}$ in the absence of the PylRS variant but in presence of 4 mM ncM or BocK. All experiments were performed in triplicates. The bars represent the mean, individual data points are shown as black dots and error bars represent the standard deviation. These measurement are steady state measurements that measure acylation at a fixed time-point. As tRNAs acylated with natural amino acids may be consumed by translation/hydrolysis at a rate that is higher than that of some of the ncMs (especially those not bearing an amino group), this might mean the apparent level of acylation with natural amino acids is lower. **a**, Quantification of $tRNA^{Pyl}$ acylation by wild-type PylRS. Only acylation with BocK is detected above background levels. We do not observe acylation with natural amino acids at levels above background.

**b**, Quantification of $tRNA^{Pyl}$ acylation by PylRS($\beta^2$-1) with compound **3**. The AQC adduct of **3** is the most abundant species. In one replicate we detected high levels of acylation with Phe. However, in the presence of **3** we did detect only minor levels of acylation with canonical amino acids.

**c**, Quantification of $tRNA^{Pyl}$ by PylRS(CA-1) with compound **4**. Only acylation with **4** is detected above background levels.

**d**, Quantification of $tRNA^{Pyl}$ acylation by PylRS(MA-1). Compound **6** cannot be detected using this method as it cannot be derivatised with AQC. The acylation levels for the quantified canonical amino acids are low and comparable to the levels in (**a**).

**e**, Quantification of $tRNA^{Pyl}$ acylation by PylRS(MA-2). Compound **7** cannot be detected using this method as it cannot be derivatised with AQC. The acylation levels for the quantified canonical amino acids is low and comparable to the levels in (**a**).

**f**, Quantification of $tRNA^{Pyl}$ acylation by PylRS($\alpha\alpha$-2). Only acylation with **8** is detected above background levels.

**g**, Quantification of $tRNA^{Pyl}$ acylation by PylRS(AC-1). Compound **S1** cannot be detected using this method as it cannot be derivatised with AQC. The acylation levels for the quantified canonical amino acids is low and comparable to the levels in (**a**). One replicate for Alanine detection was removed from the analysis due to a technical error.

**h**, Quantification of tRNA$^{Pyl}$ acylation by PylRS(AC-2). Compound **S2** cannot be detected using this method as it cannot be derivatised with AQC. The acylation levels for the quantified canonical amino acids are low and comparable to the levels in (**a**).

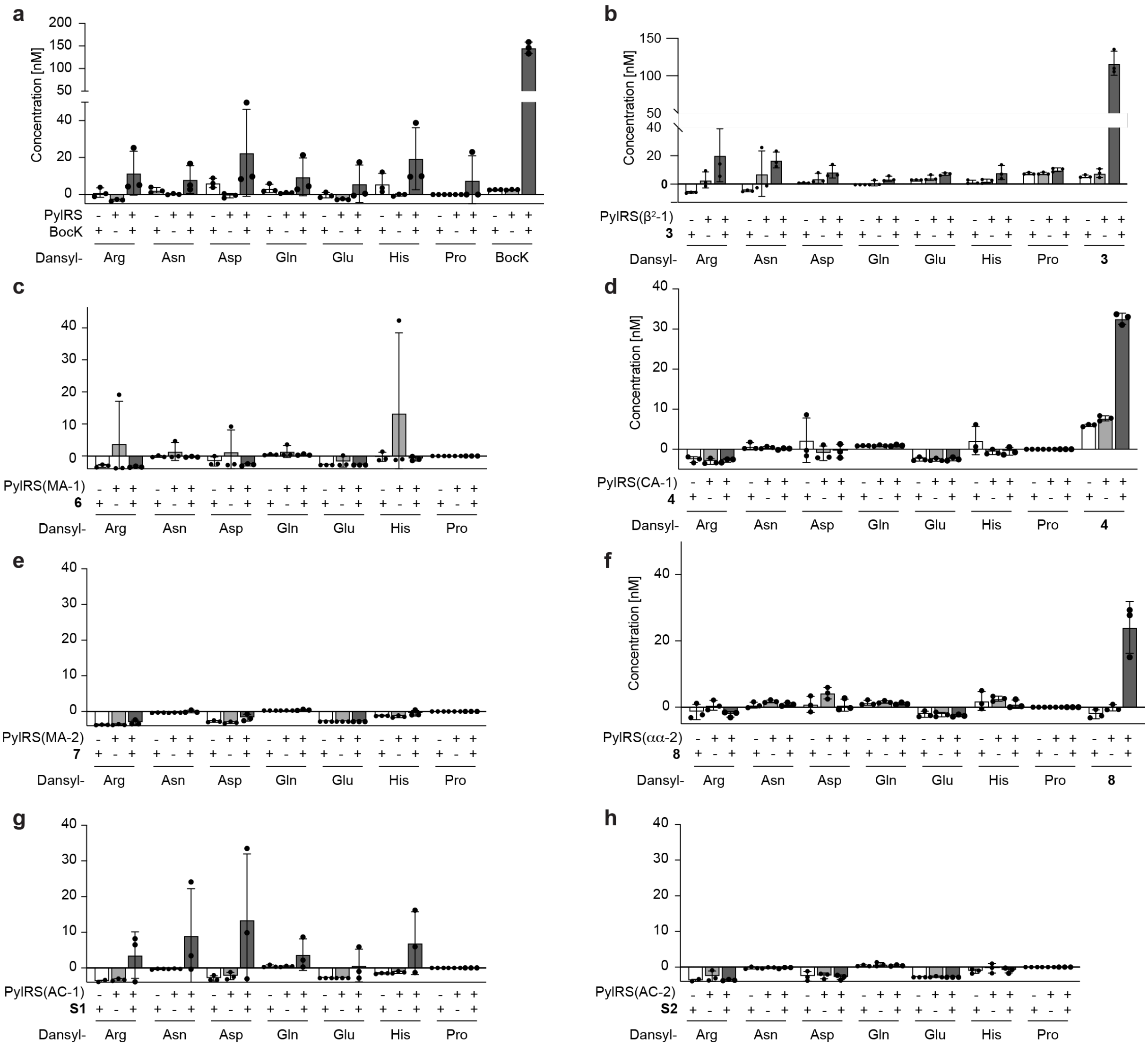


**Supplementary Figure 16. Quantification of tRNA$^{Pyl}$ acylation with ncMs and 7 natural amino acids for all PylRS variants by dansyl chloride derivatisation and LC-MS.**

To quantify tRNA$^{Pyl}$ acylation, *E. coli* cells containing a plasmid encoding both RNA$^{Pyl}$ and a PylRS variant (or wildtype PylRS) were grown in presence or absence of 4 mM ncM (or BocK). A pulldown of tRNA$^{Pyl}$ was performed and the monomer acylated onto tRNA$^{Pyl}$ was eluted by base treatment and derivatised with dansyl chloride. For each compound, the respective peak in the EIC was identified and integrated. The integrated signal was used to determine the concentration of monomer using standard curves (see **Supplementary Fig. 14**). The abundance of 7 natural amino acid (Arg, Asn, Asp, Gln, Glu,

His, Pro) was analysed and quantified for every sample in the same way. Unfortunately, we were unable to quantify serine or glycine using this method (or the analogous one described in **Supplementary Fig 15**). The data points for ncMs are from AQC-derivatised experiment and replotted from **Supplementary Figure 15**. All experiments were performed in triplicates. The bars represent the mean, individual data points are shown as black dots and error bars represent the standard deviation. These measurement are steady state measurements that measure acylation at a fixed time-point. As tRNAs acylated with natural amino acids may be consumed by translation/hydrolysis at a rate that is higher than that of some of the ncMs (especially those not bearing an amino group), this might mean the apparent level of acylation with natural amino acids is lower.

**a**, Quantification of $tRNA^{Pyl}$ acylation by wild-type PylRS. Only acylation with BocK is detected above background levels.

**b**, Quantification of $tRNA^{Pyl}$ acylation by PylRS($\beta^2$-1) with compound **3**. Only acylation with **3** is detected above background levels.

**c**, Quantification of $tRNA^{Pyl}$ acylation by PylRS(CA-1) with compound **4**. Only acylation with **10** is detected above background levels.

**d**, Quantification of $tRNA^{Pyl}$ acylation by PylRS(MA-1).The acylation levels for the quantified canonical amino acids are low and comparable to the levels in (**a**).

**e**, Quantification of $tRNA^{Pyl}$ acylation by PylRS(MA-2). The acylation levels for the quantified canonical amino acids are low and comparable to the levels in (**a**).

**f**, Quantification of $tRNA^{Pyl}$ acylation by by PylRS($\alpha\alpha$-2). Only acylation with **8** is detected above background levels.

**g**, Quantification of $tRNA^{Pyl}$ acylation by PylRS(AC-1). The acylation levels for the quantified canonical amino acids are low and comparable to the levels in (**a**).

**h**, Quantification of $tRNA^{Pyl}$ acylation by PylRS(AC-2). The acylation levels for the quantified canonical amino acids are low and comparable to the levels in (**a**).

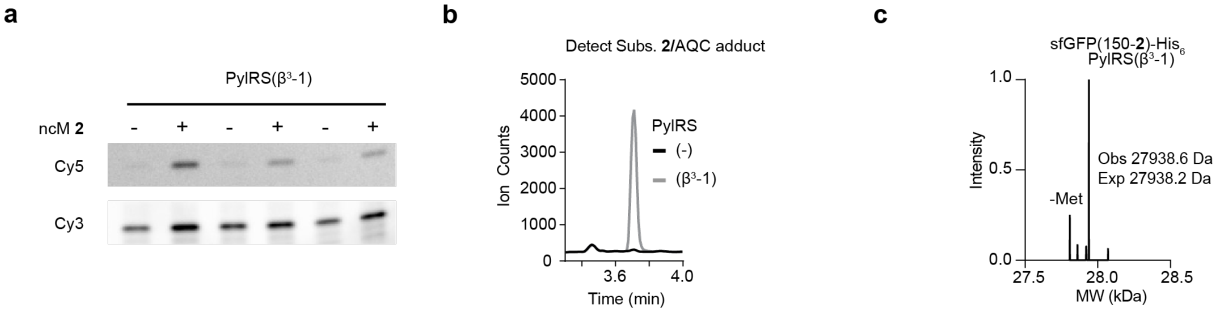


**Supplementary Figure 17. PylRS(β$^3$-1) selectively charges** 2 **onto tRNA$^{Pyl}$.**

**a**, Fluoro-tREX of the PylRS(β$^3$-1)/tRNA$^{Pyl}$ pair expressed in the presence and absence of substrate **2**.

**b**, Single ion monitoring (SIM) detection of AQC adduct of substrate **2** in samples resulting from pulldown of tRNA$^{Pyl}$ expressed in the presence of **2** and with (grey trace) or without (black trace) the aaRS variant PylRS(β$^3$-1). The trace shows charging of the correct monomer onto tRNA$^{Pyl}$ by PylRS(β$^3$-1). **c**, Intact ESI-MS of sfGFP containing ncMs **2** at position 150. Cells contained a plasmid encoding for PylRS(β$^3$-1) as well as tRNA$^{Pyl}$ and were grown in the presence of **2**.

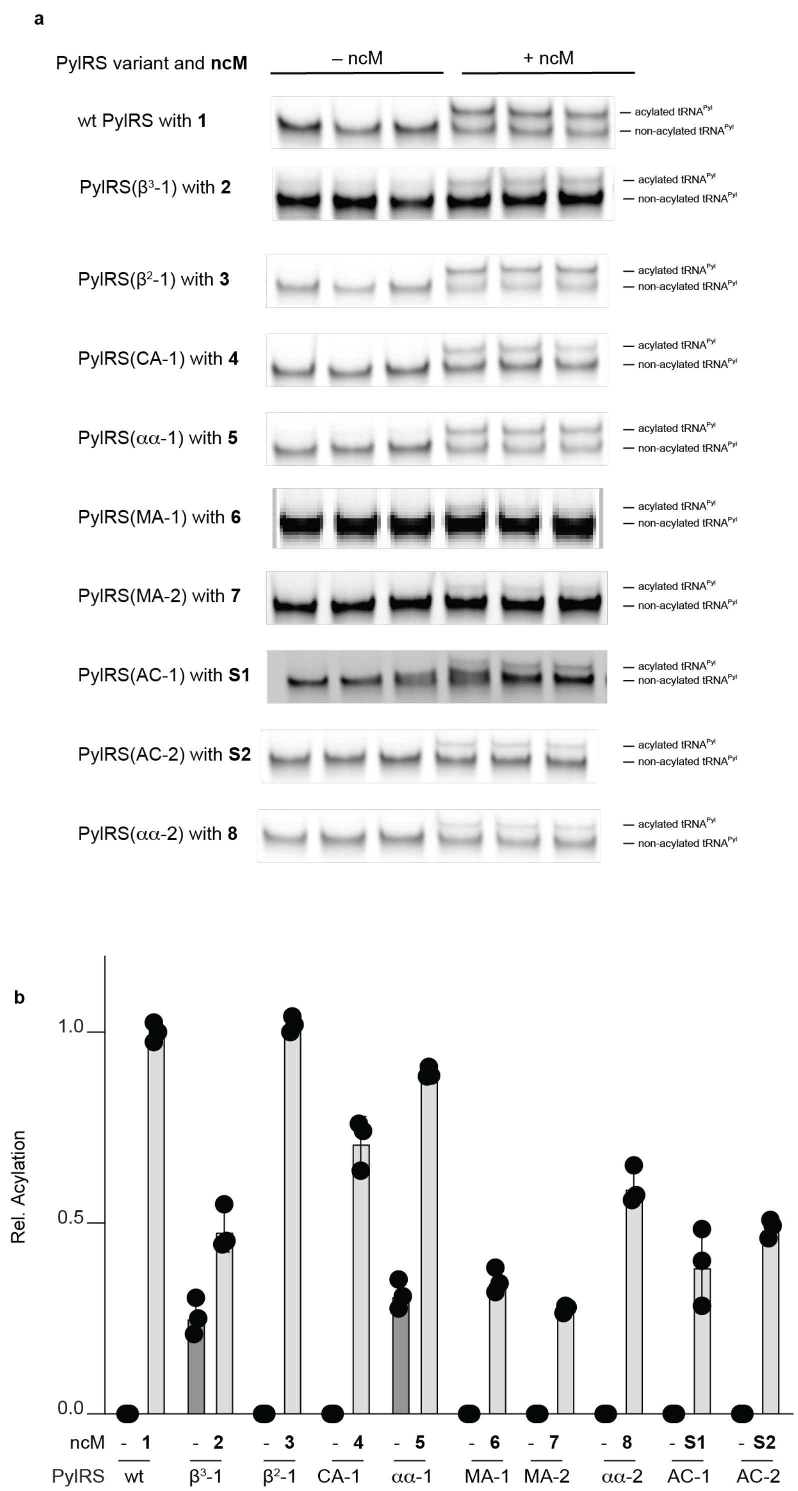


**Supplementary Figure 18. *In vivo* acylation activity of PylRS variants analysed by tREX.**

**a,** tREX gel for $RNA^{Pyl}$ extracted from cells containing the specified PylRS in the presence of 4 mM of their respective ncM. Experiments were carried out in triplicates. **b**, Quantification of gel bands shown in **a** by densitometry. The acylation activity is determined as the ratio of the signal intensity of the upper band (acylated $tRNA^{Pyl}$) divided by the sum of the signal intensity of the upper and lower bands (nonacylated $tRNA^{Pyl}$). Acylation levels are plotted as a fraction of wildtype PylRS acylation with 2 mM AllocK (47.4 ± 1.2%) which was set to 1. all monomers were added to cells at a concentration of 4 mM except for **5** (2 mM). Relative acylation in the presence of monomers was measured as 48.2 ± 5.7 % for **2**, 102.1 ± 2.1% for **3**, 71.3 ± 6.6% for **4**, 89.4 ± 1.4 % for **5** 33.7 ± 3.1% for **6**, 26.7 ± 1.0% for **7**, 59.5 ± 4.9% for **8**, 39.0 ± 10.1 % for **S1**, 48.7 ± 2.5 % for **S2**. The bars represent the mean, individual data points are shown as black dots and error bars represent the standard deviation. These measurement of acylation activity are steady state measurements that measure acylation at a fixed time-point in the absence of consumption by natural translation. They do not necessarily report on the exact catalytic efficiency of each enzyme in vitro, but they allow measurement of relative acylation in the conditions relevant to genetic code expansion in vivo. Steady state-acylation levels are an aggregate measurement and are affected by: enzyme expression levels, enzyme activity, monomer transport efficiency and deacylaton rate (dependent on efficiency of the given monomer in translation and stability of the acylated tRNA to hydrolysis). Nonetheless all these parameters are also relevant to acylation when the tRNAs are actively involved in translation. The end-point nature of these measurements means that enzymes that might acylate the tRNA at a lower rate can still lead to accumulation of the acylated tRNA, potentially appearing like they acylate at a rate that is higher than what would be the case if the monomers were actively consumed in translation.

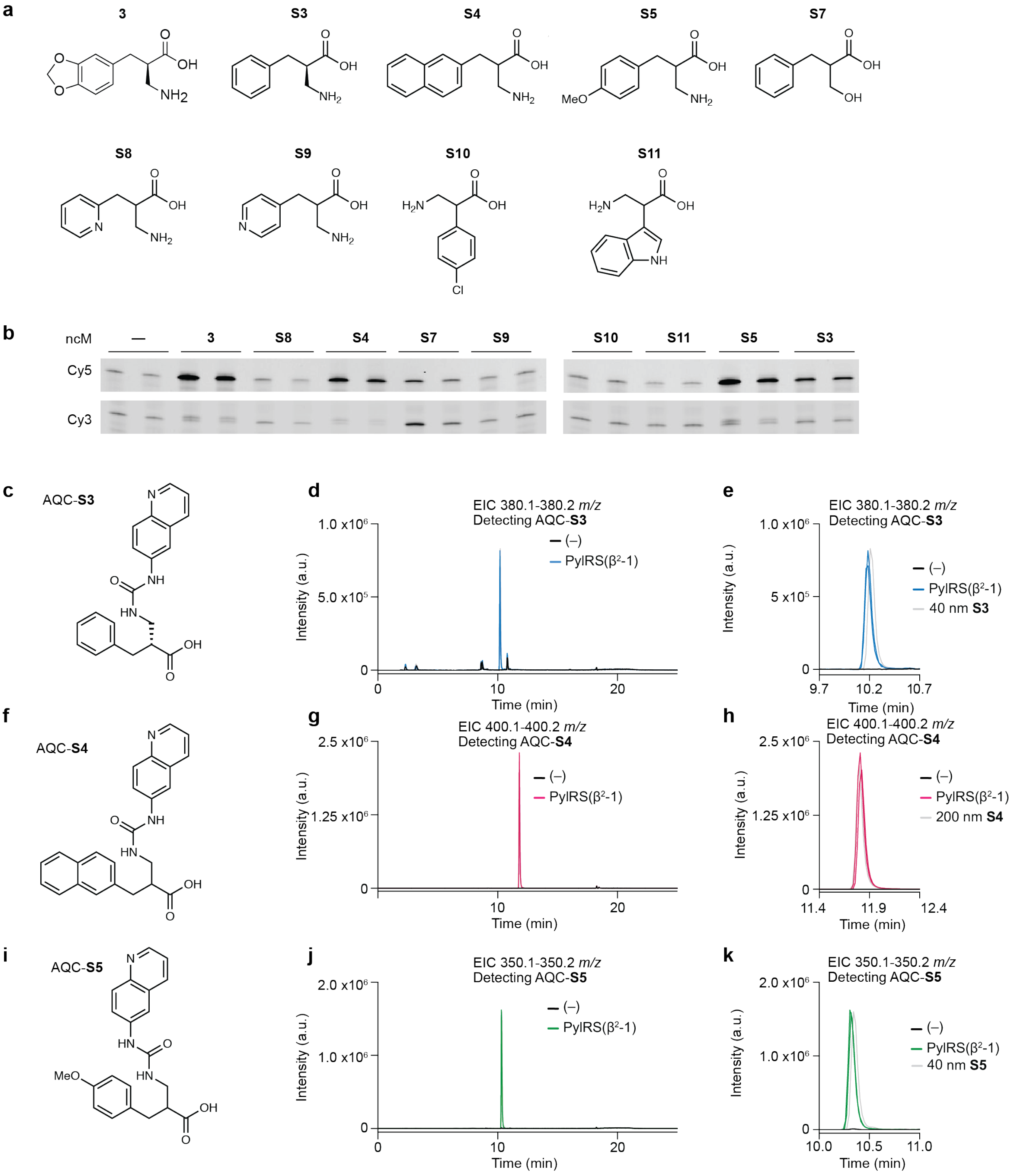


**Supplementary Figure 19. Screening the activity of PylRS(β$^2$-1) with a panel of β$^2$-amino acids.**

**a**, Chemical structure of the compounds screened with PylRS(β$^2$-1). **b**, Fluoro-tREX screen of PylRS(β$^2$-1) with a panel of β$^2$-amino acids. Experiments were performed using tRNA extracted from cells harbouring a pMB1 plasmid encoding tRNA$^{Pyl}$ and PylRS(β$^2$-1). Cells were grown in absence of any ncM or in the presence of the shown ncM (4 mM). The experiment was performed in duplicate.

**c**, **f, i**, Chemical structure of AQC-**S3,** AQC-**S4** and AQC-**S5**. **d**, Extracted ion chromatogram (EIC) LC-MS traces for AQC-**S3**. RNA$^{Pyl}$ was expressed with or without the synthetase PylRS($\beta^2$-1) and cells were grown in the presence of **S3** (4 mM). A pulldown of tRNA$^{Pyl}$ was performed and the acylated ncM was eluted by base treatment, AQC derivatised and analysed by LC-MS. Blue traces show samples in which tRNA$^{Pyl}$ was expressed in the presence of PylRS($\beta^2$-1). Black traces show samples in which tRNA$^{Pyl}$ was expressed in the absence of PylRS. **e**, zoom in on (**d**). A standard containing 40 nM of **S3** was derivatised in parallel to the samples and analysed by LC-MS. It is shown for reference (grey trace). **g**, as in (**d**) but for AQC-**S4**. Pink traces show samples in which tRNA$^{Pyl}$ was expressed in the presence of PylRS($\beta^2$-1). Black traces show samples in which tRNA$^{Pyl}$ was expressed in the absence of PylRS. **h**, zoom in on (**g**). **j**, As is (**d**), but the experiment was performed with **S5** (green traces). **k**, Zoom in on (**j**). Graphs showing LC-MS traces show both replicates of each sample in the same graph

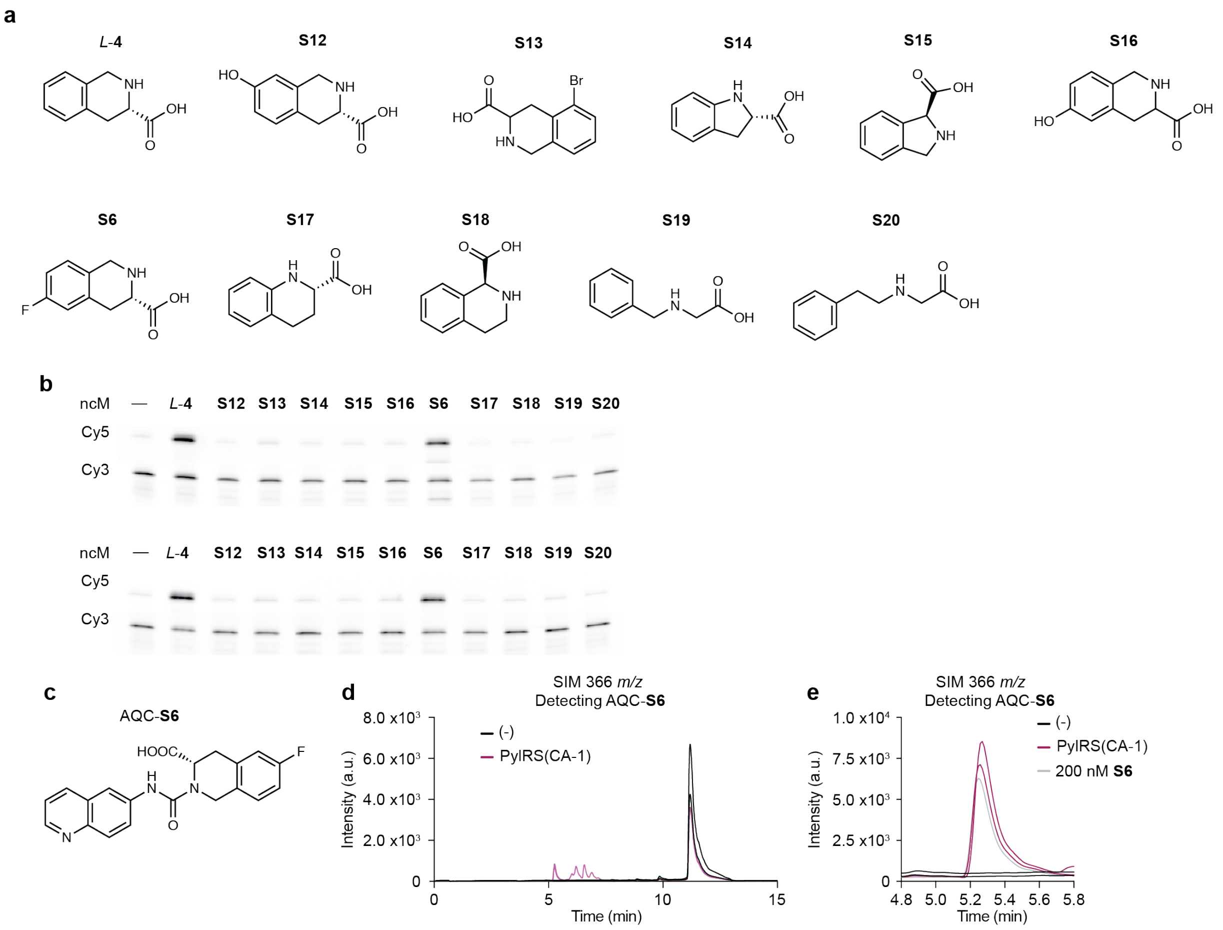


**Supplementary Figure 20. PylRS(CA-1) acylates tRNA$^{Pyl}$ with** S6.

**a**, chemical structure of the compounds screened with PylRS(CA-1). **b**, Fluoro-tREX of PylRS(CA-1) with a panel of N-alkylated compounds. Experiments were performed using tRNA extracted from cells harbouring a pMB1 plasmid encoding tRNA$^{Pyl}$ and PylRS(CA-1). Cells were grown in absence of any ncM or in the presence of 4 mM of the shown ncM. The experiment was performed in duplicate. **c**, chemical structure of the AQC derivative of **S6**, AQC-**S6**. **d**, Single ion monitoring (SIM) LC-MS traces for AQC-**S6**. tRNA$^{Pyl}$ was expressed with or without the synthetase PylRS(CA-1) and cells were grown in the presence of 4mM of **S6**. A pulldown of tRNA$^{Pyl}$ was performed and the acylated ncM was eluted by base treatment, AQC derivatised and analysed by single ion monitoring. Purple traces show samples in which tRNA$^{Pyl}$ was expressed in the presence of PylRS(CA-1). Black traces show samples in which tRNA$^{Pyl}$ was expressed in the absence of PylRS. **e**, zoom in on (**d**). A standard containing 200 mM of **S6** was derivatised in parallel to the samples and analysed by LC-MS, shown for reference (grey trace). Graphs showing LC-MS traces show both replicates of each sample in the same graph.

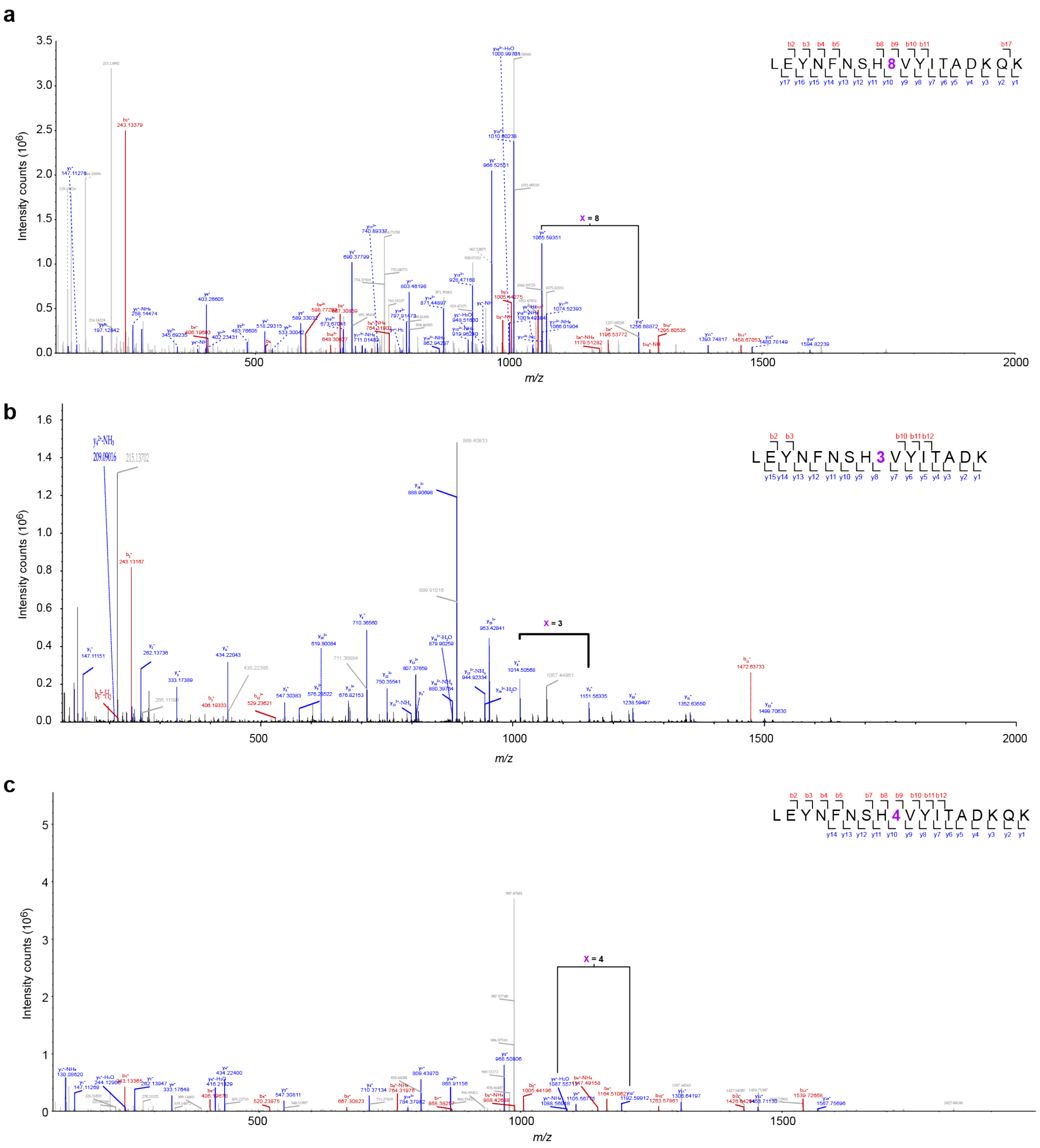


**Supplementary Figure 21. Tandem mass spectrometry analysis of sfGFP(150-X)His$_6$ containing ncMs** 8, 3, **and** 4.

MS/MS spectra of purified sfGFP containing an ncM at position 150. All proteins were expressed in the presence of 4 mM ncM and its respective PylRS variant. Fragmentation of the precursor ions yield a series of b ions (red) and y ions (blue) which confirm the incorporation of the correct ncM at position 150 of GFP. **a**, MS/MS spectrum of sfGFP(150-**8**)His$_6$ produced from cells containing PylRS(αα-2) and tRNA$^{Pyl}$ grown with **8**. **b**, MS/MS spectrum of purified sfGFP(150-(*R*)-**3**)His$_6$ produced from cells containing PylRS($\beta^2$-1) and $tRNA^{Pyl}$ grown with (*R*)-**3**. **c**, MS/MS spectrum of purified sfGFP(150-*L*-**4**)$His_6$ produced from cells containing PylRS(CA-1) and $tRNA^{Pyl}$ grown with *L*-**4**.

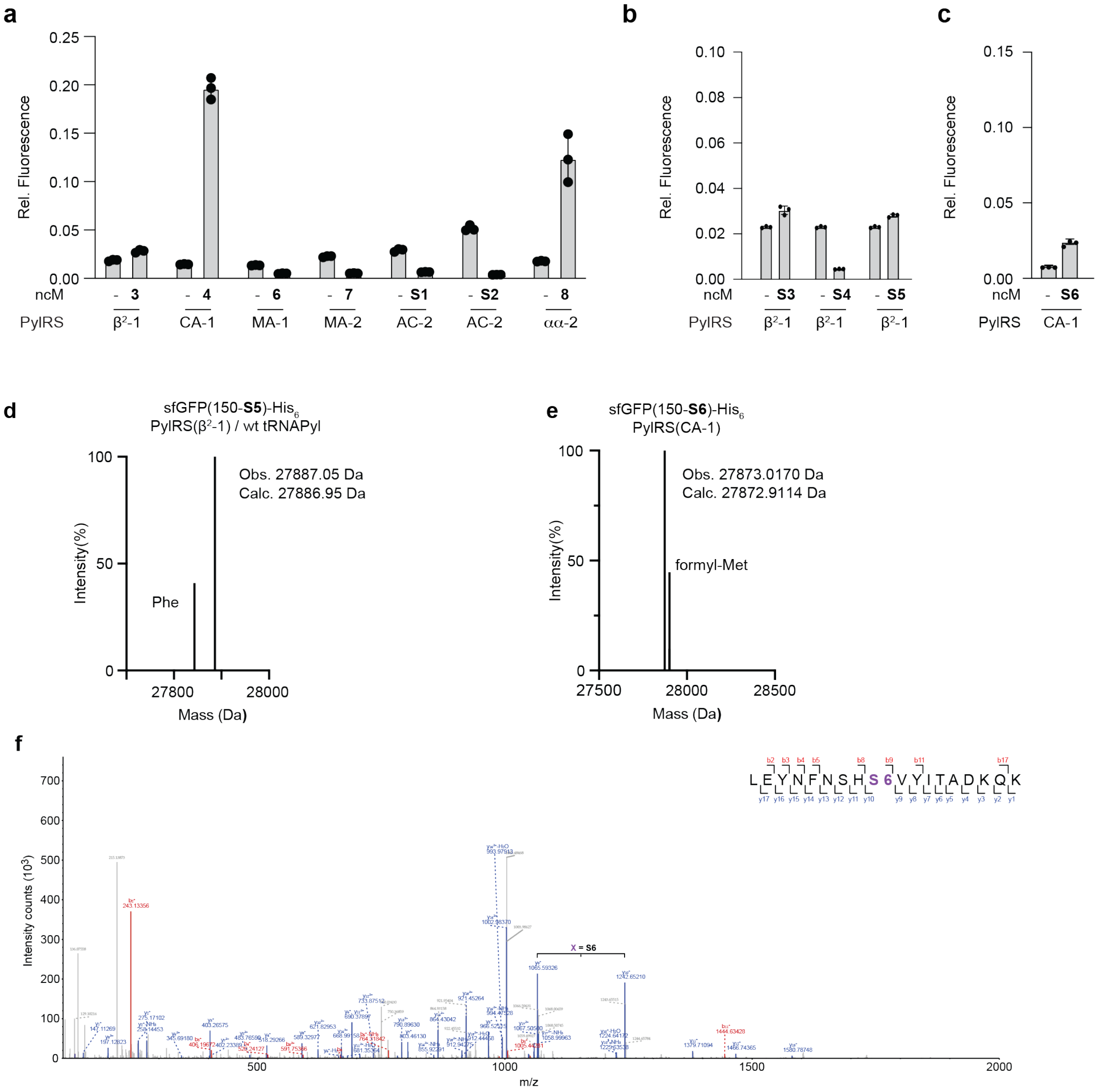


**Supplementary Figure 22. Selected PylRS variants enable the site-specific incorporation of** S5 **and** S6 **into proteins.**

**a-c**, GFP expression from cells containing *sfGFP(150TAG)His*$_6$, tRNA$^{Pyl}$ and the indicated variants of PylRS. Cells were grown in the presence and absence of the indicated ncM (4 mM). GFP fluorescence levels are shown relative to those from cells containing *sfGFP(150TAG)His*$_6$, tRNA$^{Pyl}$ and wt PylRS expressed in the presence of 2 mM of **1** (set to 1). The bar graphs show the mean of three biological replicates, individual data points are shown as black dots and error bars show the standard deviation about the mean. **d**, Deconvoluted mass spectrum of sfGFP(150-**S5**)His$_6$ purified from cells harbouring PylRS($\beta^2$-1) and *sfGFP(150TAG)His$_6$* grown with **S5**. Incorporation of **S5** is observed alongside mis-incorporation of phenylalanine. **e**, Deconvoluted mass spectrum of sfGFP(150-**S6**)His$_6$ purified from cells harbouring PylRS(CA-1) and *sfGFP(150TAG)His$_6$* grown with **S6**. **f**, MS/MS spectrum of purified sfGFP(150-**S6**)His$_6$ produced from cells containing PylRS(CA-1) and tRNA$^{Pyl}$ grown with **S6**.

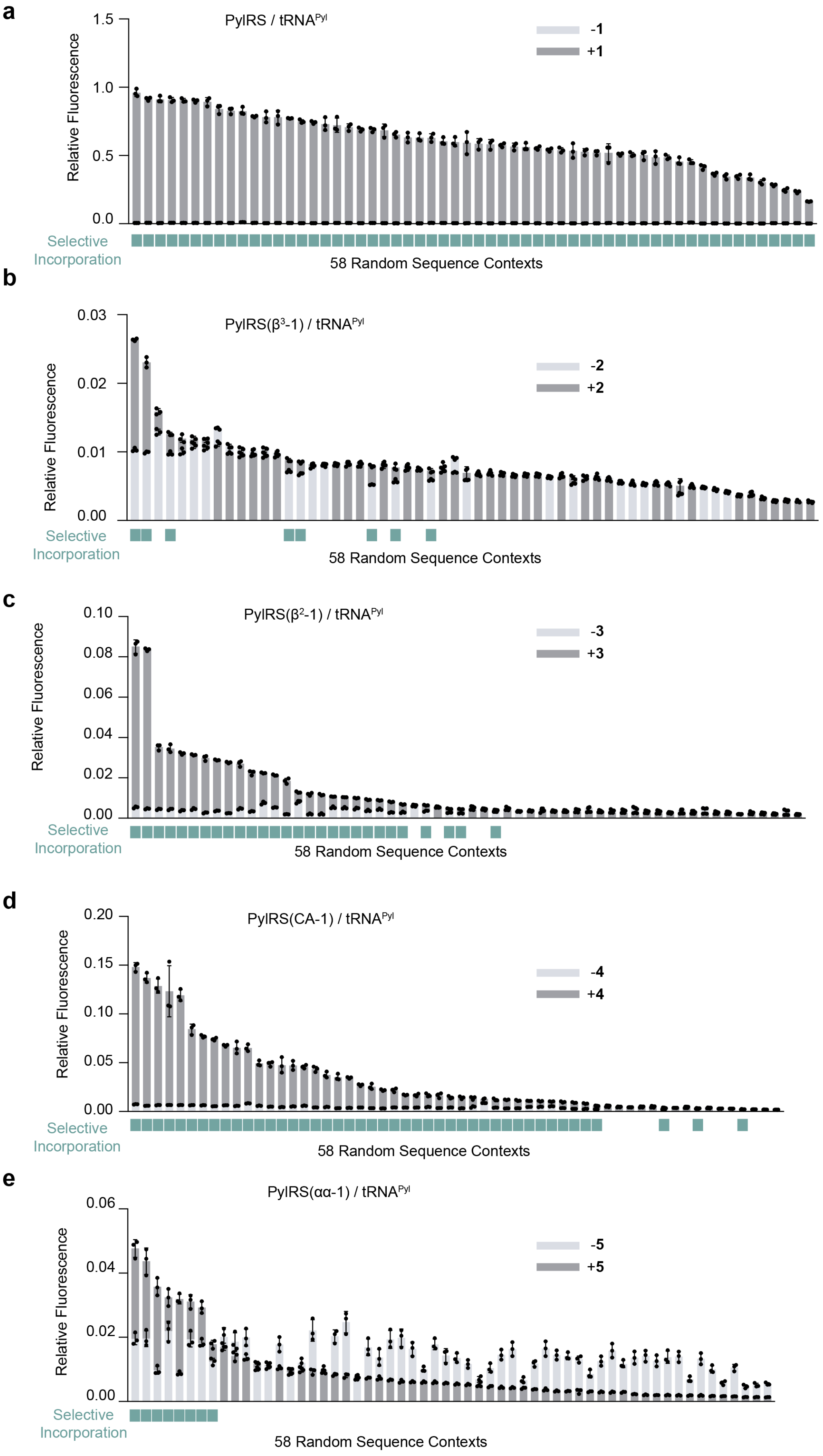
a
PylRS / tRNA^Pyl
-1
+1
Relative Fluorescence
1.5
1.0
0.5
0.0
Selective Incorporation
58 Random Sequence Contexts
b
PylRS(β³-1) / tRNA^Pyl
-2
+2
Relative Fluorescence
0.03
0.02
0.01
0.00
Selective Incorporation
58 Random Sequence Contexts
c
PylRS(β²-1) / tRNA^Pyl
-3
+3
Relative Fluorescence
0.10
0.08
0.06
0.04
0.02
0.00
Selective Incorporation
58 Random Sequence Contexts
d
PylRS(CA-1) / tRNA^Pyl
-4
+4
Relative Fluorescence
0.20
0.15
0.10
0.05
0.00
Selective Incorporation
58 Random Sequence Contexts
e
PylRS(αα-1) / tRNA^Pyl
-5
+5
Relative Fluorescence
0.06
0.04
0.02
0.00
Selective Incorporation
58 Random Sequence Contexts

**Supplementary Figure 23. Individually characterized sequence context reporters.**

**a-e,** Fluorescence values of 58 random sequence context reporters expressed with wt PylRS (**a**), PylRS($\beta^3$-1) (**b**), PylRS($\beta^2$-1) (**c**), PylRS(CA-1) (**d**), or PylRS(αα-1) (**e**) in the presence (light grey) or absence (dark grey) of **1** (**a**), **2** (**b**), **3** (**c**), **4** (**d**), or **5** (**e**). Reporters that supported selective incorporation (ratio of fluorescence in the presence of ncM divided by fluorescence in the absence of ncM >1.2) are highlighted with a green square. The threshold corresponding to the fluorescence of the context identified in Supplementary Fig. 25 is indicated for each ncM. GFP fluorescence levels are shown relative to those from cells containing *sfGFP(150TAG)His$_6$*, tRNA$^{Pyl}$ and wt PylRS expressed in the presence of 2 mM of **1** (set to 1).

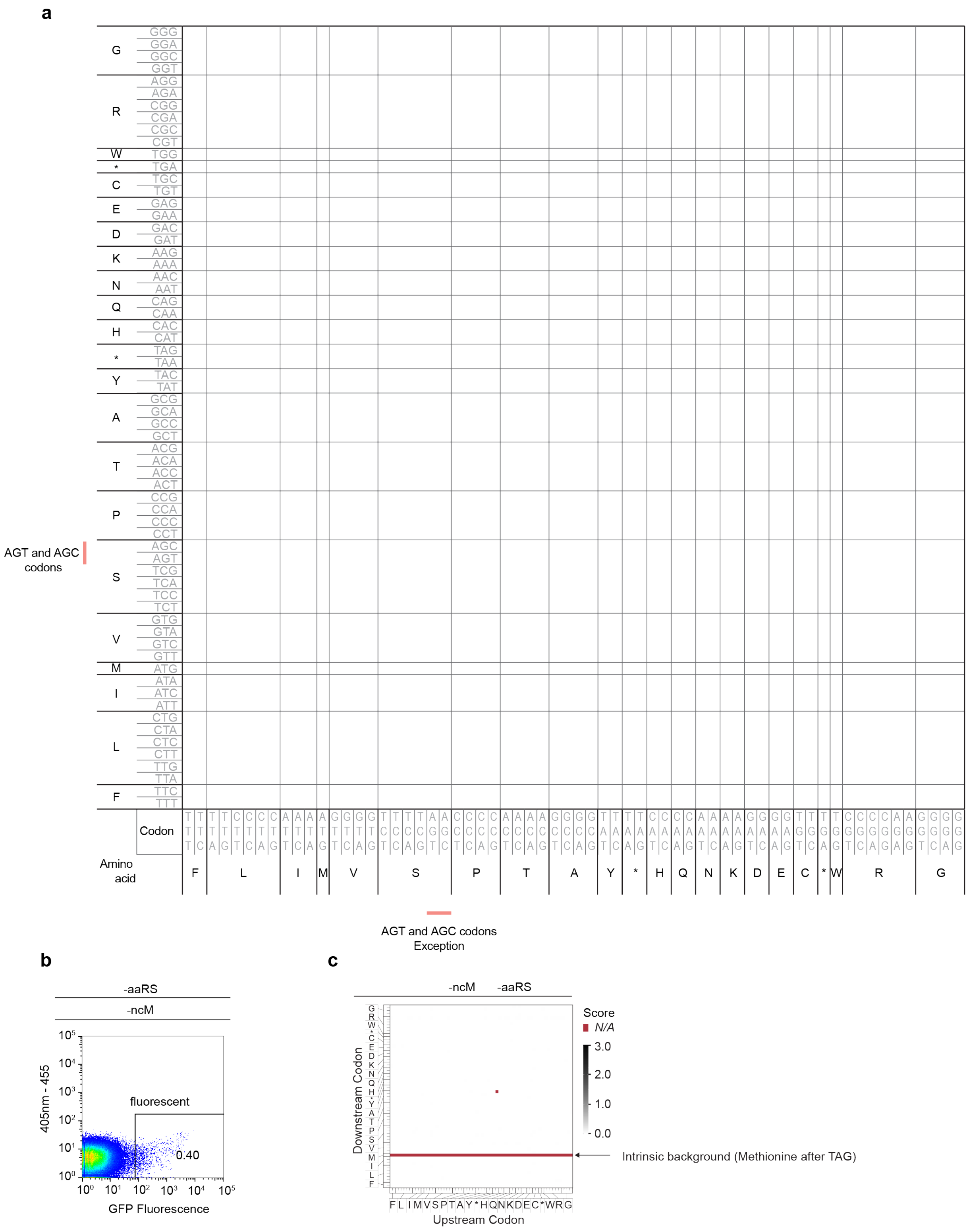


**Supplementary Figure 24. Arrangement of the 64x64 codons for sequence context mapping.**

**a**, Arrangement of 64 codons along one axis used for the representation of incorporation efficiency matrices. Codons are hierarchically arranged by: the second base of the codon, then the first base of the codon, and lastly the third base of the codon. For each base (in all three positions), the employed hierarchy is: first T, then C, then A, and lastly G. The AGT and AGC serine codons and the AGA and AGG arginine codons constitute exceptions and have been placed next to their corresponding synonymous codons. Overall, this arrangement results in codons encoding the same amino acids being grouped together.

**b,** Fluorescence distribution measured by FACS of N-terminal sequence context library expressed in the absence of any aaRS and ncM.

**c**, Incorporation efficiency heat maps for translation the N-terminal sequence context library expressed in the absence of any aaRS and ncM, i.e. background.

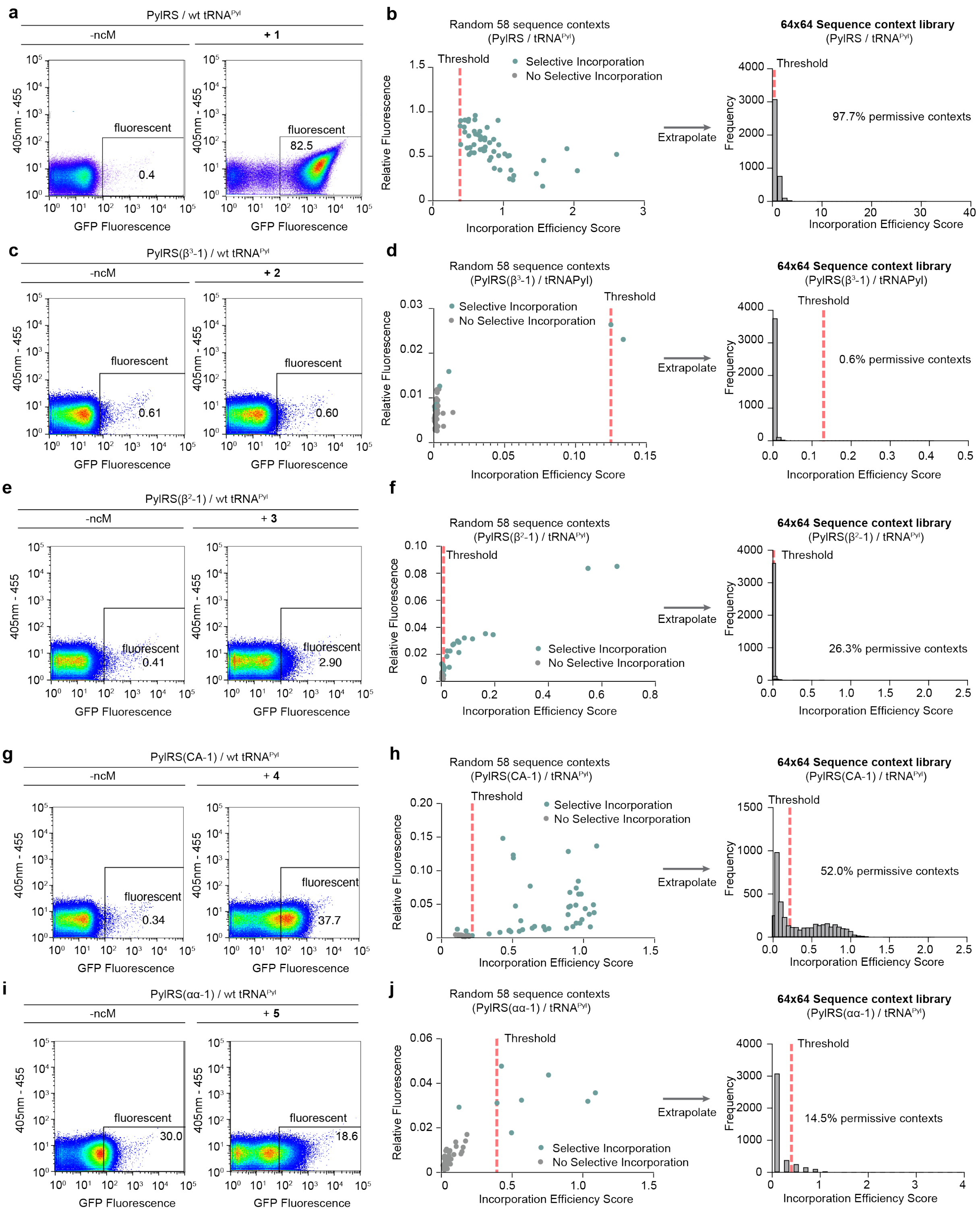


**Supplementary Figure 25. FACS and NGS analysis of sequence context libraries.**

**a,c,e,g,i**, Fluorescence distribution measured by FACS of N-terminal sequence context library expressed with wt PylRS (**a**), PylRS($\beta^3$-1) (**c**), PylRS($\beta^2$-1) (**e**), PylRS(CA-1) (**g**), or PylRS($\alpha\alpha$-1) (**i**) in the absence (left) or presence (right) of **1** (**a**), **2** (**c**), **3** (**e**), **4** (**g**), or **5** (**i**). The fluorescence gate used to select permissive sequence contexts and the fraction of cells passing it are displayed in each graph. Experiments were performed in triplicate resulting in comparable results. The represented mean and standard deviation values were calculated with all replicates. Scatter plots represent GFP-fluorescence vs. a second channel (excitation at 405 nm, emission at 455).

**b,d,f,h,j,** Fluorescence values of 58 random sequence context reporters expressed with wt PylRS (**b**), PylRS($\beta^3$-1) (**d**), PylRS($\beta^2$-1) (**f**), PylRS(CA-1) (**h**), or PylRS($\alpha\alpha$-1) (**j**) in the presence of their respective amino acids were plotted against the incorporation efficiency score derived by FACS/NGS analysis. Sequences with a relative incorporation score higher than the lowest experimentally active variant (>1.2 fold change upon addition of ncM), and no-lower than the highest experimentally inactive variant , were considered as above the threshold (red dashed line). Individual incorporation efficiency threshold values for each ncM were determined to be 0.386 for ncM **1**, 0.125 for ncM **2**, 0.008 for ncM **3**, 0.216 for ncM **4** and 0.393 for ncM **5**. This threshold value was used to extrapolate the number of permissive sequence context across the entire sequence context library as measured by FACS/NGS. The distributions of incorporation efficiency scores for both samples are shown as histograms. The threshold indicated as a red dashed line.

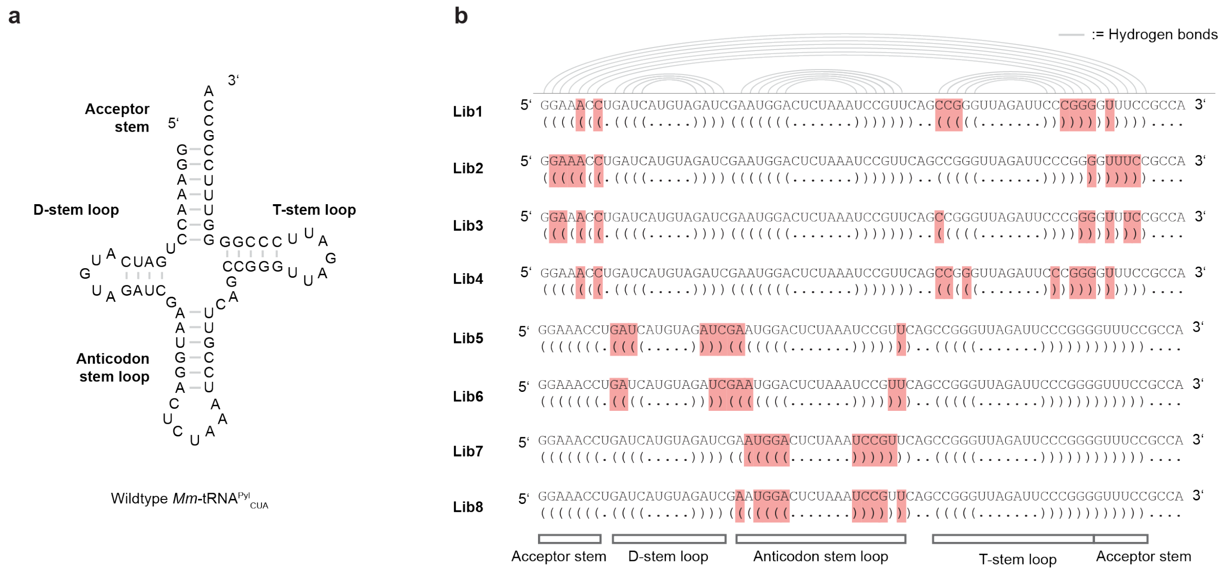


**Supplementary Figure 26. Eight libraries target different regions in tRNA$^{Pyl}$.**

**a**, Secondary structure of *Methanosarcina mazei* (*Mm*) tRNA$^{Pyl}_{CUA}$ (referred to as tRNA$^{Pyl}$).

**b**, tRNA library design. Positions targeted for mutagenesis in each of the six libraries are marked in red. Base pairs which form the secondary structure are shown by connecting grey lines between the involved nucleotides.

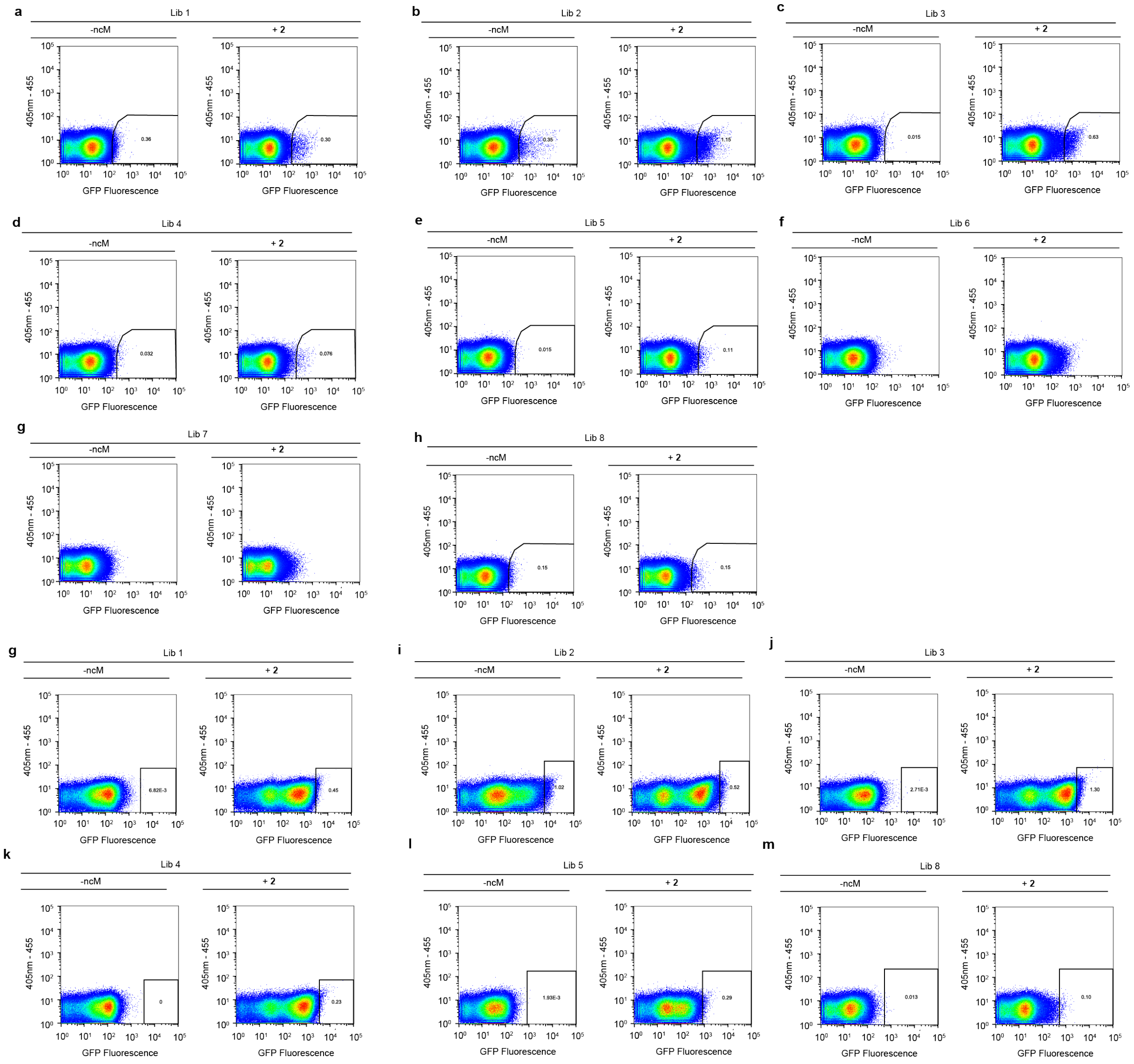


**Supplementary Figure 27. FACS-based tRNA selections for a β$^{3}$-amino acid.**

**a-h**, FACS of libraries 1-8 expressed in the presence and absence of **2**. Scatter plots represent GFP-fluorescence vs. a second channel (excitation at 405 nm, emission at 455 nm). We observed populations of highly fluorescent cells in the presence of **2** for libraries 1, 2, 3 and 6. For library 2 we also observed a large population of highly fluorescent cells in the absence of **2**, suggesting a population of non-orthogonal tRNAs in the library.

**g-m**, Second round of FACS of libraries of preselected libraries 1-5 and 8 expressed in the presence and absence of **2**. Scatter plots represent GFP-fluorescence vs. a second channel (excitation at 405 nm, emission at 455 nm). Libraries 1 and 3 were sorted according to the shown gates and further analysed by NGS. Experiments were performed in triplicate resulting in comparable results.

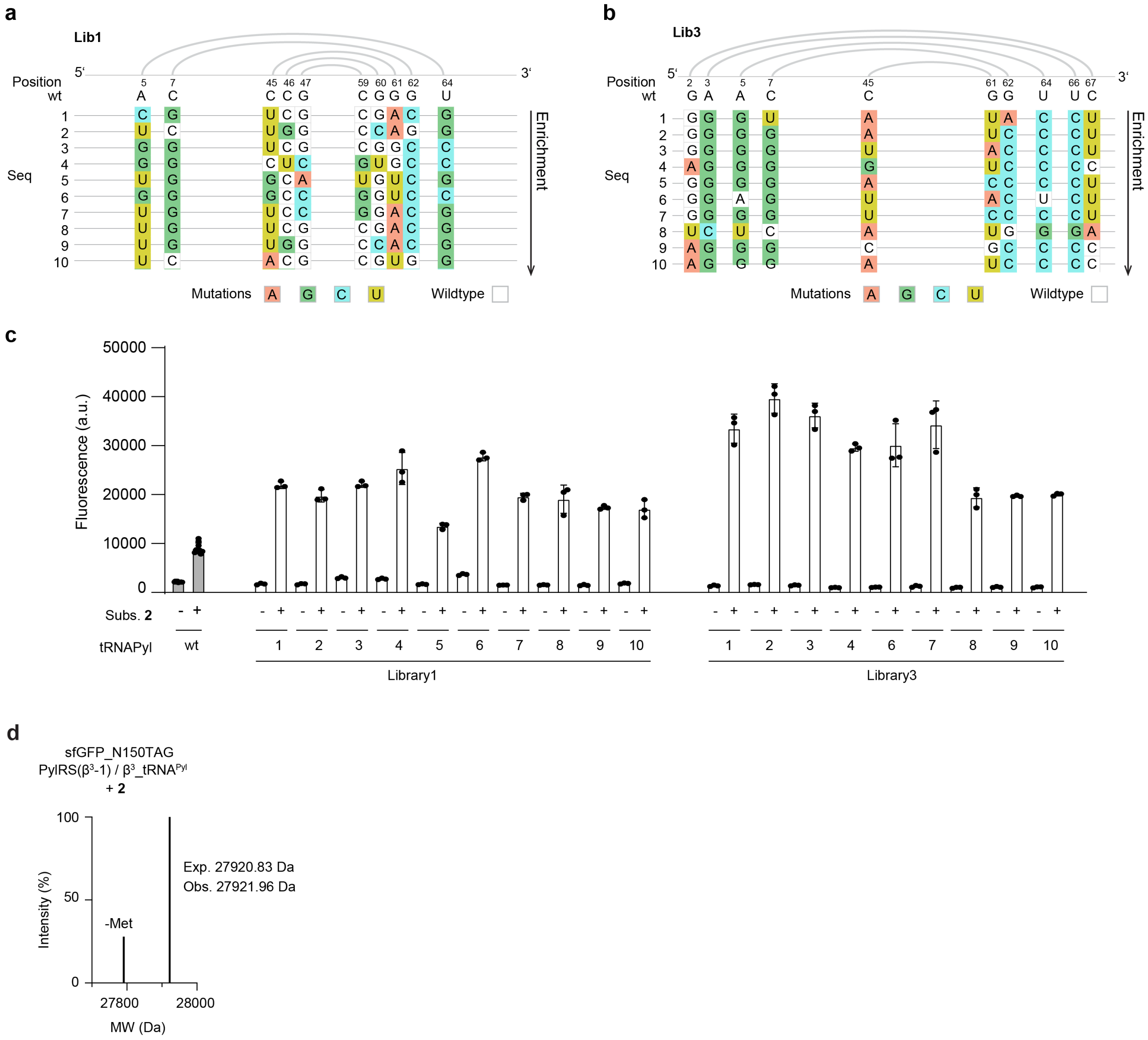


**Supplementary Figure 28. Selection and characterisation of library 1 and library 3 tRNAs for a $\beta^3$-amino acid.**

**a, b**, Top ten most enriched hits from library 1 (**a**) and library 3 (**b**) evolution for substrate **2**. Only the targeted positions are shown. Mutated positions are coloured. The secondary structure topological pairing of the targeted residues is shown by connecting grey lines.

**c,** GFP fluorescence of individually tested sequences resulting from evolutions with library 1 (left) and library 3 (right). Cells were transformed with a sfGFP(150TAG)$His_6$ reporter and a plasmid encoding for PylRS($\beta^3$-1) and the tRNA sequence. Protein expression was induced in the presence and absence of substrate **2**. GFP fluorescence levels are shown relative to those from cells containing *sfGFP(150TAG)*$His_6$, $tRNA^{Pyl}$ and wt PylRS expressed in the presence of 2 mM of **1** (set to 1).

**d**, Deconvoluted mass spectrum of sfGFP(150-**2**)$His_6$ purified from cells harbouring PylRS($\beta^3$-1) and *sfGFP(150TAG)*$His_6$ grown with **2**.

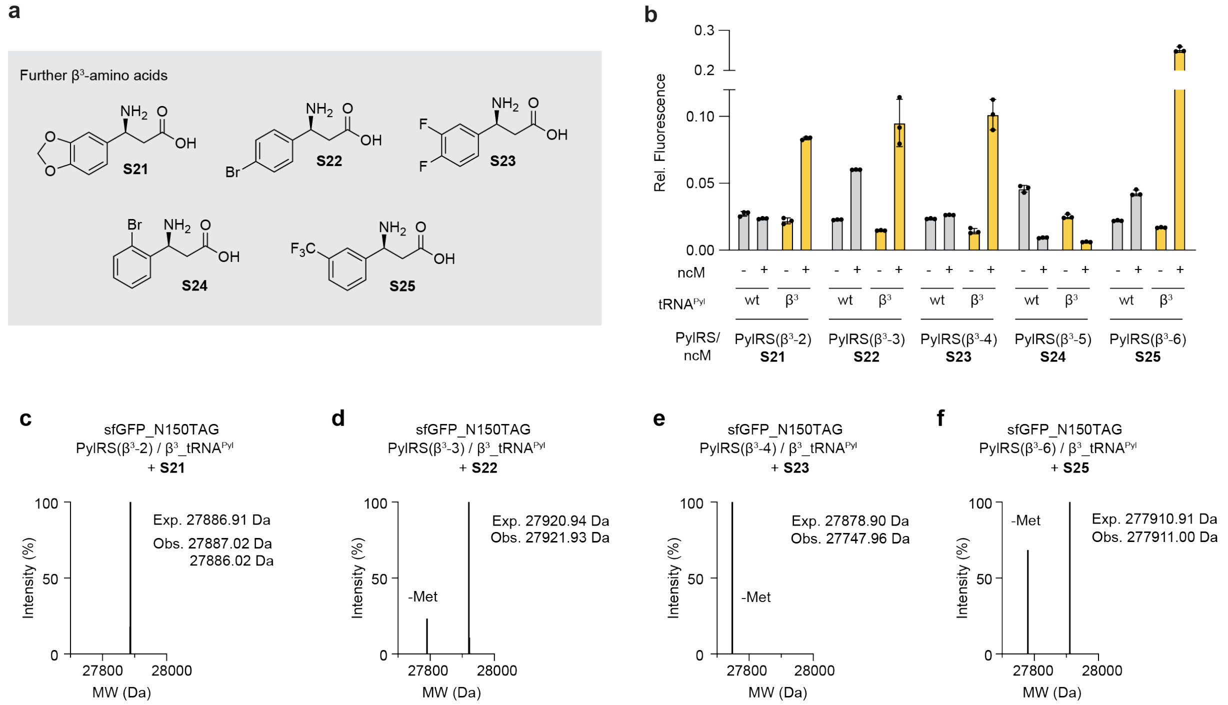


**Supplementary Figure 29. tRNA evolution enables incorporation of previously inaccessible β$^3$-amino acids.**

**a**, Structures of further β$^3$-amino acids **S21-S25**.

**b**, GFP fluorescence of cells transformed with a sfGFP(150TAG)His$_6$ reporter and a plasmid encoding for the indicated PylRS variant and either a wt tRNA$^{Pyl}$ or β$^3$_tRNA$^{Pyl}$. Cells were grown in the absence or presence of the respective ncM **S21-S25**. Fluorescence was normalised to expression of sfGFP(150TAG)His$_6$ with wt PylRS in the presence of **1** (2 mM).

**c-f**, Intact ESI-MS of sfGFP containing ncMs **S21**, **S22**, **S23**, and **S25** respectively at position 150. Cells contained a plasmid encoding for the respective PylRS variant as well as the engineered β$^3$_tRNA$^{Pyl}$ and expressed in the presence of each respective ncM. The observed mass corresponded to quantitative incorporation of the correct ncM in all cases.

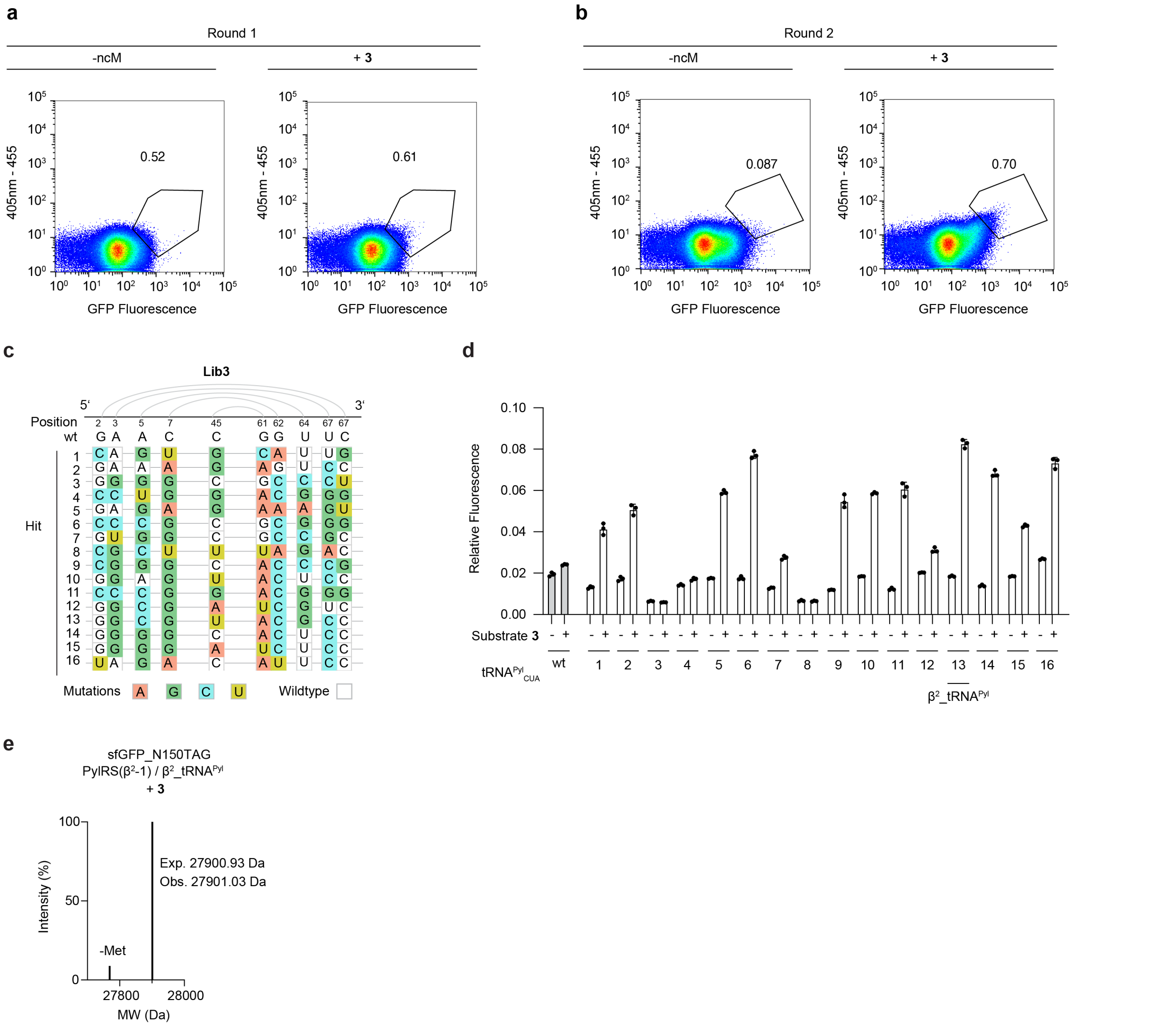

**Supplementary Figure 30. FACS-based tRNA selection for a β²-amino acid.**

**a-b**, Two-round FACS of library 3 expressed in the presence and absence of **3**. Scatter plots represent GFP-fluorescence vs. a second channel (excitation at 405 nm, emission at 455 nm).

**c**, Top 16 most enriched hits from library 3 evolution for substrate **3**. Only the targeted positions are shown. Mutated positions are coloured. The secondary structure topological pairing of the targeted residues is shown by connecting grey lines.

**d**, GFP fluorescence of individually tested sequences resulting from evolutions with library 3. Cells were transformed with a sfGFP(150TAG)His$_6$ reporter and a plasmid encoding for PylRS(β$^2$-1) and the tRNA sequence. Protein expression was induced in the presence and absence of substrate **3**.

**e**, Deconvoluted mass spectrum of sfGFP(150-**3**)$His_6$ purified from cells harbouring PylRS($\beta^2$-1) and *sfGFP(150TAG)His*$_6$ grown with **3**.

.

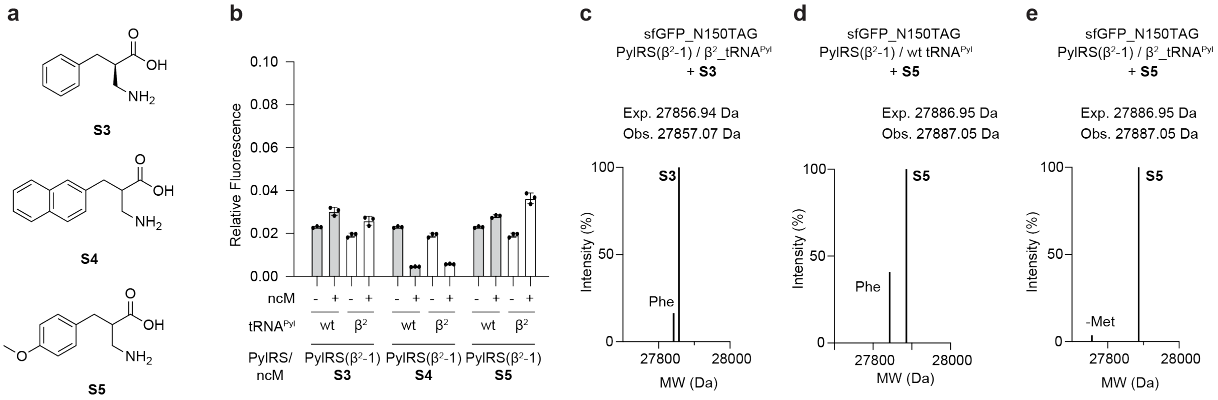


**Supplementary Figure 31. Increased incorporation of further $\beta^2$-amino acids through engineered tRNA.**

**a**, Structures of different $\beta^2$-amino acids **S3**, **S4** and **S5**.

**b**, GFP fluorescence of cells transformed with a sfGFP(150TAG)$His_6$ reporter and a plasmid encoding for PylRS($\beta^2$-1) and either wt $tRNA^{Pyl}$ or $\beta^2$_$tRNA^{Pyl}$. Cells were expressed with and without the respective ncMs (**S3-S5**). Fluorescence was normalised to expression of sfGFP(150TAG) with wt PylRS in the presence of **1** (2 mM).

**c**,**d**,**e** Intact ESI-MS of sfGFP containing ncMs **S3** (**c**) and **S5** (**d** and **e**) respectively at position 150. Cells contained a plasmid encoding for PylRS($\beta^2$-1) as well as the engineered $\beta^2$_$tRNA^{Pyl}$ (**c** and **e**) or wt $tRNA^{Pyl}$ (**d**) and expressed in the presence of each respective ncM. Incorporation of **S3** is observed with $\beta^2$_$tRNA^{Pyl}$, with a minor peak corresponding to phenylalanine misincorporation. Data in (**d**) is duplicated from **Supplementary Fig. 22d**. $\beta^2$_$tRNA^{Pyl}$ allows quantitative incorporation in the presence of **S5**.

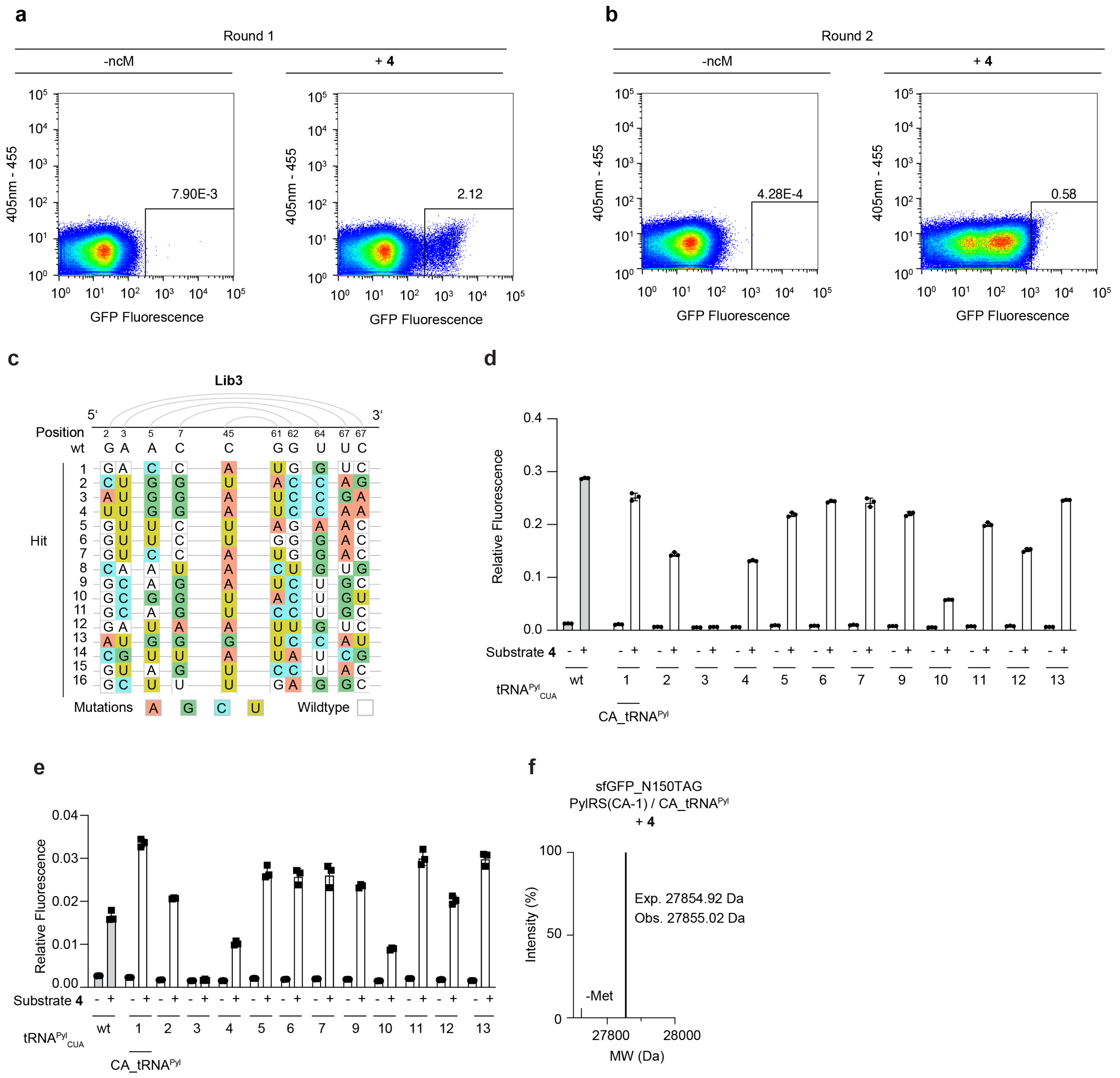


**Supplementary Figure 32. FACS-based tRNA selection for an N-cyclic amino acid.**

**a-b**, Two-round FACS of library 3 expressed in the presence and absence of **4**. Scatter plots represent GFP-fluorescence vs. a second channel (excitation at 405 nm, emission at 455 nm).

**c**, Top 16 most enriched hits from an evolution for substrate **4** run with tRNA library 3. Only the targeted positions are shown. Mutated positions are coloured. The secondary structure topological pairing of the targeted residues is shown by connecting grey lines.

**d**, GFP fluorescence of individually tested sequences resulting from evolutions with library 3. Cells were transformed with a sfGFP(150TAG)$His_6$ reporter and a plasmid encoding for PylRS(CA-1) and the tRNA sequence. Protein expression was induced in the presence and absence of substrate **4**.

Fluorescence was normalised to expression of sfGFP(150TAG) with wt PylRS in the presence of **1** (2 mM).

**e**, GFP fluorescence of individually tested sequences resulting from evolutions with library 3. Cells were transformed with an N-terminal sfGFP reporter with a Cys-TAG-Pro context (TGC-TAG-CCA) and a plasmid encoding for PylRS(CA-1) and the tRNA sequence. Protein expression was induced in the presence and absence of substrate **4**. Cells grown in the presence of CA-$tRNA^{Pyl}$ showed the highest level of monomer-dependent GFP fluorescence. Fluorescence was normalised to expression of sfGFP(150TAG) with wt PylRS in the presence of **1** (2 mM).

**f**, Deconvoluted mass spectrum of sfGFP(150-**4**)$His_6$ purified from cells harbouring PylRS(CA-1) and *sfGFP(150TAG)*$His_6$ grown with **4**.

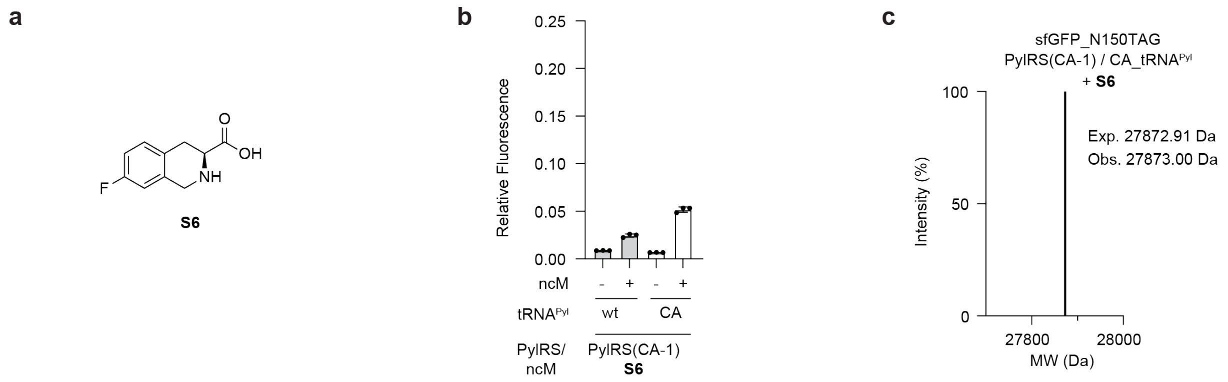


**Supplementary Figure 33. Increased incorporation of a further N-cyclic amino acid through engineered tRNA.**

**a**, Structure of ncM **S6**. **b**, GFP fluorescence of cells transformed with a sfGFP(150TAG)$His_6$ reporter and a plasmid encoding for PylRS(CA-1) and either wt $tRNA^{Pyl}$ or $CA\_tRNA^{Pyl}$. Expression was conducted with and without substrate **S6**. Fluorescence was normalised to expression of sfGFP(150TAG) with wt PylRS in the presence of **1** (2 mM). **c**, Intact ESI-MS of sfGFP containing ncM **S6** at position 150. Cells contained a plasmid encoding for the respective PylRS variant as well as the engineered $CA\_tRNA^{Pyl}$ and protein was expressed in the presence of **S6**. The observed mass corresponded to quantitative incorporation of the correct ncM.

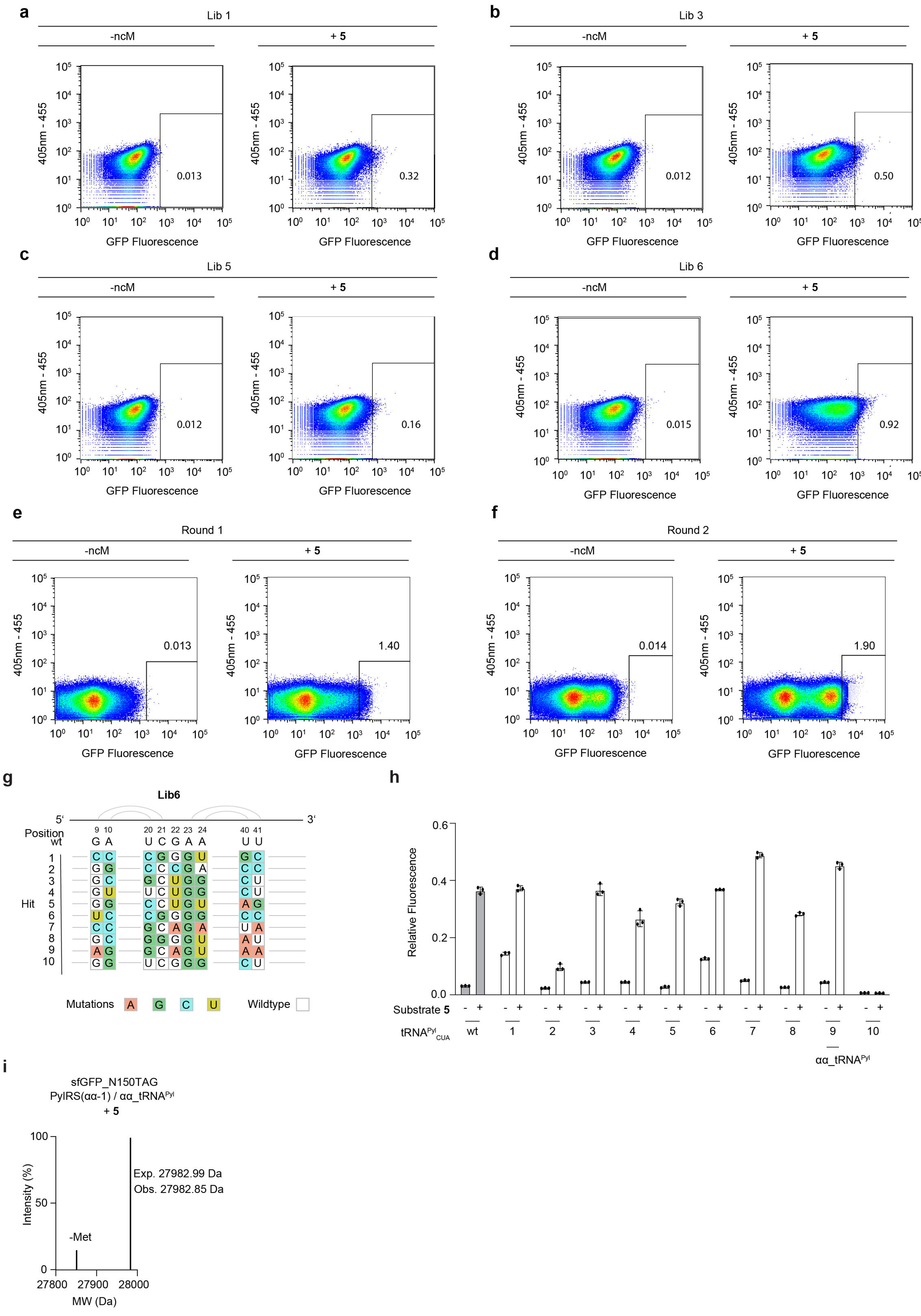

**Supplementary Figure 34. FACS-based tRNA selection for substrate** 5.

**a-d**, Flow-cytometry-based screening of four different libraries for suitability of tRNA evolution for substrate **5**. Libraries 1 (**a**), 3 (**b**), 8 (**c**), and 6 (**d**) were expressed in the presence and absence of **5**.

Scatter plots represent GFP-fluorescence vs. a second channel (excitation at 405 nm, emission at 455 nm).

**e-f**, Two-round FACS of library 6 expressed in the presence and absence of **5**. Scatter plots represent GFP-fluorescence vs. a second channel (excitation at 405 nm, emission at 455 nm).

**g**, Top 10 most enriched hits from library 6 evolution for substrate **5**. Only the targeted positions are shown. Mutated positions are coloured. The secondary structure topological pairing of the targeted residues is shown by connecting grey lines.

**h**, GFP fluorescence of individually tested sequences resulting from evolutions with library 6. Cells were transformed with a sfGFP(150TAG)$His_6$ reporter and a plasmid encoding for PylRS(αα-1) and the tRNA sequence. Protein expression was induced in the presence and absence of substrate **5**. Fluorescence was normalised to expression of sfGFP(150TAG) with wt PylRS in the presence of **1** (2 mM).

**i**, Deconvoluted mass spectrum of sfGFP(150-**5**)$His_6$ purified from cells harbouring PylRS(αα-1) and *sfGFP(150TAG)His*$_6$ grown with **5**.

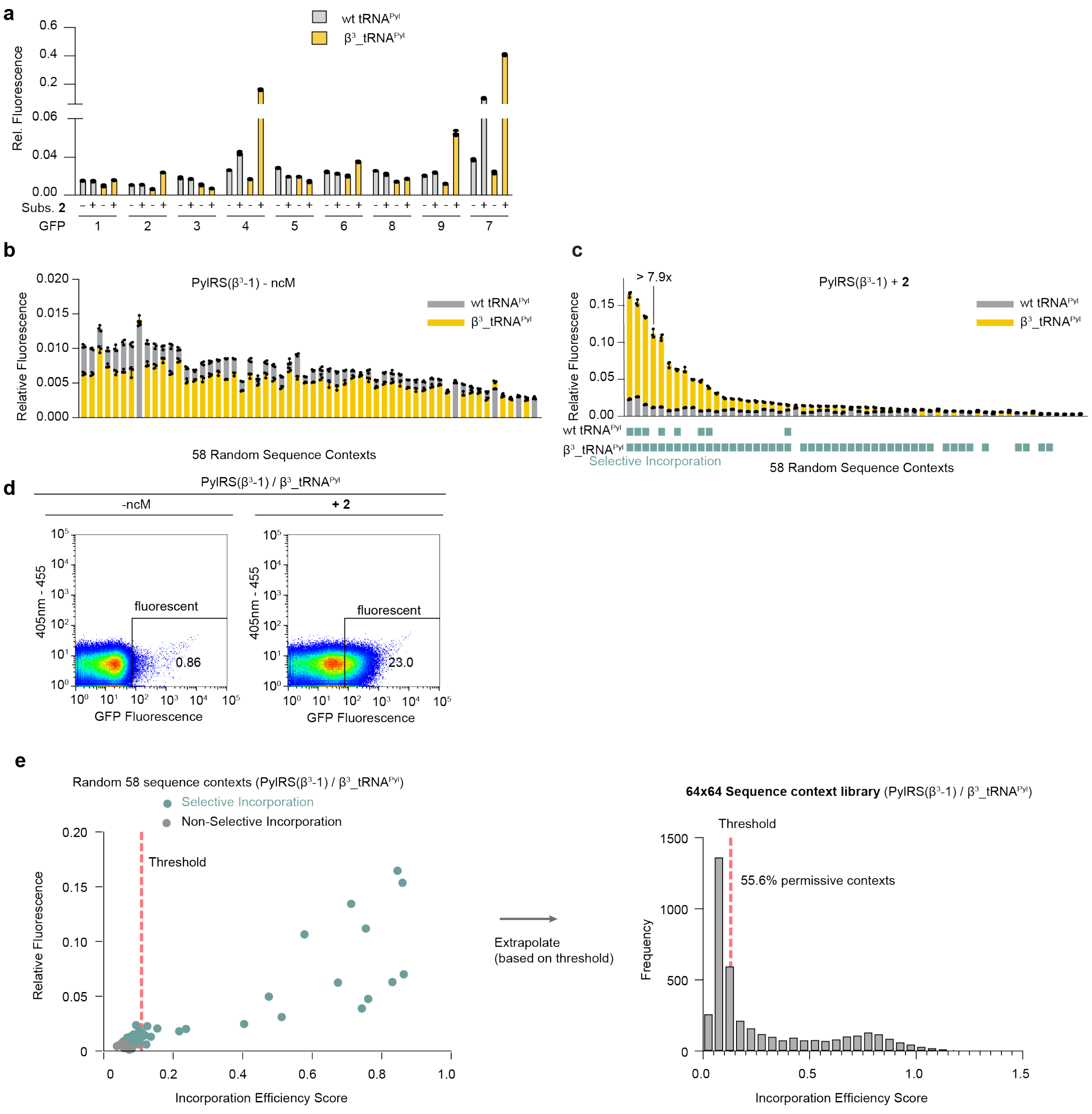


**Supplementary Figure 35. FACS analysis of sequence context libraries for β$^3$_tRNA$^{Pyl}$.**

**a** GFP fluorescence from cells transformed with sfGFP reporters containing a TAG codon at 9 different positions throughout the protein. Expression was performed with cells bearing a plasmid encoding for PylRS(β$^3$-1) and wt tRNA$^{Pyl}$ or β$^3$_tRNA$^{Pyl}$ in the presence and absence of ncM **2**. The positions of the suppressed codon in the sfGFP reporters 1-9 are: 39, 44, 50, 98, 109, 116, 150, 154, and 187.

**b**,**c**, Fluorescence values of 58 random sequence context reporters expressed with PylRS($\beta^3$-1) and either wt $tRNA^{Pyl}$ or $\beta^3$_$tRNA^{Pyl}$ pair in the absence (**a**) or presence (**b**) of **2**. Fluorescence was normalised to expression of sfGFP(150TAG) with wt PylRS in the presence of **1** (2 mM).

**d**, Fluorescence distribution measured by FACS of N-terminal sequence context library expressed with the PylRS($\beta^3$-1)/ $\beta^3$_$tRNA^{Pyl}$ pair in the absence (left) and presence (right) of **2**.

**e**, Fluorescence values of 58 random sequence context reporters expressed with the PylRS($\beta^3$-1)/$\beta^3$_$tRNA^{Pyl}$ pair in the presence of **2** were plotted against the incorporation efficiency score derived by FACS/NGS analysis. Sequences with a relative incorporation score higher than the lowest experimentally active (>1.2 fold change upon addition of ncM) variant were considered to support selective incorporation of **2**. We set a threshold value for the incorporation efficiency score at the lowest context supporting selective incorporation, but above the highest non-selective incorporation at a sequence context (red dashed line). This threshold value (0.109) was used to extrapolate the number of permissive sequence context across the entire sequence context library as measured by FACS/NGS. The distributions of incorporation efficiency scores for both samples are shown as histograms. The threshold derived is indicated as a red dashed line.

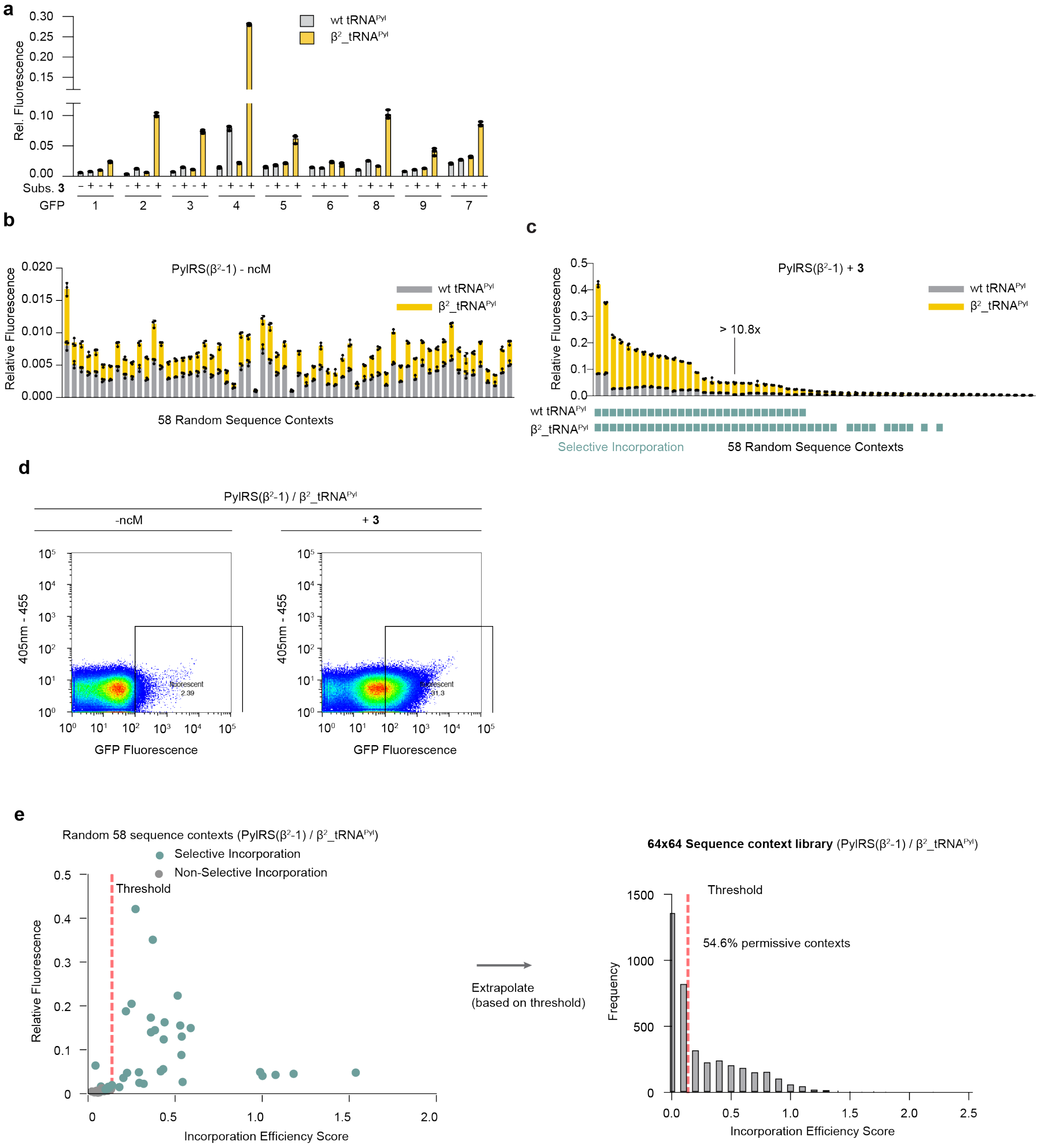


**Supplementary Figure 36. FACS analysis of sequence context libraries for β$^2$_tRNA$^{Pyl}$.**

**a,** GFP fluorescence from cells transformed with sfGFP reporters containing a TAG codon at 9 different positions throughout the protein. Expression was performed with cells bearing a plasmid encoding for PylRS(β$^2$-1) and wt tRNA$^{Pyl}$ or β$^2$_tRNA$^{Pyl}$ in the presence and absence of ncM **3**. The positions of the suppressed codon in the sfGFP reporters 1-9 are: 39, 44, 50, 98, 109, 116, 150, 154, and 187.

**b**,**c**, Fluorescence values of 58 random sequence context reporters expressed with PylRS($\beta^2$-1)/ ) and either wt tRNA$^{Pyl}$ or $\beta^2$_tRNA$^{Pyl}$ pair in the absence (**a**) or presence (**b**) of **3**. Fluorescence was normalised to expression of sfGFP(150TAG) with wt PylRS in the presence of **1** (2 mM).

**d**, Fluorescence distribution measured by FACS of N-terminal sequence context library expressed with the PylRS($\beta^2$-1)/ $\beta^2$_tRNA$^{Pyl}$ pair in the absence (left) and presence (right) of **3**.

**e**, Fluorescence values of 58 random sequence context reporters expressed with the PylRS($\beta^2$-1)/$\beta^2$_tRNA$^{Pyl}$ pair in the presence of **3** were plotted against the incorporation efficiency score derived by FACS/NGS analysis. Sequences with a relative incorporation score higher than the lowest experimentally active (>1.2 fold change upon addition of ncM) variant were considered to support selective incorporation of **3**. We set a threshold value for the incorporation efficiency score at the lowest context supporting selective incorporation, but above the highest non-selective incorporation at a sequence context (red dashed line). This threshold value (0.137) was used to extrapolate the number of permissive sequence context across the entire sequence context library as measured by FACS/NGS. The distributions of incorporation efficiency scores for both samples are shown as histograms. The threshold derived is indicated as a red dashed line.

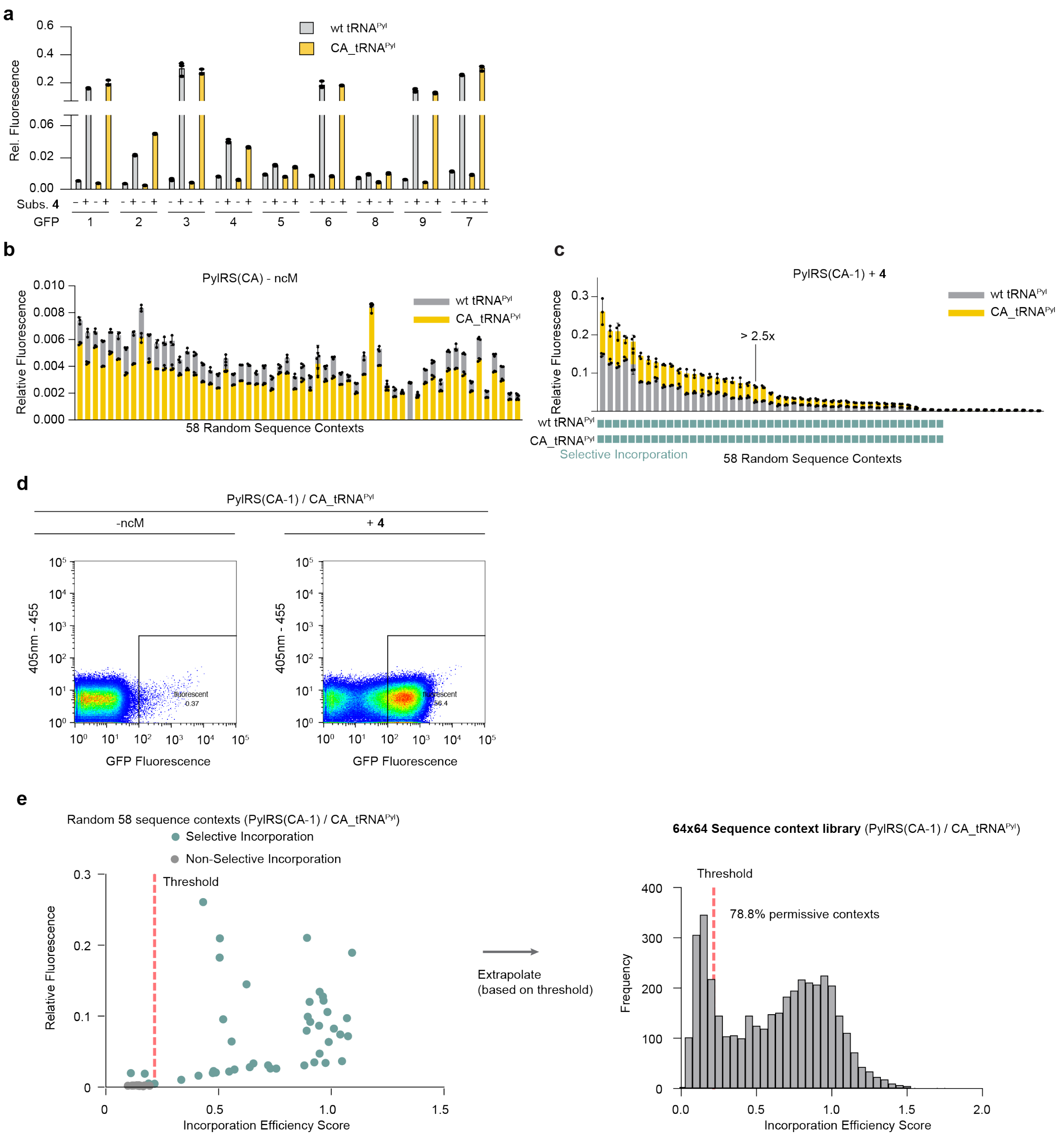


**Supplementary Figure 37. FACS analysis of sequence context libraries for CA_tRNA$^{Pyl}$.**

**a,** GFP fluorescence from cells transformed with sfGFP reporters containing a TAG codon at 9 different positions throughout the protein. Expression was performed with cells bearing a plasmid encoding for PylRS(CA-1) and wt tRNA$^{Pyl}$ or CA_tRNA$^{Pyl}$ in the presence and absence of ncM **4**. The positions of the suppressed codon in the sfGFP reporters 1-9 are: 39, 44, 50, 98, 109, 116, 150, 154, and 187.

**b**,**c**, Fluorescence values of 58 random sequence context reporters expressed with PylRS(CA-1) ) and either wt $tRNA^{Pyl}$ or CA_$tRNA^{Pyl}$ pair in the absence (**a**) or presence (**b**) of **4**. Fluorescence was normalised to expression of sfGFP(150TAG) with wt PylRS in the presence of **1** (2 mM).

**d**, Fluorescence distribution measured by FACS of N-terminal sequence context library expressed with the PylRS(CA-1)/ CA_$tRNA^{Pyl}$ pair in the absence (left) and presence (right) of **4**.

**e**, Fluorescence values of 58 random sequence context reporters expressed with the PylRS(CA-1)/CA_$tRNA^{Pyl}$ pair in the presence of **4** were plotted against the incorporation efficiency score derived by FACS/NGS analysis. Sequences with a relative incorporation score higher than the lowest experimentally active (>1.2 fold change upon addition of ncM) variant were considered to support selective incorporation of **4**. We set a threshold value for the incorporation efficiency score at the lowest context supporting selective incorporation, but above the highest non-selective incorporation at a sequence context (red dashed line). This threshold value (0.216) was used to extrapolate the number of permissive sequence context across the entire sequence context library as measured by FACS/NGS. The distributions of incorporation efficiency scores for both samples are shown as histograms. The threshold derived is indicated as a red dashed line.

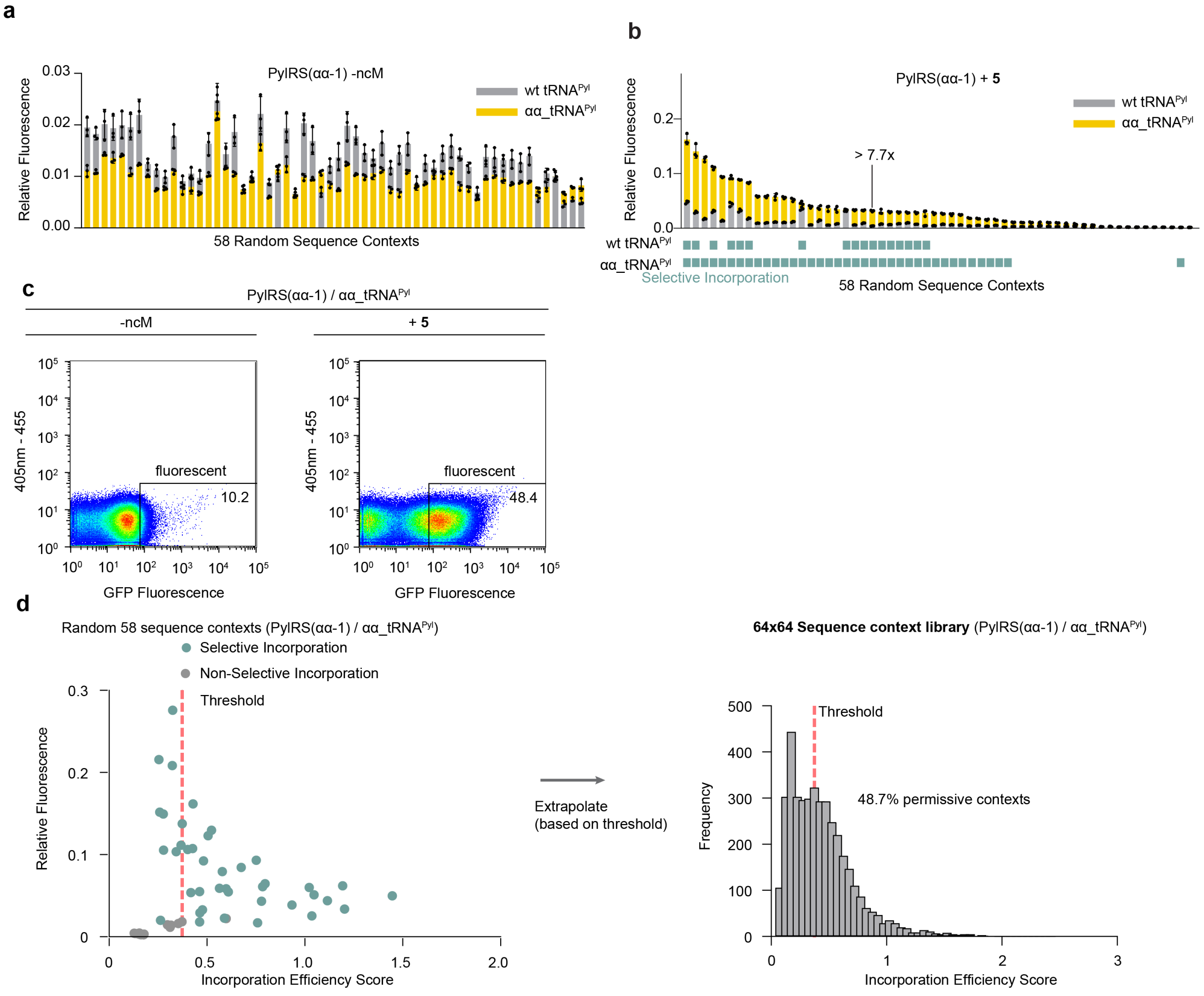


**Supplementary Figure 38. FACS analysis of sequence context libraries for αα_tRNA$^{Pyl}$.**

**a,** GFP fluorescence from cells transformed with sfGFP reporters containing a TAG codon at 9 different positions throughout the protein. Expression was performed with cells bearing a plasmid encoding for PylRS(αα-1) and wt tRNA$^{Pyl}$ or αα_tRNA$^{Pyl}$ in the presence and absence of ncM **5**. The positions of the suppressed codon in the sfGFP reporters 1-9 are: 39, 44, 50, 98, 109, 116, 150, 154, and 187.

**b**,**c**, Fluorescence values of 58 random sequence context reporters expressed with PylRS(αα-1) and either wt tRNA$^{Pyl}$ or αα_tRNA$^{Pyl}$ pair in the absence (**a**) or presence (**b**) of **5**. Fluorescence was normalised to expression of sfGFP(150TAG) with wt PylRS in the presence of **1** (2 mM).

**d**, Fluorescence distribution measured by FACS of N-terminal sequence context library expressed with the PylRS(αα-1)/αα_tRNA$^{Pyl}$ pair in the absence (left) and presence (right) of **5**.

**e**, Fluorescence values of 58 random sequence context reporters expressed with the PylRS(αα-1)/ aa _tRNA$^{Pyl}$ pair in the presence of **5** were plotted against the incorporation efficiency score derived by FACS/NGS analysis. Sequences with a relative incorporation score higher than the lowest experimentally active (>1.2 fold change upon addition of ncM) variant were considered to support selective incorporation of **5**. We set a threshold value for the incorporation efficiency score at the lowest context supporting selective incorporation, but above the highest non-selective incorporation at a sequence context (red dashed line). This threshold value (0.401) was used to extrapolate the number of permissive sequence context across the entire sequence context library as measured by FACS/NGS. The distributions of incorporation efficiency scores for both samples are shown as histograms. The threshold derived is indicated as a red dashed line.

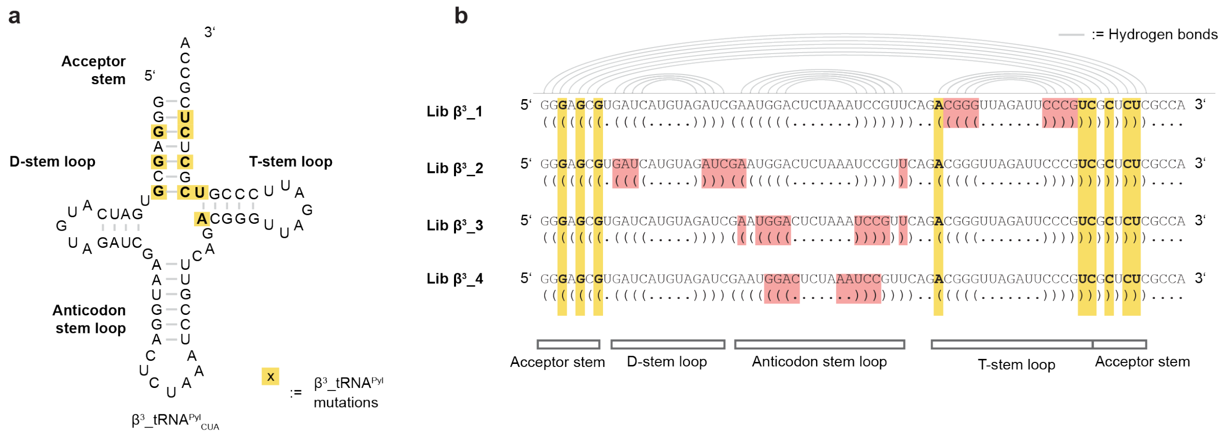


**Supplementary Figure 39. Four libraries target different regions in β$^3$_tRNA$^{Pyl}$.**

**a**, Secondary structure of β$^3$_tRNA$^{Pyl}$ based on *Methanosarcina mazei* (*Mm*) tRNA$^{Pyl}_{CUA}$.

**b**, tRNA library design. Secondary structure representation (**a**) and sequence (**b**) of β$^3$_tRNA$^{Pyl}$. Positions targeted for mutagenesis in each of the four libraries are marked in red. Base pairs which form the secondary structure are shown by connecting grey lines between the involved nucleotides.

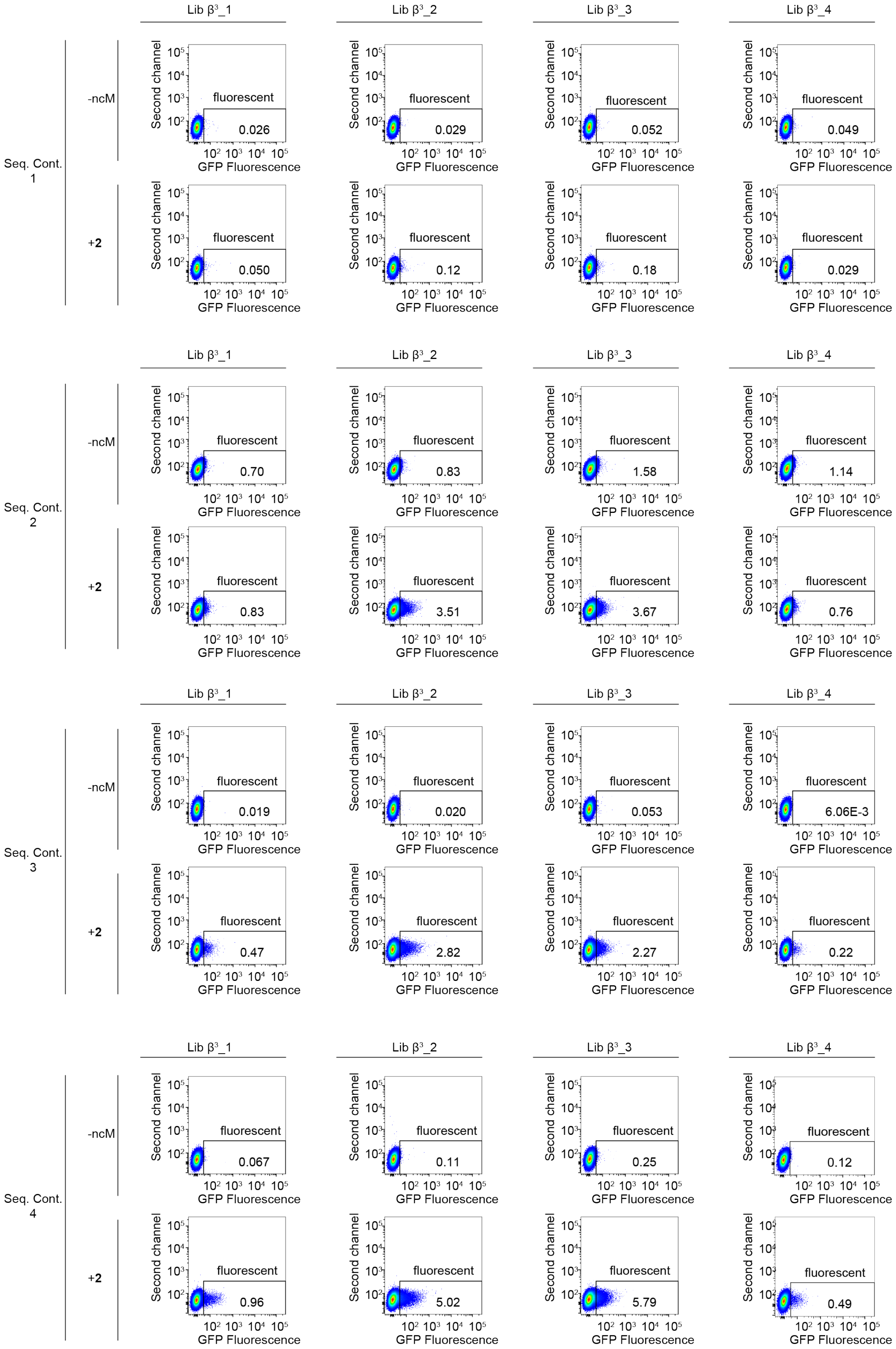

Lib β³_1
Lib β³_2
Lib β³_3
Lib β³_4
Seq. Cont. 1
-ncM
0.026
0.029
0.052
0.049
+2
0.050
0.12
0.18
0.029
Seq. Cont. 2
-ncM
0.70
0.83
1.58
1.14
+2
0.83
3.51
3.67
0.76
Seq. Cont. 3
-ncM
0.019
0.020
0.053
6.06E-3
+2
0.47
2.82
2.27
0.22
Seq. Cont. 4
-ncM
0.067
0.11
0.25
0.12
+2
0.96
5.02
5.79
0.49
fluorescent
Second channel
GFP Fluorescence
10²
10³
10⁴
10⁵

**Supplementary Figure 40. Screening of different tRNA libraries for a $\beta^3$-amino acid across different sequence context.**

**a-d**, Flow-cytometry-based screening of four different libraries for suitability of tRNA evolution for substrate **2**. Libraries $\beta^3$-1, $\beta^3$-2, $\beta^3$-3, and $\beta^3$-4 were expressed in the presence and absence of **2** with s.c.1-, s.c.2-, s.c.3, and s.c.4- reporters. Scatter plots represent GFP-fluorescence vs. a second channel (excitation at 405 nm, emission at 455 nm).

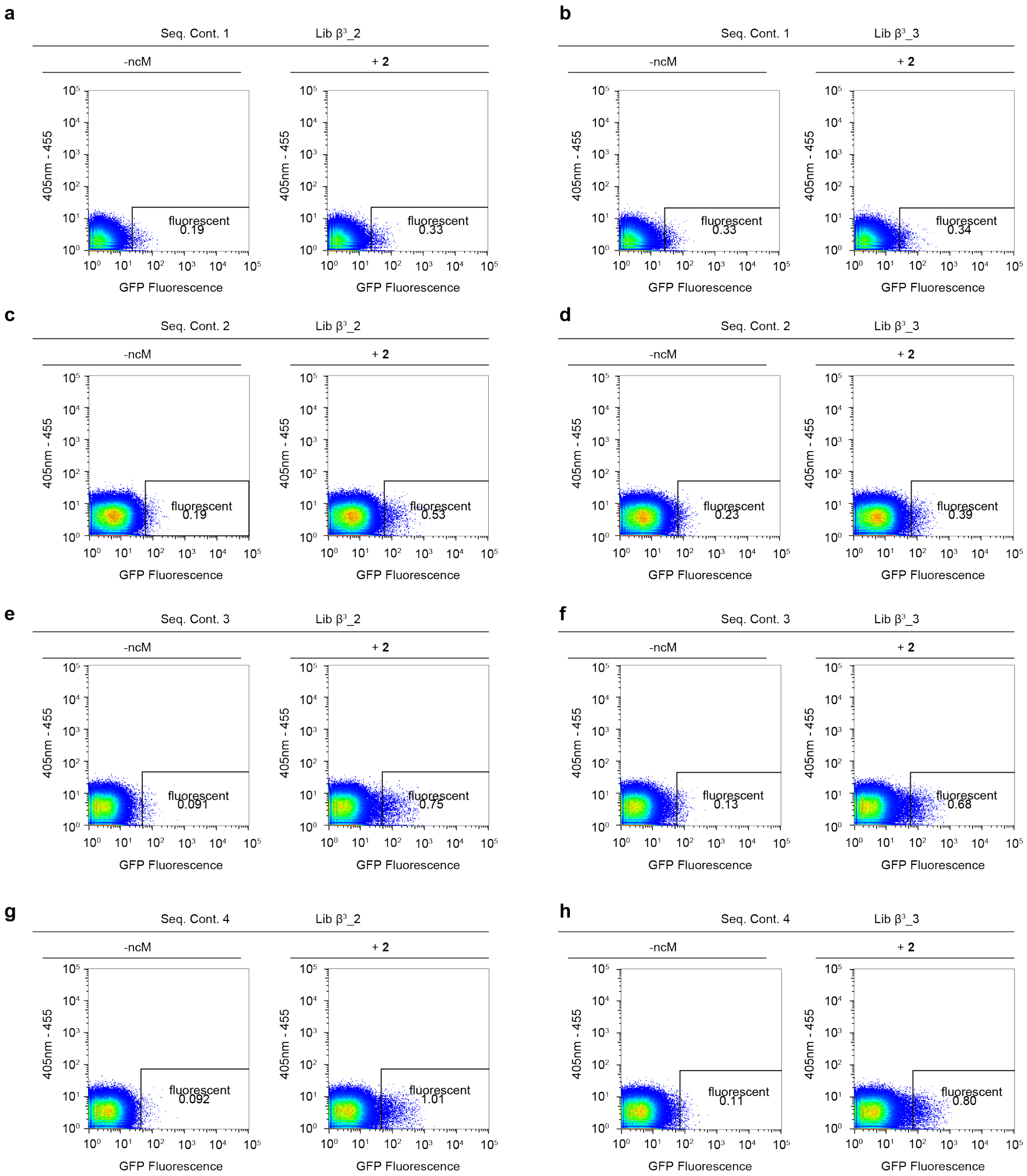


**Supplementary Figure 41. First round of FACS-based tRNA selections.**

**a-h**, FACS of libraries $\beta^3$-2 (**a**, **c**, **e**, **g**) and $\beta^3$-3 (**b**, **d**, **f**, **h**) expressed in the presence and absence of **2** with s.c.1- (**a**, **b**), s.c.2 (**c**, **d**), s.c.3 (**e**, **f**), and s.c.4 (**g**, **h**) reporters. Scatter plots represent GFP-fluorescence vs. a second channel (excitation at 405 nm, emission at 455 nm).

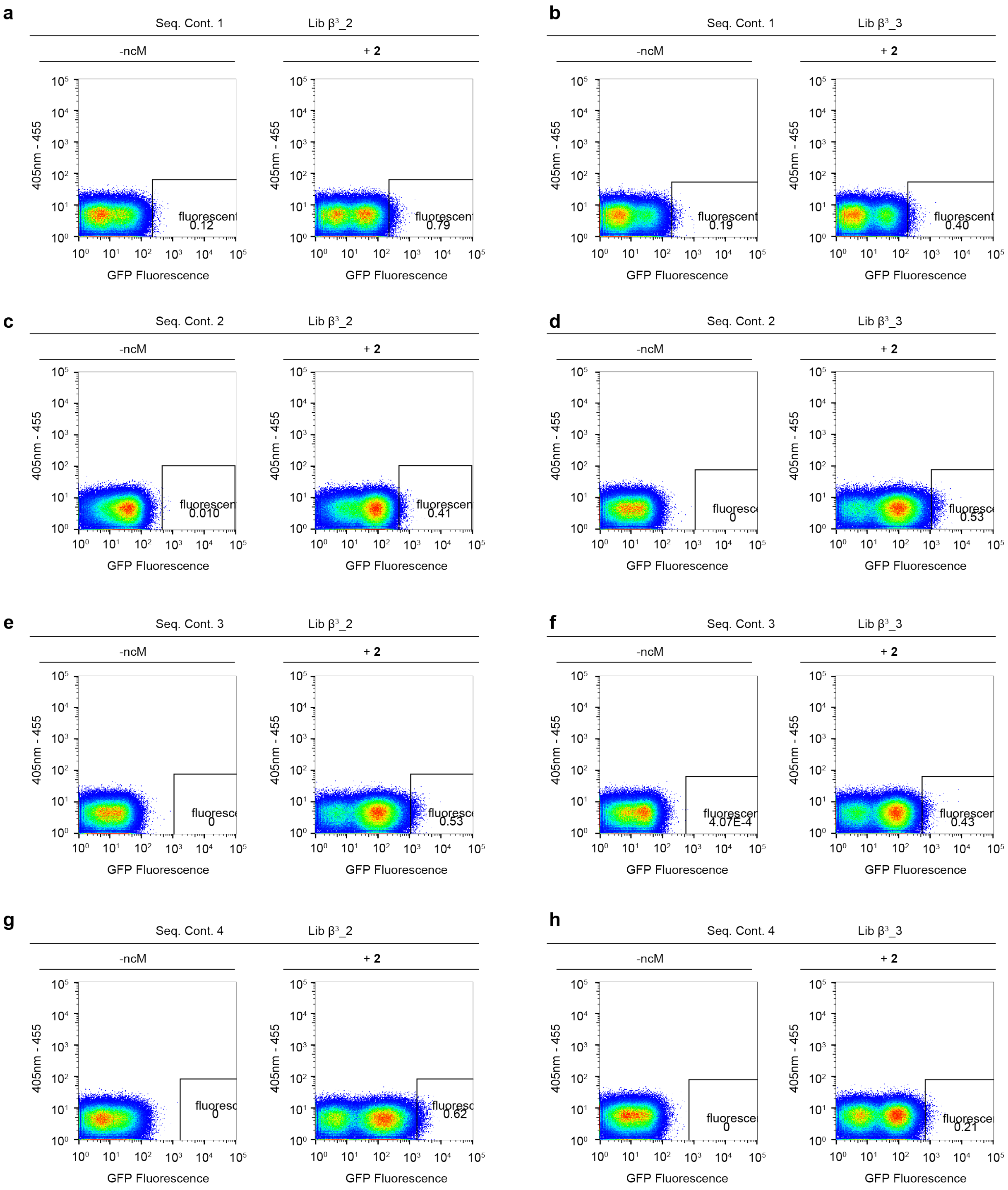


**Supplementary Figure 42. Second round of FACS-based tRNA selections.**

**a-h**, FACS of libraries $\beta^3$-2 (**a**, **c**, **e**, **g**) and $\beta^3$-3 (**b**, **d**, **f**, **h**) expressed in the presence and absence of **2** with s.c.1 (**a**, **b**), s.c.2 (**c**, **d**), s.c.3 (**e**, **f**), and s.c.4 (**g**, **h**) reporters. Scatter plots represent GFP-fluorescence vs. a second channel (excitation at 405 nm, emission at 455 nm).

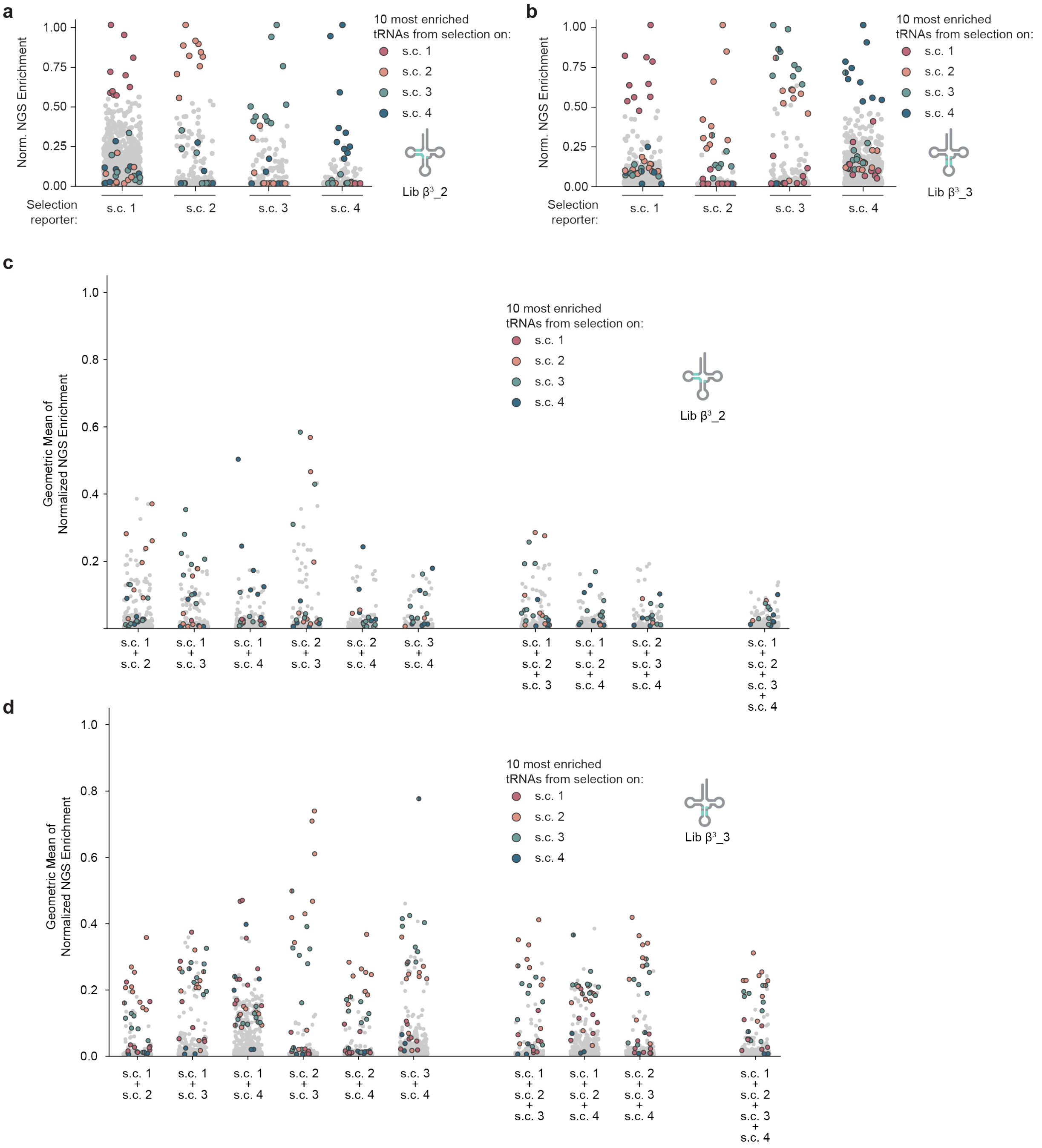


**Supplementary Figure 43. NGS overlap of FACS-based tRNA selections.**

**a,b**, Normalised sequencing enrichment values for two-round tRNA selections performed with library $\beta^3$_2 targeting the D-arm (**a**) or library $\beta^3$_3 targeting the anticodon stem (**b**) using s.c.1, s.c.2, s.c.3 and s.c.4 reporters. The enrichment of each tRNA variant is normalised to the largest value in each selection. The top ten enriched sequences of each selection are highlighted in colour across all four parallel selections.

**c,d**, NGS overlap of sequences of library $\beta^3$-2 (**c**) or $\beta^3$-3 (**d**) selected on different sequence contexts. The geometric mean of the normalised enrichment of each sequence across all combinations of selected sequence contexts is shown. The top 10 enriched hits from each individual selection are highlighted.

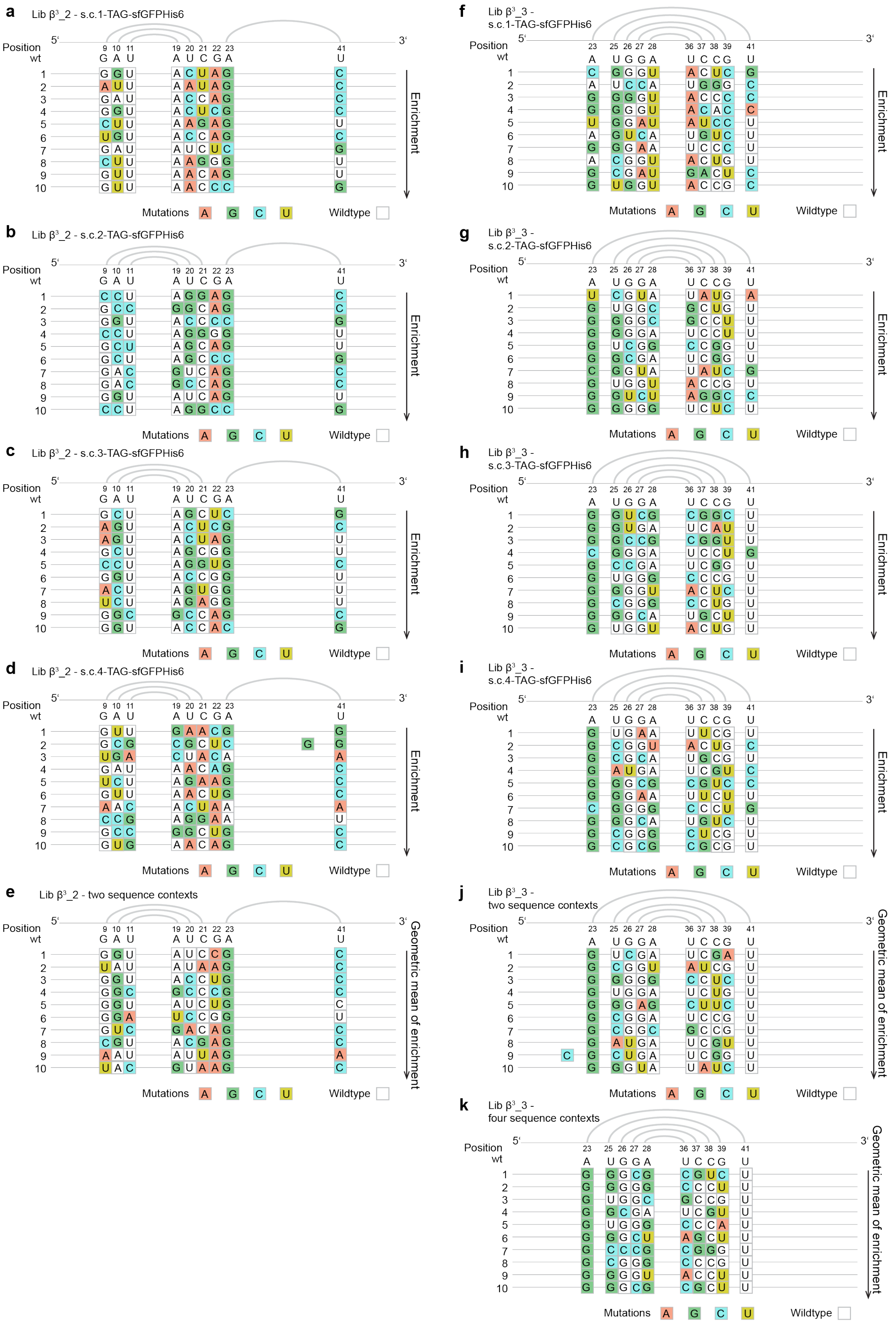
a Lib β³_2 - s.c.1-TAG-sfGFPHis6
b Lib β³_2 - s.c.2-TAG-sfGFPHis6
c Lib β³_2 - s.c.3-TAG-sfGFPHis6
d Lib β³_2 - s.c.4-TAG-sfGFPHis6
e Lib β³_2 - two sequence contexts
f Lib β³_3 - s.c.1-TAG-sfGFPHis6
g Lib β³_3 - s.c.2-TAG-sfGFPHis6
h Lib β³_3 - s.c.3-TAG-sfGFPHis6
i Lib β³_3 - s.c.4-TAG-sfGFPHis6
j Lib β³_3 - two sequence contexts
k Lib β³_3 - four sequence contexts
Position
wt
5'
3'
Enrichment
Geometric mean of enrichment
Mutations A G C U
Wildtype

**Supplementary Figure 44. Sequences of tRNAs derived from sequence context tRNA selections for the β$^3$-amino acid 2.**

**a-e**, Top 10 most enriched hits from the evolution with library β$^3$_2 selected with reporters s.c.1 (**a**), s.c.2 (**b**), s.c.3-TAG (**c**), and s.c.4-TAG (**d**). Additionally, the 10 sequences with the highest geometric mean of enrichment across two sequence contexts were chosen for characterisation (**e**). Only the targeted positions are shown. Mutated positions are coloured. The secondary structure topological pairing of the targeted residues is shown by connecting grey lines.

**f-k**, Top 10 most enriched hits from the evolution with library β$^3$_3 selected with reporters s.c.1 (**f**), s.c.2 (**g**), s.c.3 (**h**), and s.c.4 (**i**). Additionally, the 10 sequences with the highest geometric mean of enrichment across two and four sequence contexts were chosen for characterisation (**j**, **k**). Only the targeted positions are shown. Mutated positions are coloured. The secondary structure topological pairing of the targeted residues is shown by connecting grey lines.

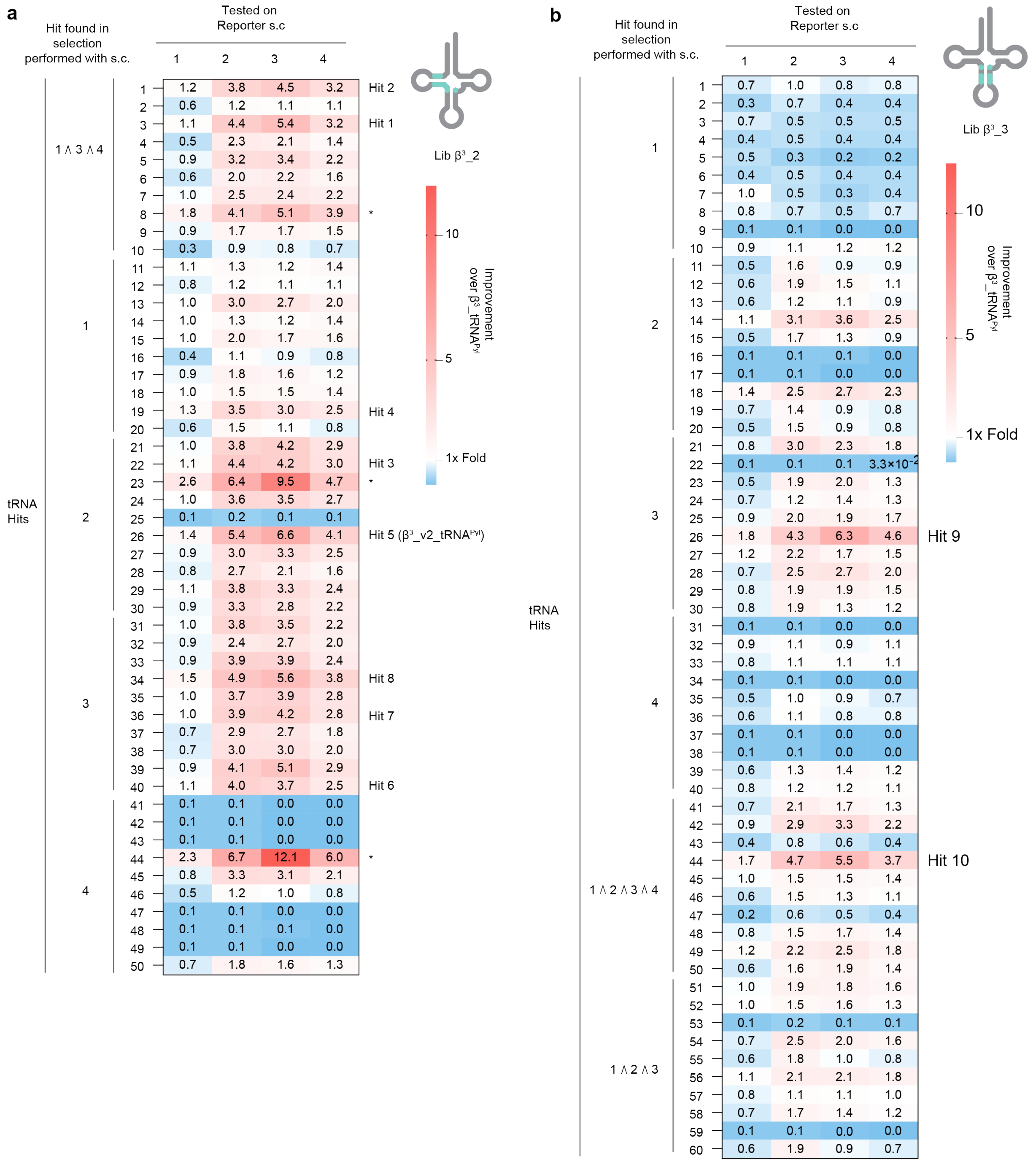


**Supplementary Figure 45. Screening of tRNAs derived from sequence context tRNA selections for β$^3$-amino acid 2.**

**a** and **b**, GFP fluorescence values of tRNA sequences resulting from parallel selections with library β$^3$_2 (**a**) or library β$^3$_3 (**b**) tested using s.c.1, s.c.2, s.c.3, and s.c.4 as reporters. Plasmids harbouring PylRS(β$^3$-1) and the corresponding tRNA$^{Pyl}$ hit were transformed into cells containing one of the reporters and expression was conducted in the presence of **2**. The tRNA sequences are arranged by the selection from which they were chosen. Sequences enriched in multiple selections were tested in addition to the top ten most enriched hits of each selection. Fluorescence values are given relative to that of the parent $\beta^3$_tRNA$^{Pyl}$ with the same reporter.

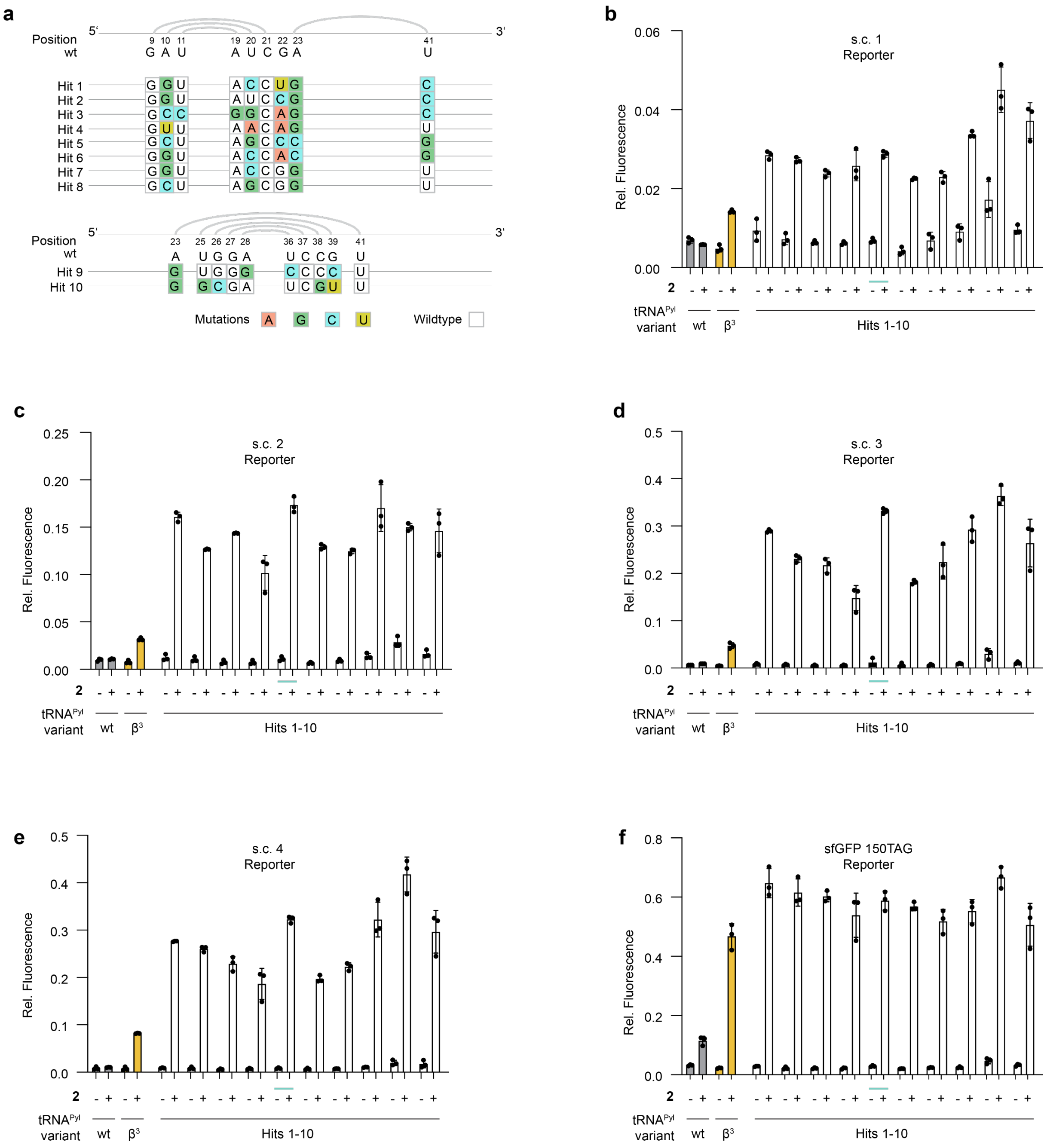


**Supplementary Figure 46. Testing ncM dependence of tRNAs derived from sequence context tRNA selections for β³-amino acid** 2**.**

**a**, tRNA hits selected on the basis of the primary screen in **Supplementary Fig. 45**. Only the targeted positions are shown. Mutated positions are coloured. The secondary structure topological pairing of the targeted residues is shown by connecting grey lines.

**b-f**, GFP fluorescence values of tRNA sequences resulting from parallel selections with library $\beta^3$_2 or library $\beta^3$_3. Hits were expressed with PylRS($\beta^3$-1) and tested with reporters: s.c.1 (**b**), s.c.2 (**c**), s.c.3 (**d**), s.c.4 (**e**) and sfGFP(150TAG) (**f**). Expression was conducted in the presence and absence of **2.** Fluorescence was normalised to expression of sfGFP(150TAG) with wt PylRS in the presence of **1** (2 mM). Hit 5, subsequently named $\beta^3$_v2_tRNA$^{Pyl}$ and used for subsequent experiments is underlined in blue.

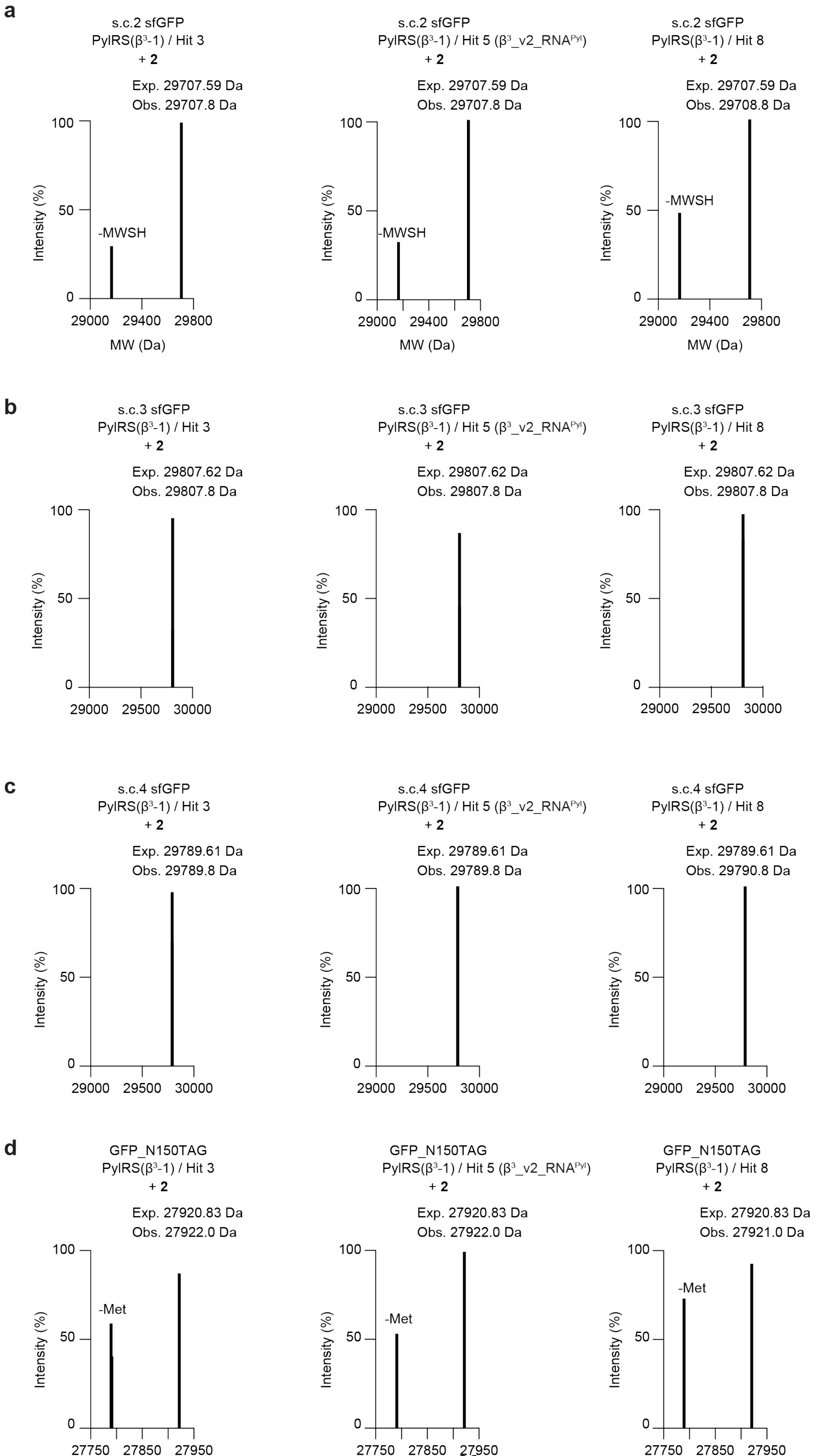
a
s.c.2 sfGFP
PylRS(β3-1) / Hit 3
+ 2
Exp. 29707.59 Da
Obs. 29707.8 Da
-MWSH
s.c.2 sfGFP
PylRS(β3-1) / Hit 5 (β3_v2_RNAPyl)
+ 2
Exp. 29707.59 Da
Obs. 29707.8 Da
-MWSH
s.c.2 sfGFP
PylRS(β3-1) / Hit 8
+ 2
Exp. 29707.59 Da
Obs. 29708.8 Da
-MWSH
Intensity (%)
MW (Da)
b
s.c.3 sfGFP
PylRS(β3-1) / Hit 3
+ 2
Exp. 29807.62 Da
Obs. 29807.8 Da
s.c.3 sfGFP
PylRS(β3-1) / Hit 5 (β3_v2_RNAPyl)
+ 2
Exp. 29807.62 Da
Obs. 29807.8 Da
s.c.3 sfGFP
PylRS(β3-1) / Hit 8
+ 2
Exp. 29807.62 Da
Obs. 29807.8 Da
c
s.c.4 sfGFP
PylRS(β3-1) / Hit 3
+ 2
Exp. 29789.61 Da
Obs. 29789.8 Da
s.c.4 sfGFP
PylRS(β3-1) / Hit 5 (β3_v2_RNAPyl)
+ 2
Exp. 29789.61 Da
Obs. 29789.8 Da
s.c.4 sfGFP
PylRS(β3-1) / Hit 8
+ 2
Exp. 29789.61 Da
Obs. 29790.8 Da
d
GFP_N150TAG
PylRS(β3-1) / Hit 3
+ 2
Exp. 27920.83 Da
Obs. 27922.0 Da
-Met
GFP_N150TAG
PylRS(β3-1) / Hit 5 (β3_v2_RNAPyl)
+ 2
Exp. 27920.83 Da
Obs. 27922.0 Da
-Met
GFP_N150TAG
PylRS(β3-1) / Hit 8
+ 2
Exp. 27920.83 Da
Obs. 27921.0 Da
-Met

**Supplementary Figure 47. Mass spectrometry characterisation of tRNA hits.**

Intact ESI-MS of sfGFP produced from s.c.2 (**a**), s.c.3 (**b**), and s.c.4 (**c**) and sfGFP(150TAG)$His_6$ (**d**) reporter in cells transformed with a plasmid encoding for PylRS($\beta^3$-1) and $tRNA^{Pyl}$ variants obtained from the second stage of evolution with substrate **2**. The expected monoisotopic masses correspond to the found masses in all cases confirming the quantitative incorporation of ncM **2**.

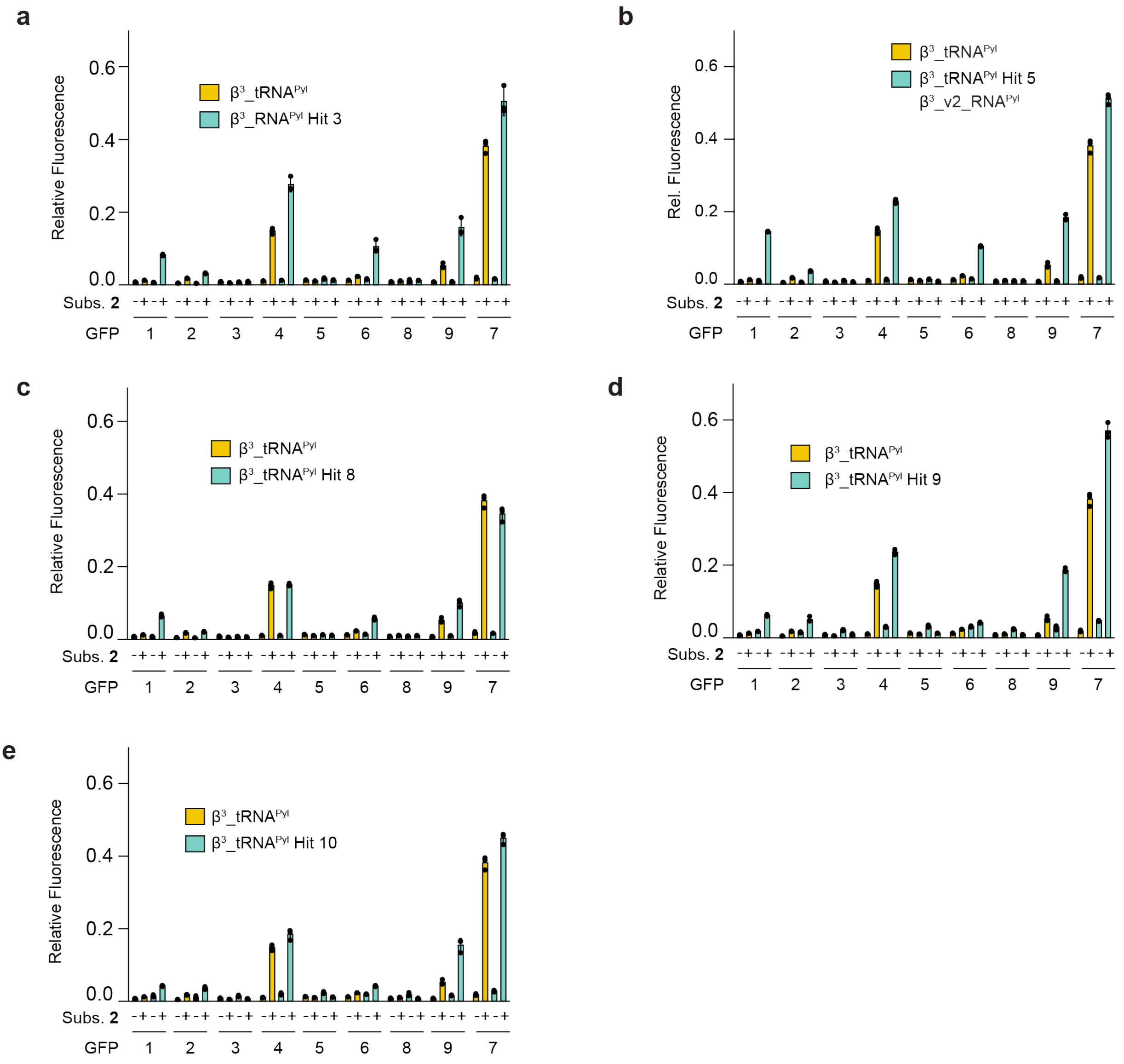


**Supplementary Figure 48. Testing tRNAs derived from sequence context tRNA selections for β$^3$-amino acid** 2 **with collection of 9 GFP reporters.**

**a-e** GFP fluorescence from cells transformed with sfGFP reporters containing a TAG codon at 9 different positions throughout the protein. Expression was performed with cells bearing a plasmid encoding for PylRS(β$^3$-1) and a tRNA hit from the second iteration of evolution with substrate **2**: Hit 3 (**a**), Hit 5/ β$^3$_v2_tRNA$^{Pyl}$ (**b**), Hit 8 (**c**), Hit 9 (**d**), Hit 10 (**e**), in the presence and absence of ncM **2**. An expression was conducted also with the parent β$^3$_tRNA$^{Pyl}$ and the data for this is shown in all panels. The positions of the suppressed codon in the sfGFP reporters 1-9 are: 39, 44, 50, 98, 109, 116, 150, 154, and 187.

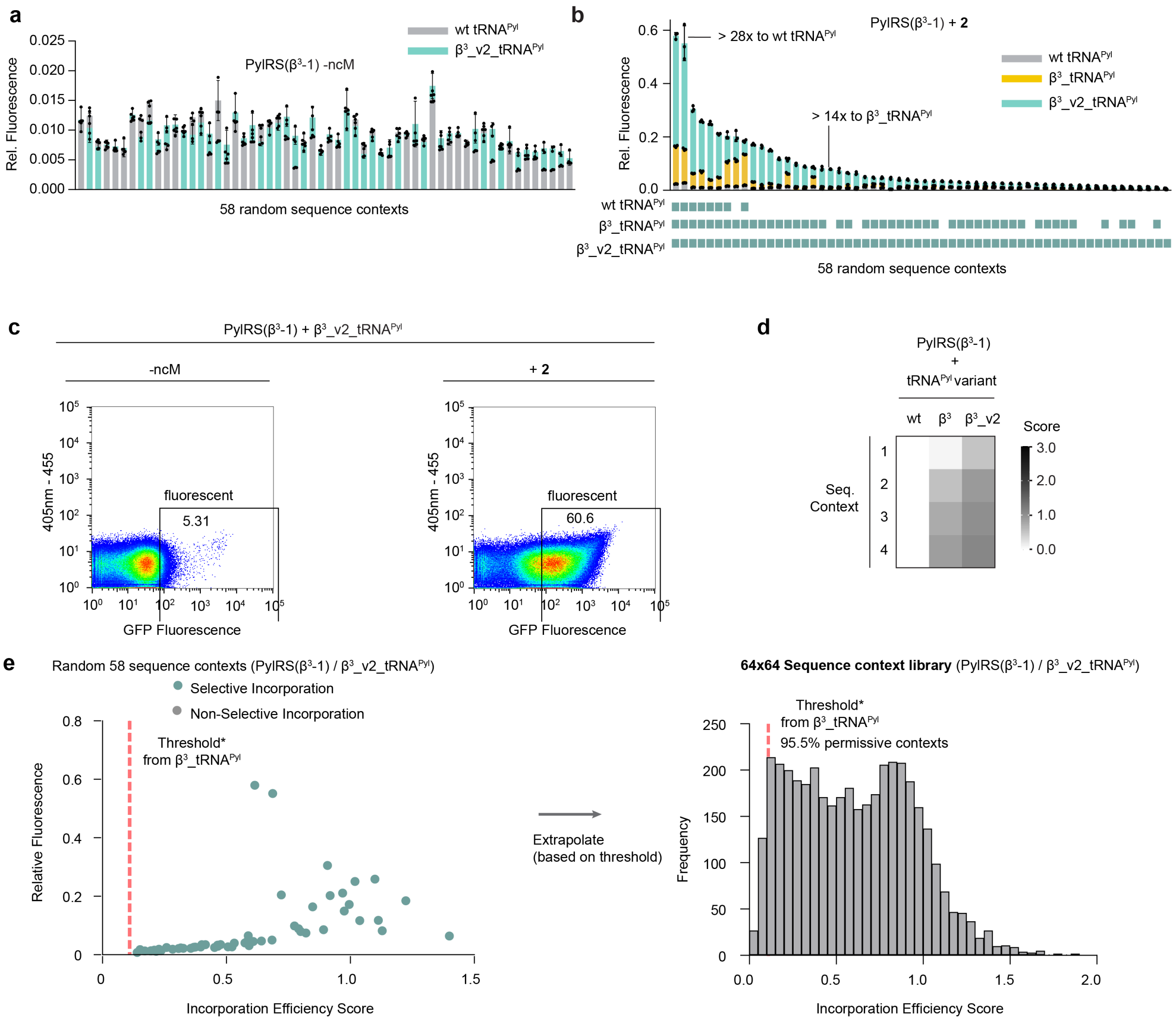


**Supplementary Figure 49. FACS analysis of sequence context libraries for second generation $\beta^3$_v2_tRNA$^{Pyl}$.**

**a,b**, GFP fluorescence levels of 58 random sequence contexts transformed into cells harbouring PylRS($\beta^3$-1) and either wt tRNA$^{Pyl}$, $\beta^3$_tRNA$^{Pyl}$ or $\beta^3$_v2_tRNA$^{Pyl}$ in the absence (**a**) or presence (**b**) of **2**. Fluorescence was normalised to expression of sfGFP(150TAG) with wt PylRS in the presence of **1** (2 mM).

**c**, Fluorescence distribution measured by FACS of N-terminal sequence context library expressed with PylRS($\beta^3$-1) and $\beta^3$_v2_tRNA$^{Pyl}$ in the absence (left) or presence of **2** (right).

**d**, Incorporation efficiency score values for the four selected sequence contexts 1, 2, 3, and 4 with PylRS($\beta^3$-1) paired with either wt tRNA$^{Pyl}$, $\beta^3$_tRNA$^{Pyl}$ or $\beta^3$_v2_tRNA$^{Pyl}$. Values were extracted from data shown in **Fig. 5h**.

**e**, Fluorescence values of 58 random sequence context reporters expressed with the PylRS($\beta^3$-1)/$\beta^3$_v2_tRNA$^{Pyl}$ pair in the presence of **2** were plotted against the incorporation efficiency score derived by FACS/NGS analysis. Sequences with a relative incorporation score higher than the lowest experimentally active (>1.2 fold change upon addition of ncM) variant were considered to support selective incorporation of **2**. We set a threshold value for the incorporation efficiency score at the lowest context supporting selective incorporation, but above the highest non-selective incorporation at a sequence context (red dashed line). This threshold value (taken from $\beta^3$_tRNA$^{Pyl}$ experiments, 0.109) was used to extrapolate the number of permissive sequence context across the entire sequence context library as measured by FACS/NGS. The distributions of incorporation efficiency scores for both samples are shown as histograms. The threshold derived is indicated as a red dashed line.

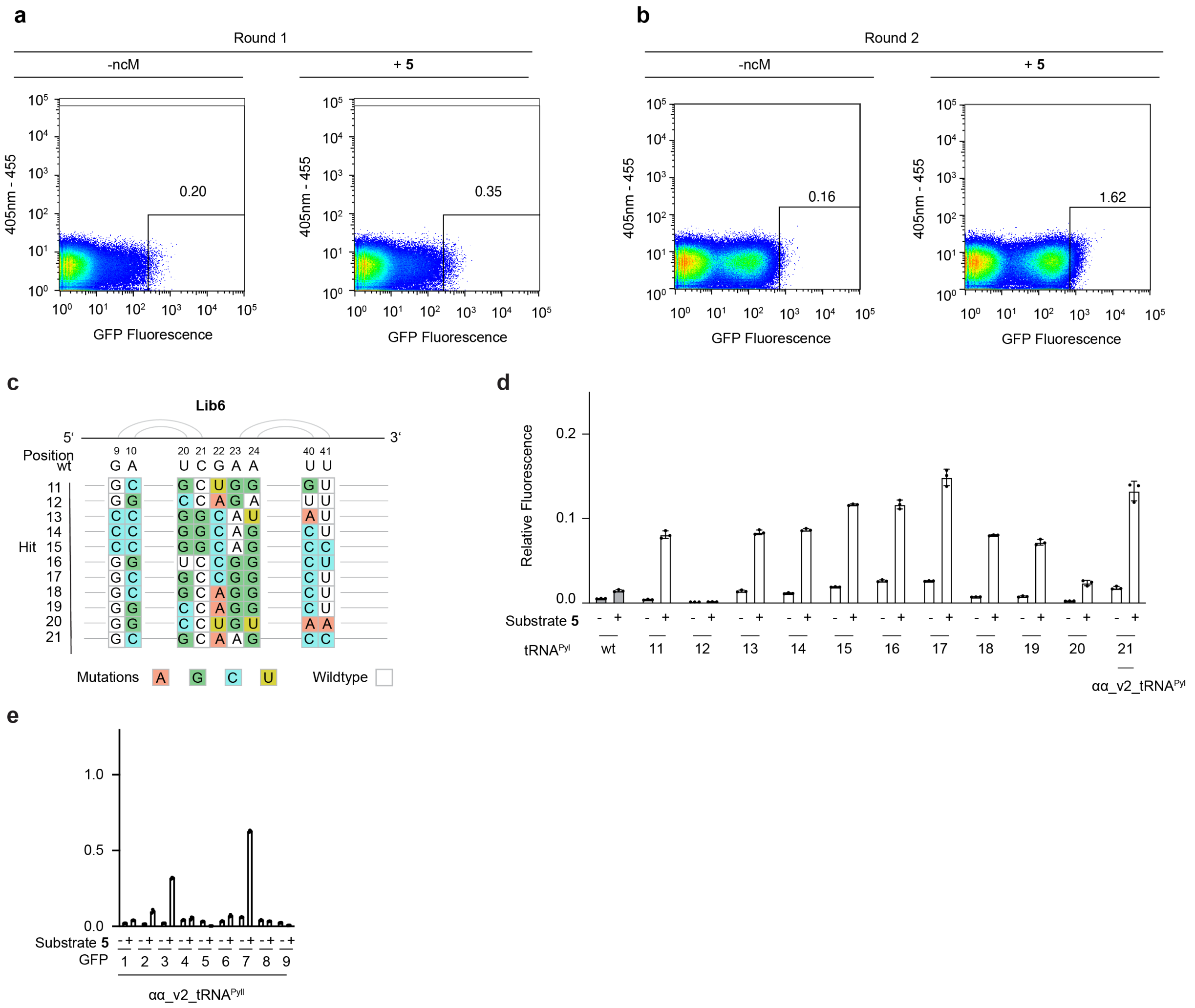


**Supplementary Figure 50. FACS-based tRNA selection for α,α-disubstituted amino acid on s.c.50.**

**a-b**, Two-round FACS of library 6 expressed in the presence and absence of **5**, selected on s.c.50, a less permissive sequence context than sfGFP(150TAG)-His$_6$. Scatter plots represent GFP-fluorescence vs. a second channel (excitation at 405 nm, emission at 455 nm).

**c**, Top 11 most enriched hits from library 6 evolution on s.c.50 for substrate **5**. Only the targeted positions are shown. Mutated positions are coloured. The secondary structure topological pairing of the targeted residues is shown by connecting grey lines.

**d**, GFP fluorescence of individually tested sequences resulting from evolutions with library 6. Cells were transformed with a s.c.50 reporter and a plasmid encoding for PylRS(αα-1) and different tRNA variants. Protein expression was induced in the presence and absence of substrate **5**. Fluorescence was normalised to expression of sfGFP(150TAG) with wt PylRS in the presence of **1** (2 mM).

**e**, GFP fluorescence of cells transformed with wildtype and evolved tRNAs expressing different sfGFP reporters. Cells were transformed with sfGFP reporters and a plasmid encoding for PylRS(αα-1) and αα _v2_tRNA$^{Pyl}$. The positions of the suppressed codon in the sfGFP reporters 1-9 are: 39, 44, 50, 98, 109, 116, 150, 154, and 187. Fluorescence was normalised to expression of sfGFP(150TAG) with wt PylRS in the presence of **1** (2 mM).

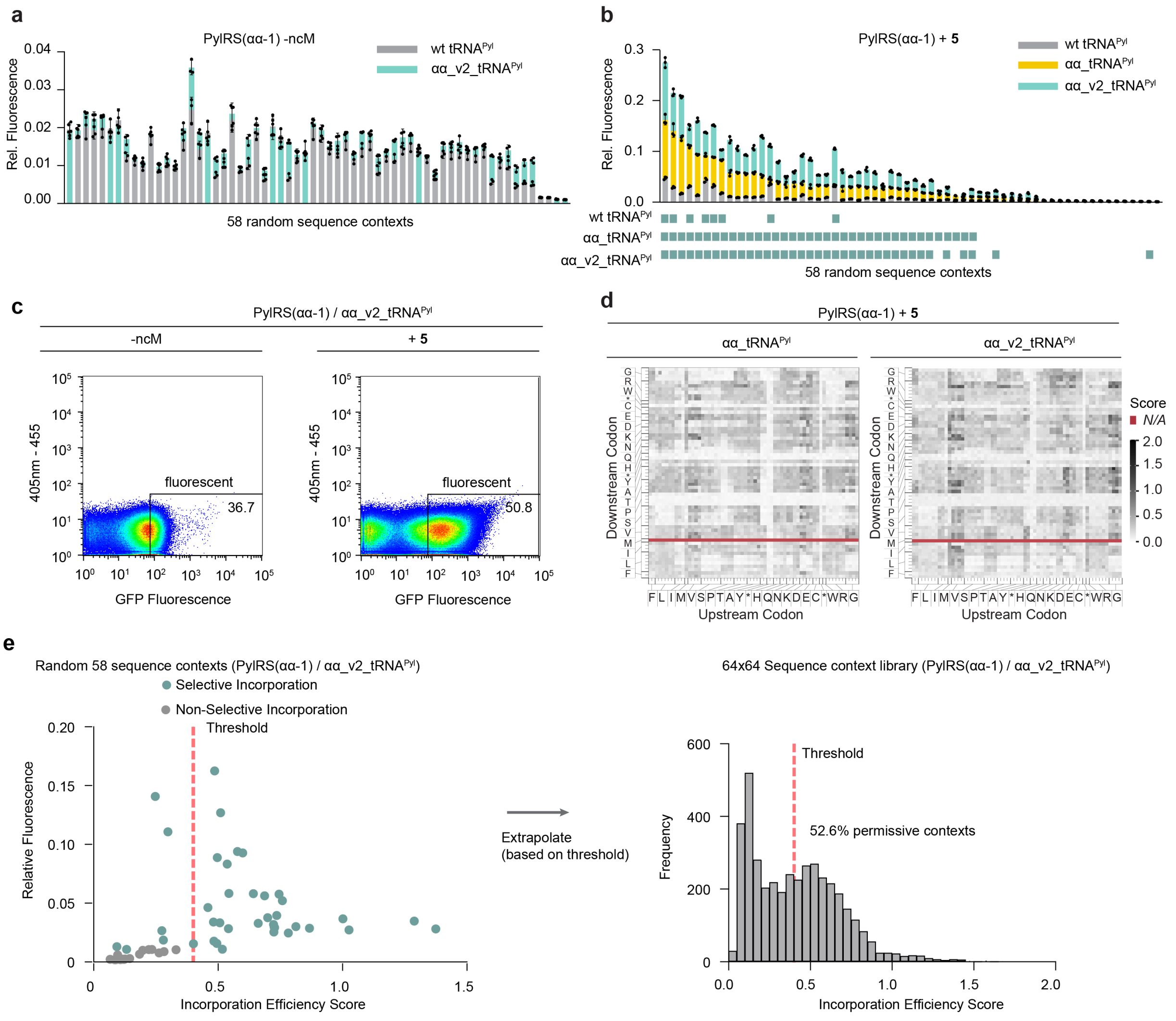


**Supplementary Figure 51. Characterisation of αα_v2_tRNA$^{Pyl}$.**

**a**, GFP fluorescence levels of 58 random sequence contexts transformed into cells harbouring PylRS(αα -1) and either wt tRNA$^{Pyl}$ (grey), αα _tRNA$^{Pyl}$ (yellow) or further evolved αα_v2_tRNA$^{Pyl}$ (blue) and expressed in the absence (**a**) or (**b**) presence of substrate **5**. Fluorescence was normalised to expression of sfGFP(150TAG) with wt PylRS in the presence of **1** (2 mM).

**c**, Fluorescence distribution measured by FACS of N-terminal sequence context library expressed with PylRS(αα-1) with αα_v2_tRNA$^{Pyl}$ in the absence or presence of **5**.

**d**, Incorporation efficiency heat maps for translation of substrate **5** mediated by PylRS(αα-1) and wt tRNA$^{Pyl}$ (left) or further evolved αα_v2_tRNA$^{Pyl}$ (right). Data for αα_tRNA$^{Pyl}$ is duplicated from **Fig. 4n**

**e**, Fluorescence values of 58 random sequence context reporters expressed with the PylRS(αα-1)/αα _v2_tRNA$^{Pyl}$ pair in the presence of **5** were plotted against the incorporation efficiency score derived by FACS/NGS analysis. Sequences with a relative incorporation score higher than the lowest experimentally active (>1.2 fold change upon addition of ncM) variant were considered to support selective incorporation of **5**. We set a threshold value for the incorporation efficiency score at the lowest context supporting selective incorporation, but above the highest non-selective incorporation at a sequence context (red dashed line). This threshold value (0.373) was used to extrapolate the number of permissive sequence context across the entire sequence context library as measured by FACS/NGS. The distributions of incorporation efficiency scores for both samples are shown as histograms. The threshold derived is indicated as a red dashed line.

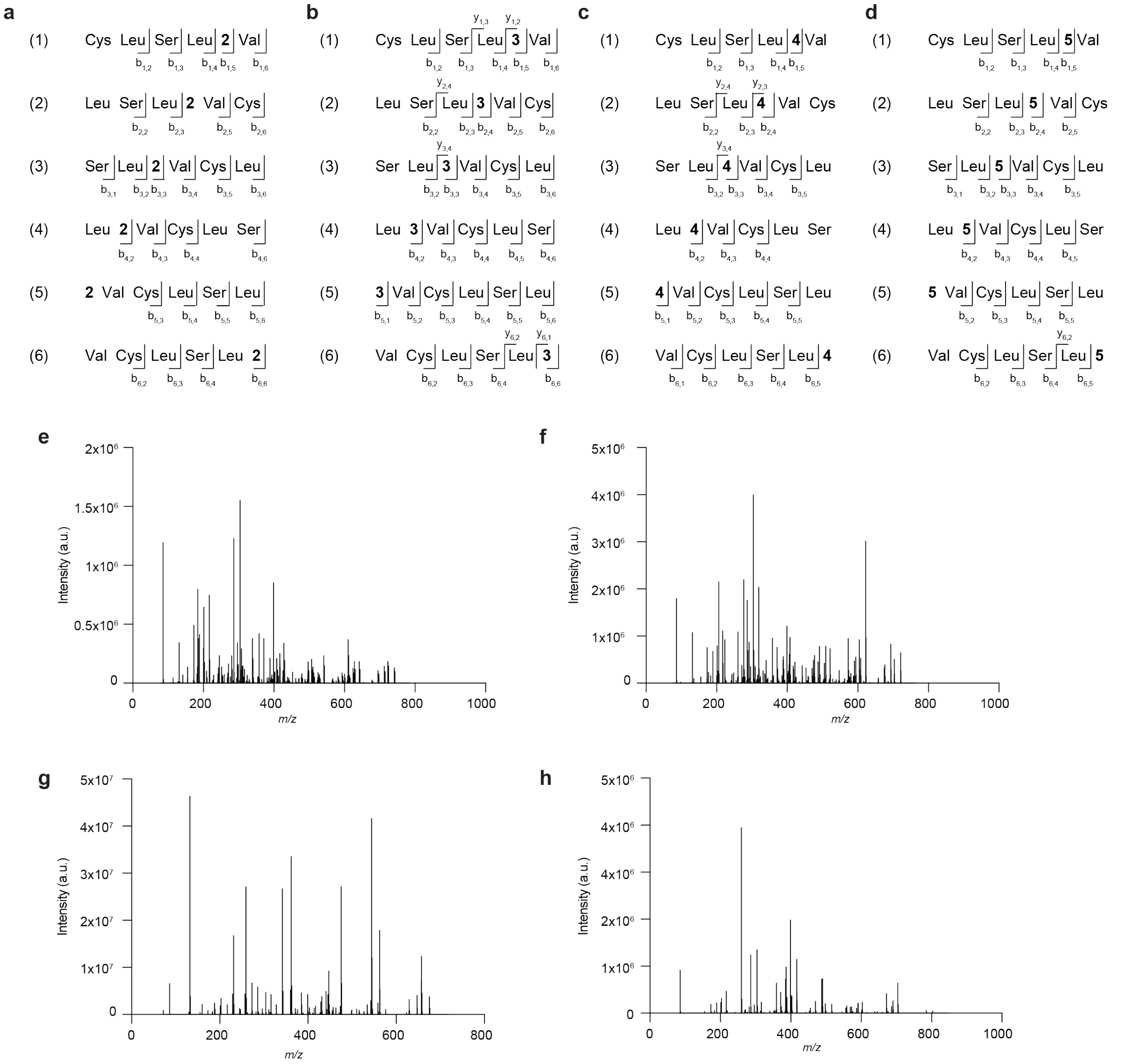


**Supplementary Figure 52. Raw MS2 spectra for cyclic peptides**

**a-h**, MS/MS fragmentation patterns and spectra of purified cyclic CLSL-X-V peptides where X corresponds to **2** (**a,e**, parent *m/z* = 741.2640), **3** (**b,f**, parent *m/z* = 721.3589), **4** (**c,g**, parent *m/z* = 675.3534), or **5** (**d,h,** parent *m/z* = 803.2657).

**a**

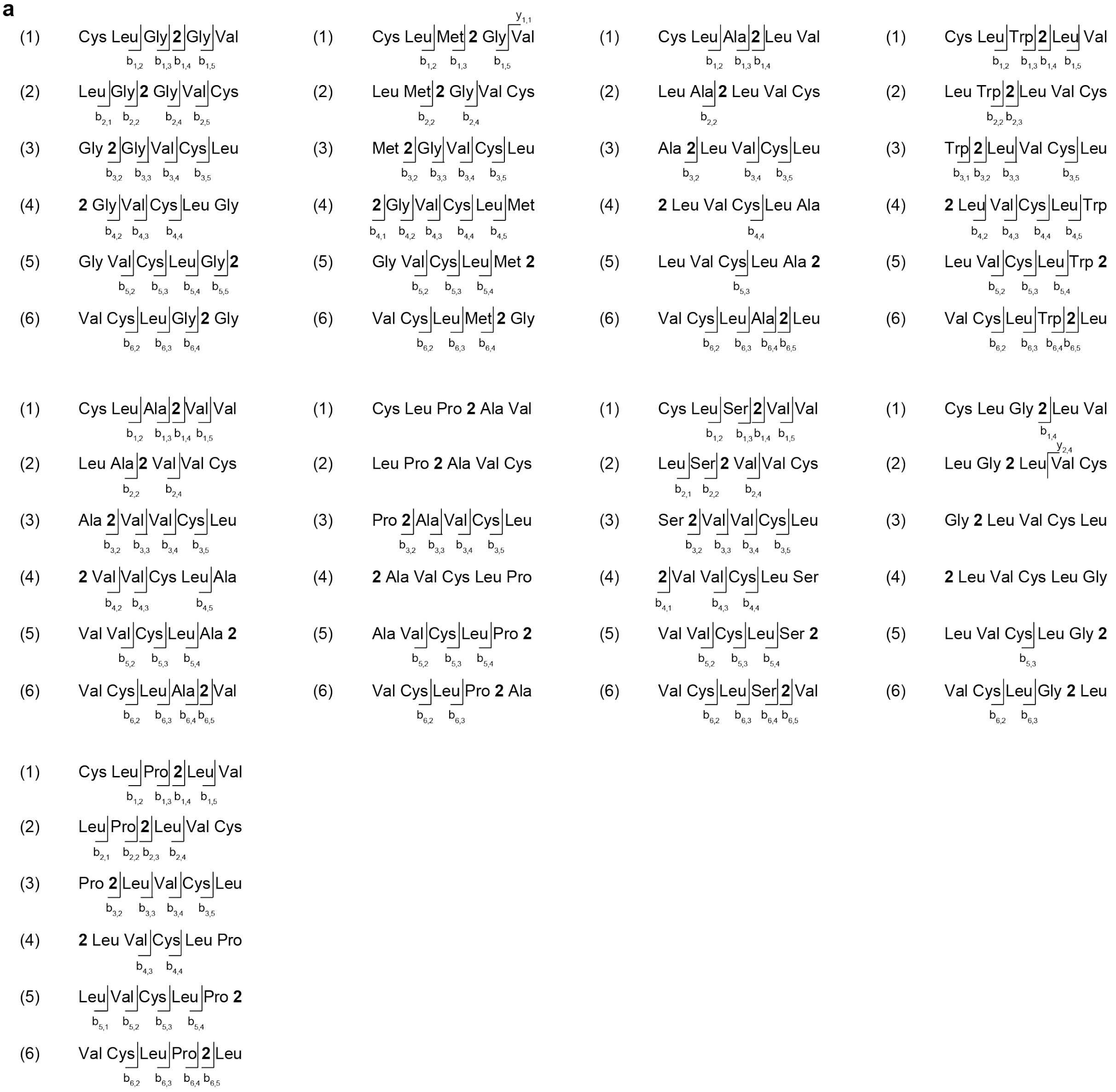


**Supplementary Figure 53. MS2 fragmentation for CLX$_1$-ncM-X$_2$V peptides**

**a**, MS/MS fragmentation patterns of purified cyclic CLX$_1$-ncM-X$_2$V where the ncM corresponds to **2**. A full list of tested peptides is included in Supplementary Data 1.

Supplementary Fig. 3d Top panel Cy5

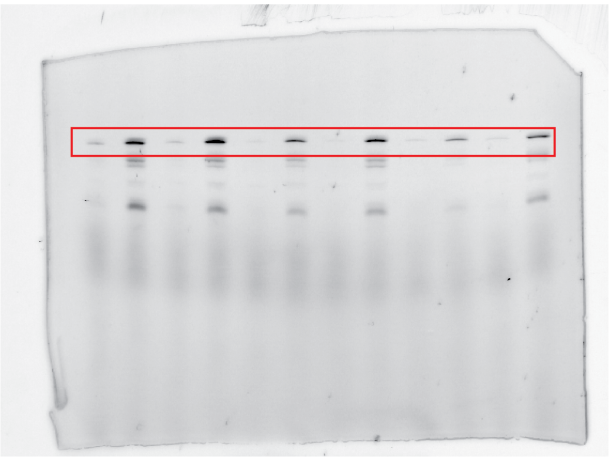

Supplementary Fig. 3d Top panel Cy3

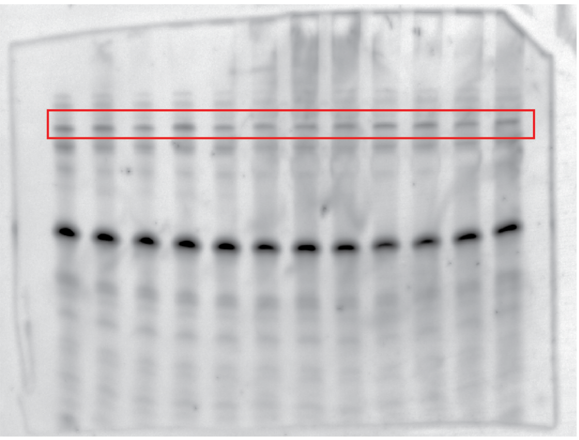

Supplementary Fig. 3d Second panel Cy5

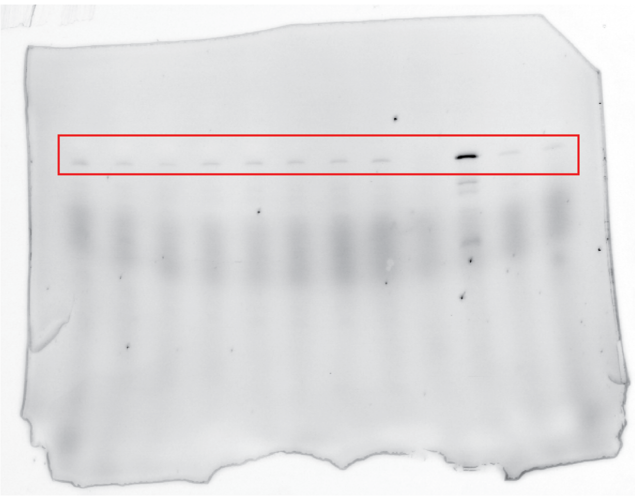

Supplementary Fig. 3d Second panel Cy3

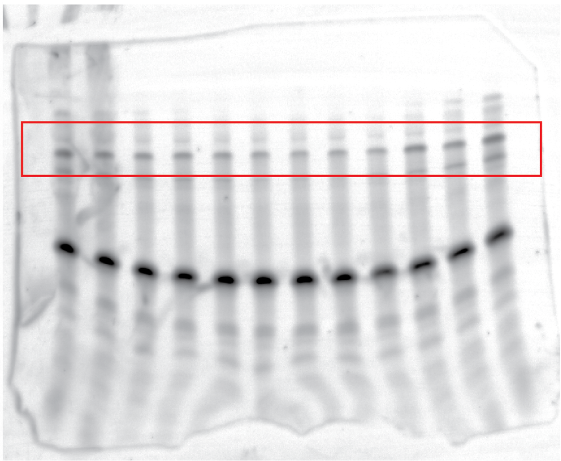

Supplementary Fig. 3d Third panel Cy5

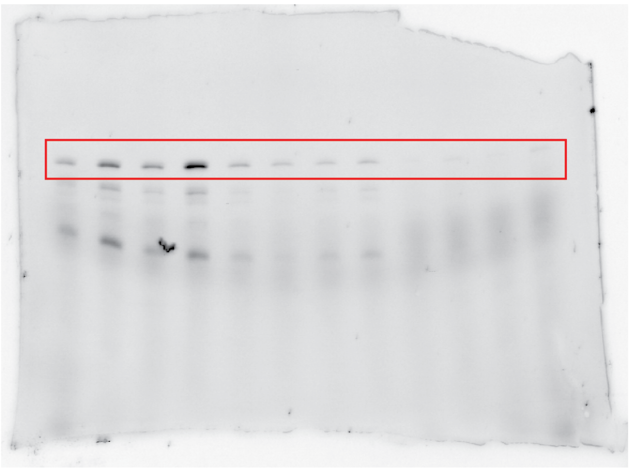

Supplementary Fig. 3d Third panel Cy3

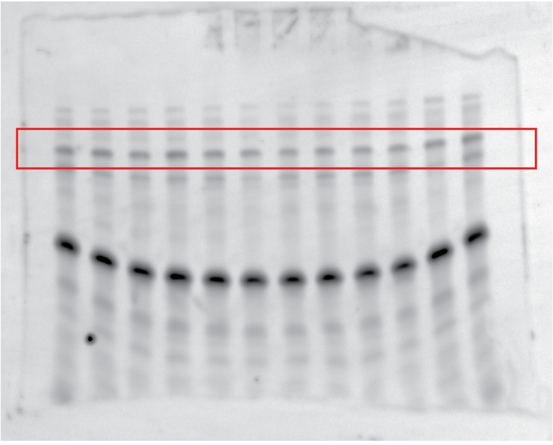

Supplementary Fig. 3d last panel Cy5

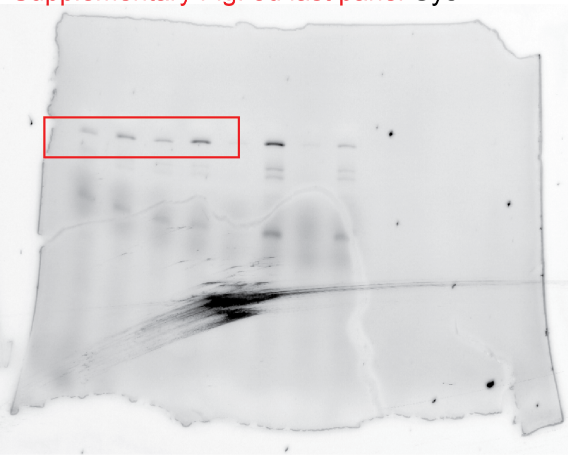

Supplementary Fig. 3d last panel Cy3

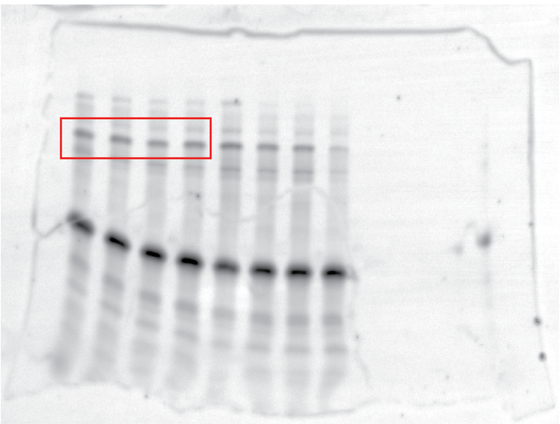

Supplementary Fig. 4d top panel Cy5

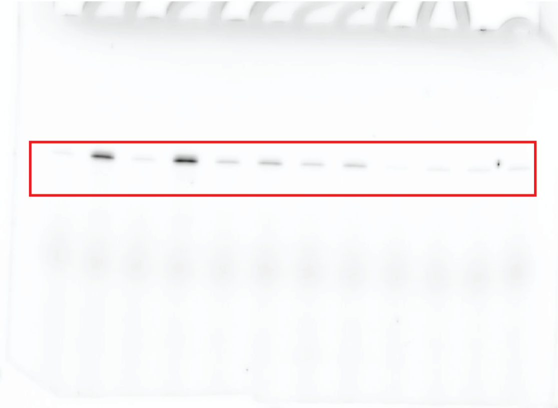

Supplementary Fig. 4d top panel Cy3

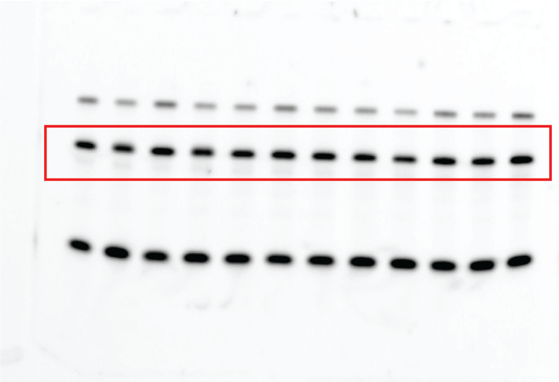

Supplementary Fig. 4d middle panel Cy5

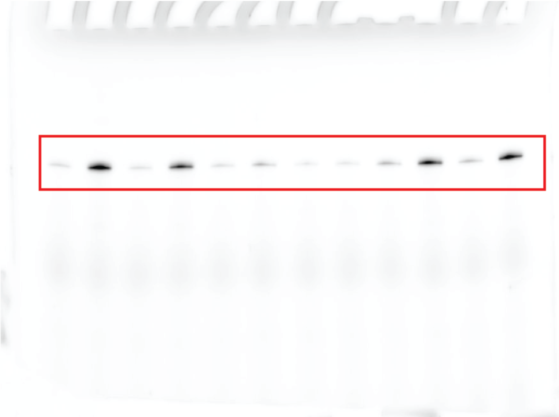

Supplementary Fig. 4d middle panel Cy3

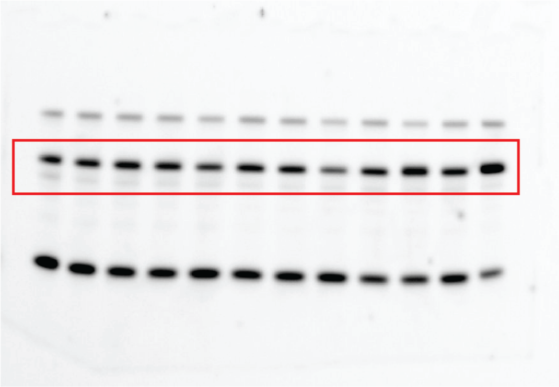

Supplementary Fig. 4d bottom panel Cy5

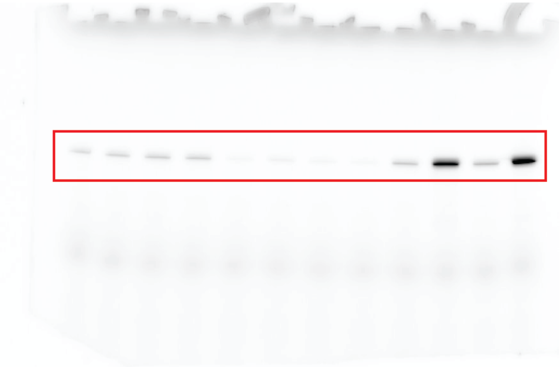

Supplementary Fig. 4d bottom panel Cy3

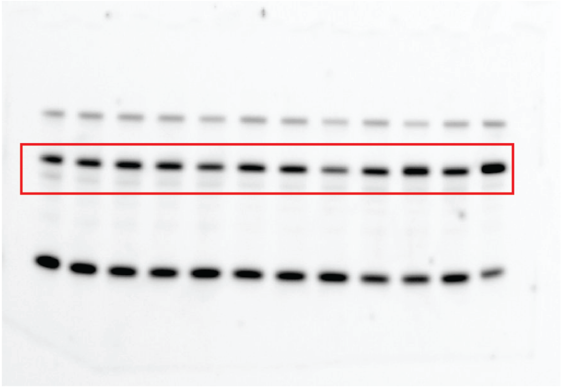

Fig.1f and Supplementary Fig.5d top panel, Cy5

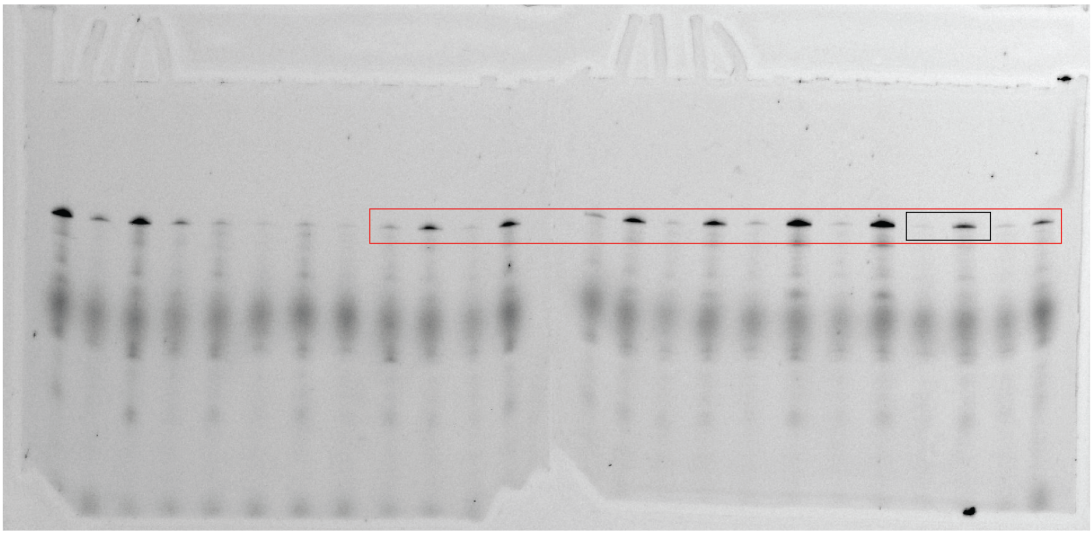

Fig.1f and Supplementary Fig.5d top panel, Cy3

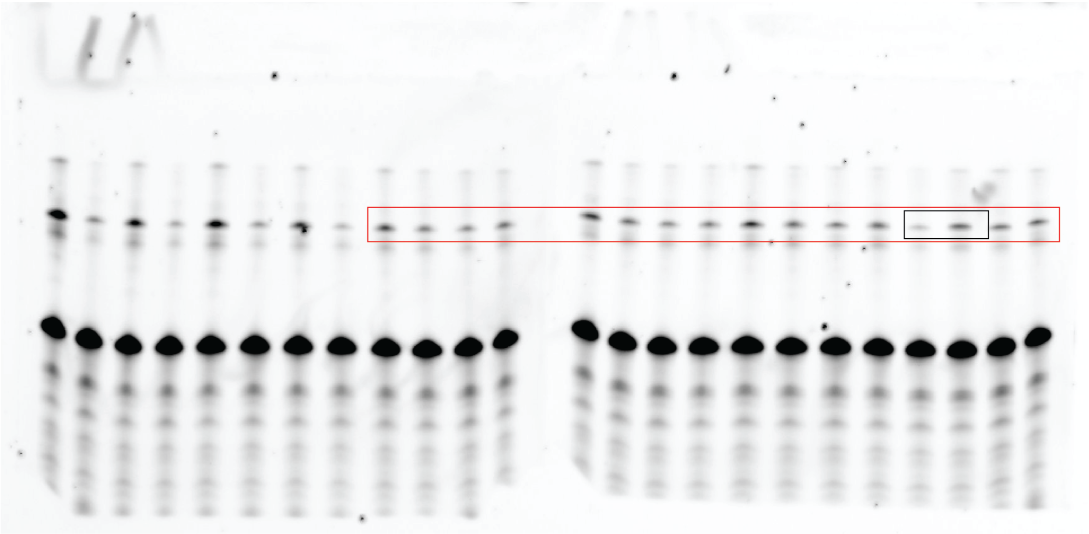

Supplementary Fig.5d middle panel, Cy5

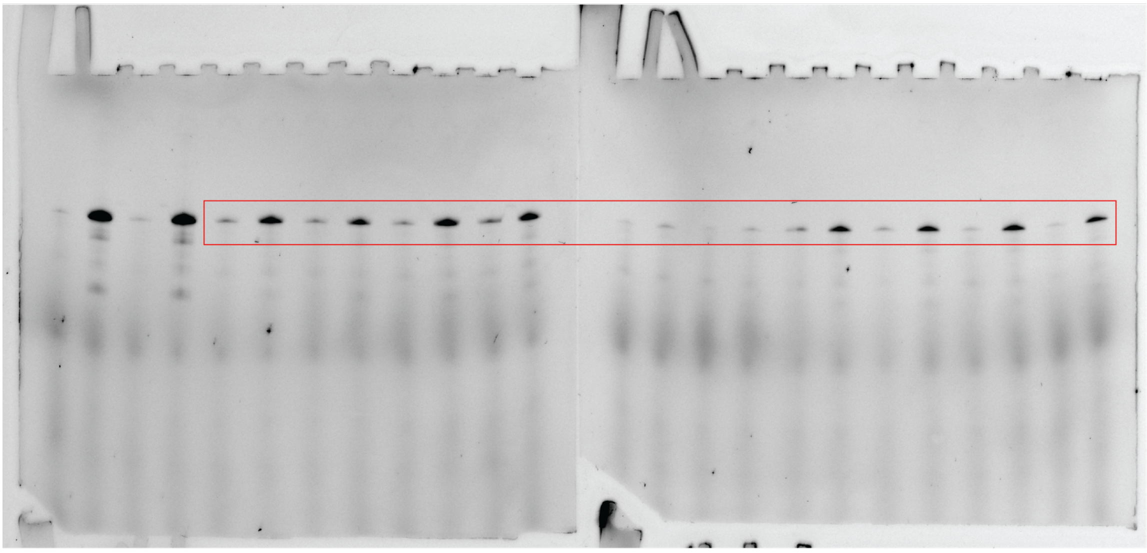

Supplementary Fig.5d middle panel, Cy3

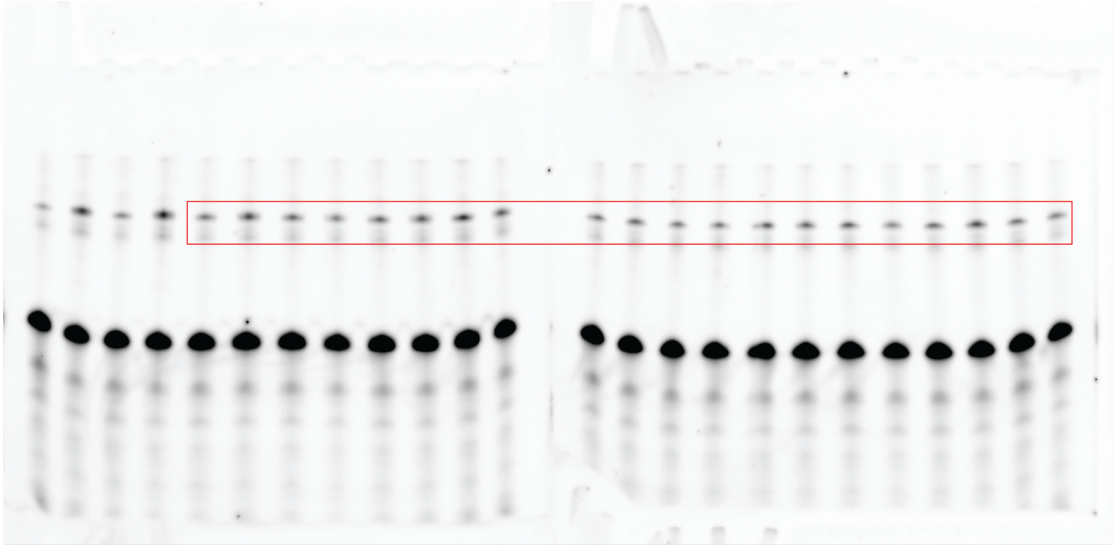

Supplementary Fig.5d bottom panel, Cy5

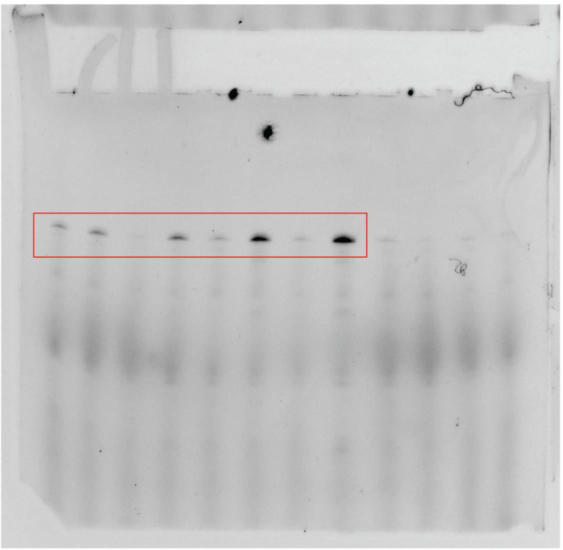

Supplementary Fig.5d bottom panel,, Cy3

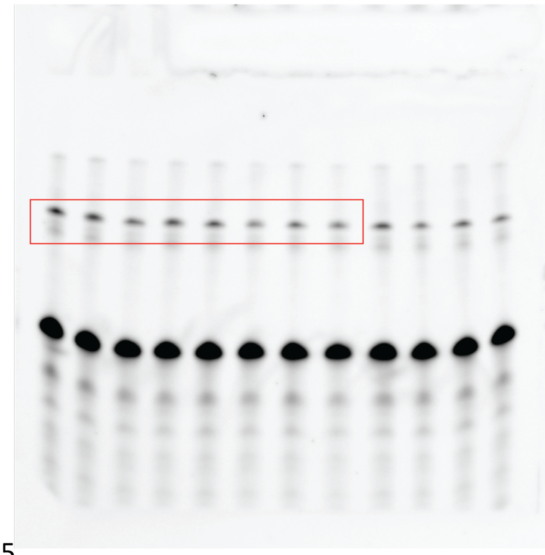

Supplementary Fig.6d top panel, Cy5

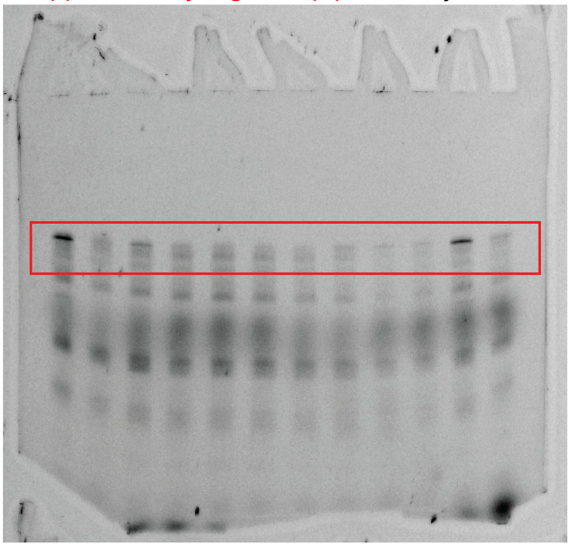

Supplementary Fig.6d top panel Cy3

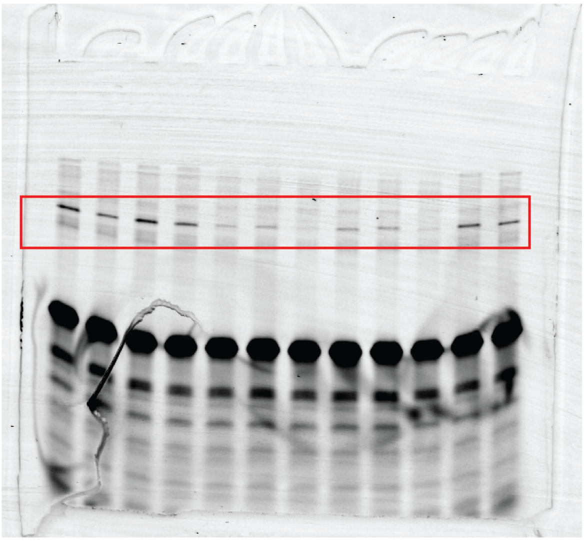

Supplementary Fig.6d second panel Cy5

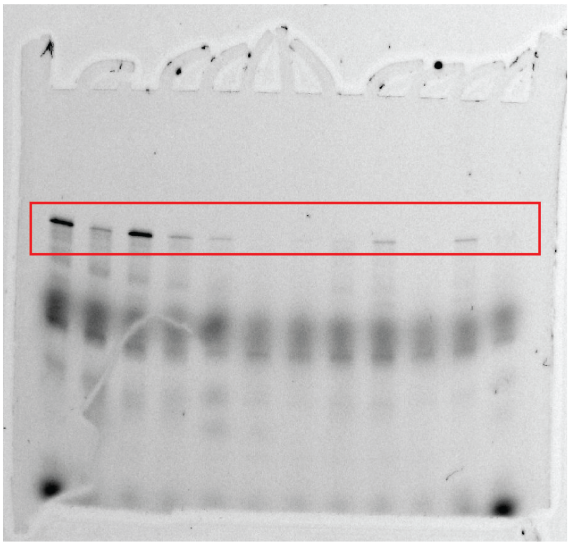

Supplementary Fig.6d second panel Cy3

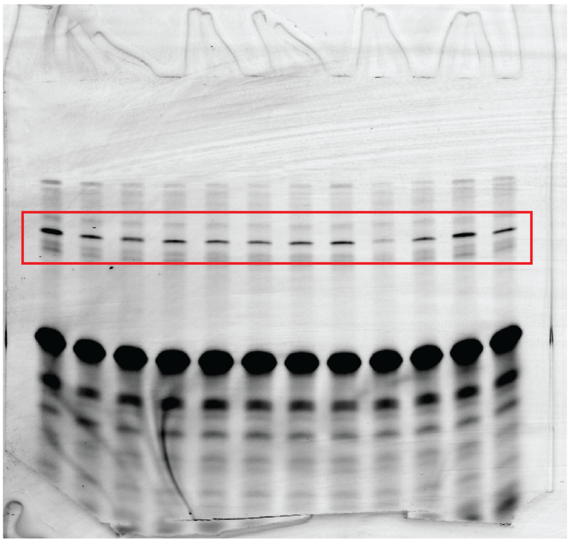

Supplementary Fig.6d third panel Cy5

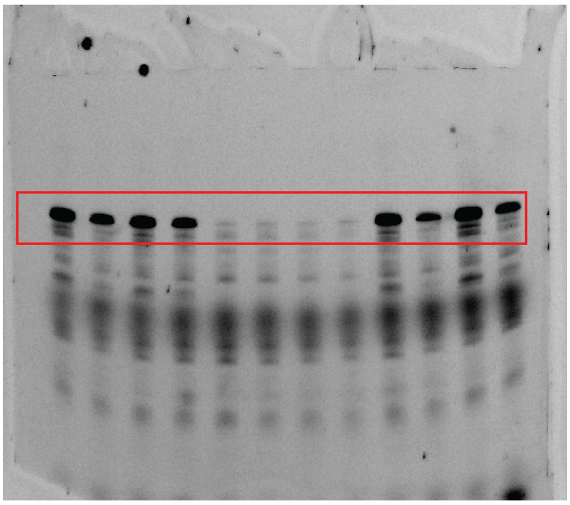

Supplementary Fig.6d third panel Cy3

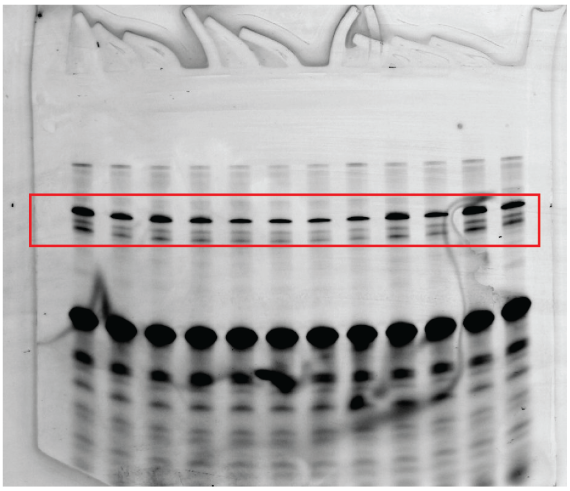

Fig. 1h, Supplementary Fig.6d bottom panel Cy5

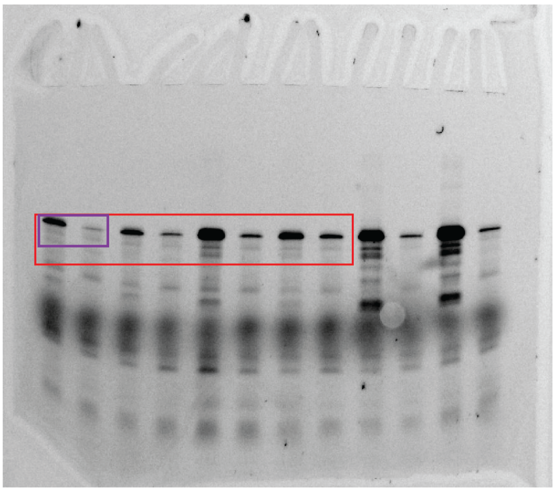

Fig. 1h, Supplementary Fig.6d bottom panell Cy3

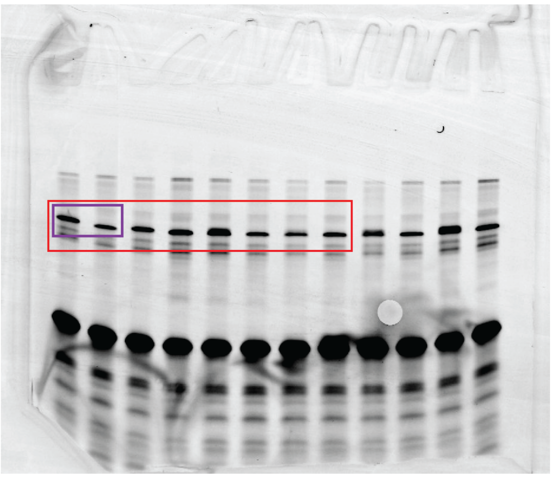

Fig. 1j, Supplementary Fig.7d top left panel Cy5

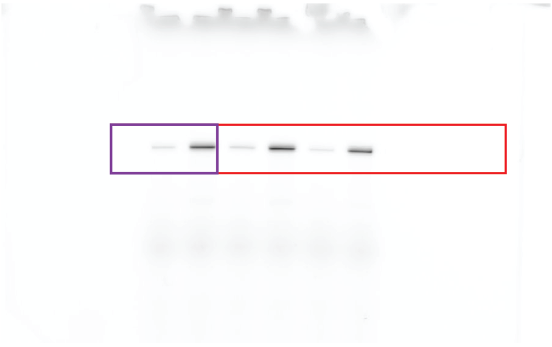

Fig. 1j, Supplementary Fig.7d top left panel Cy3

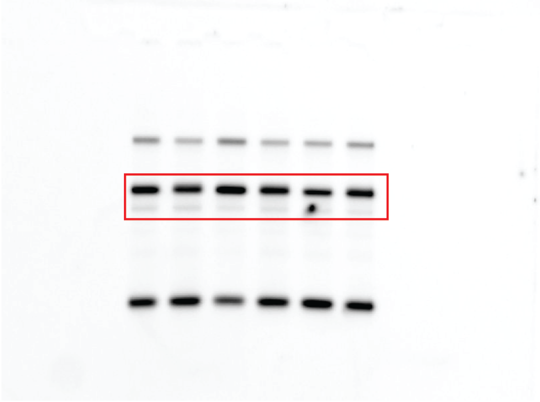

Supplementary Fig.7d top right panel Cy5

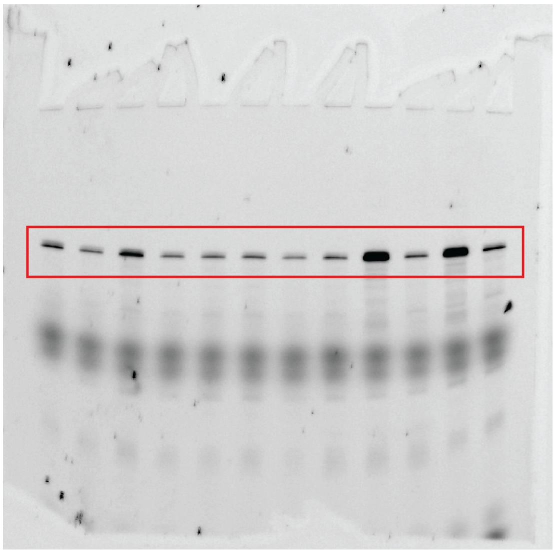

Supplementary Fig.7d top right panel Cy3

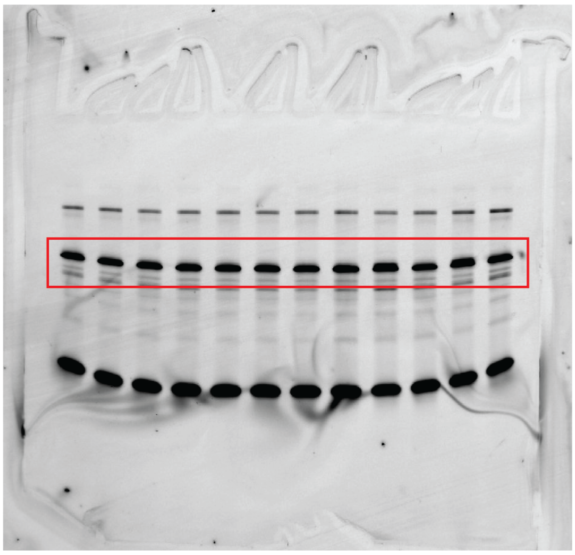

Supplementary Fig.7d bottom panel Cy5

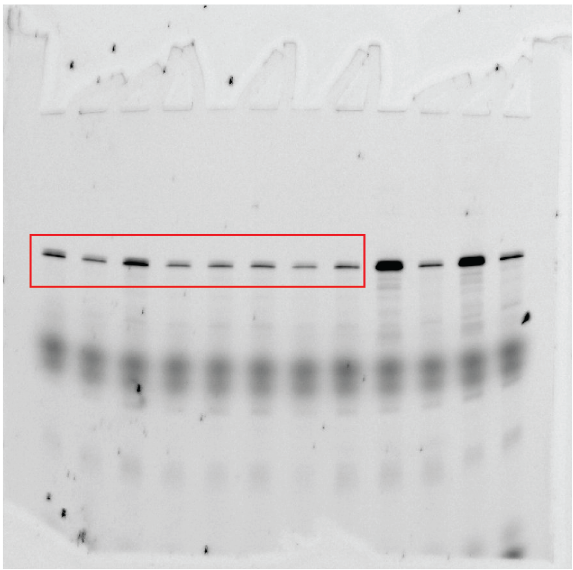

Supplementary Fig.7d bottom panel Cy3

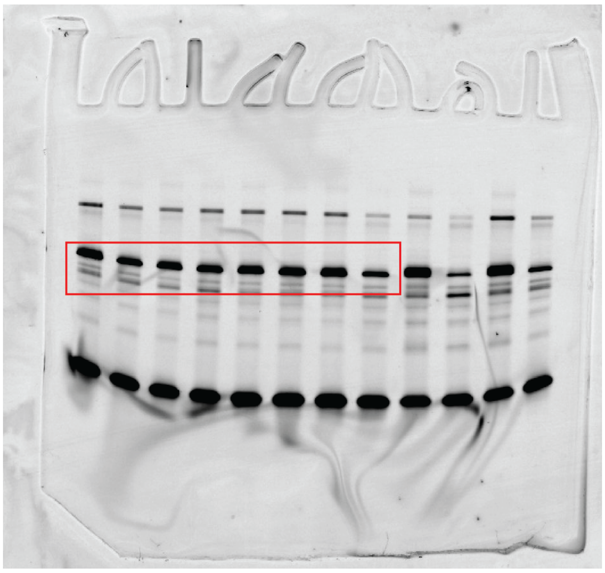

Supplementary Fig.9d, Cy5

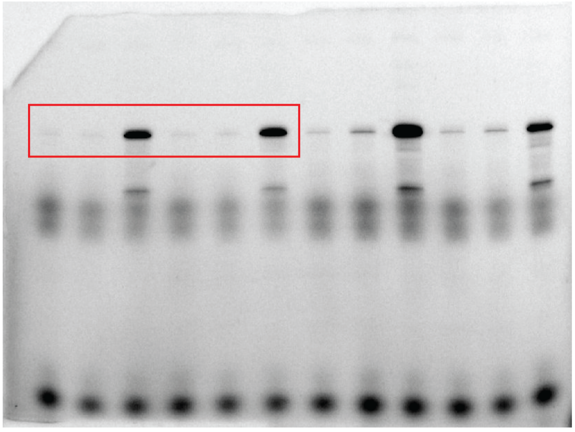

Supplementary Fig.9d, Cy3

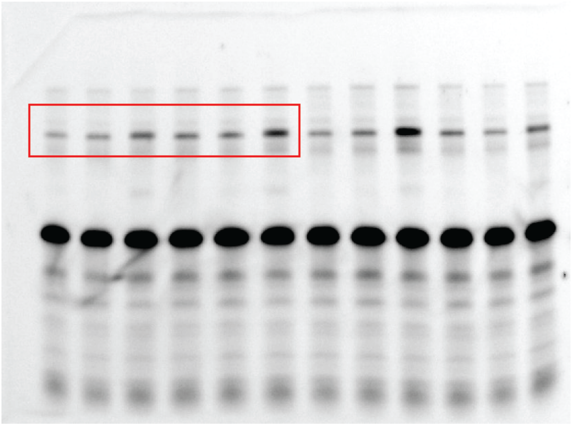

Supplementary Fig.10d Cy5

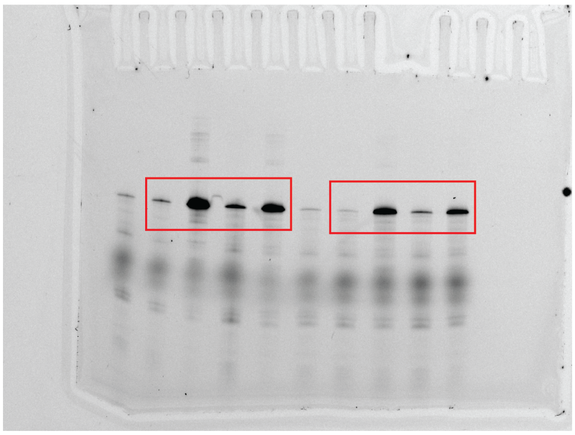

Supplementary Fig.10d Cy3

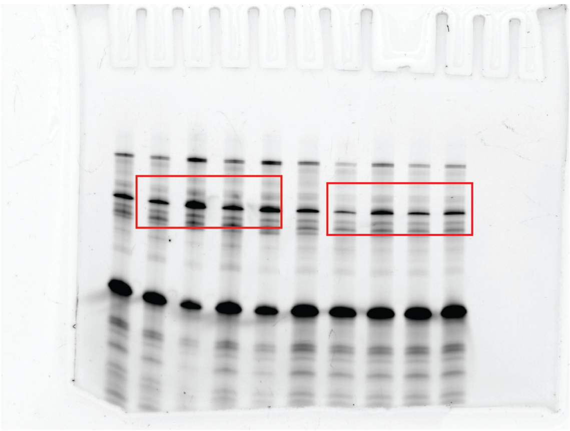

Supplementary Fig.11h, Cy5

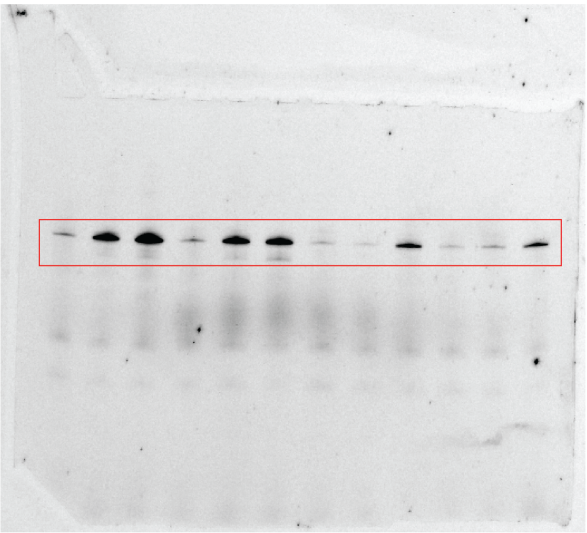

Supplementary Fig.11h, Cy3

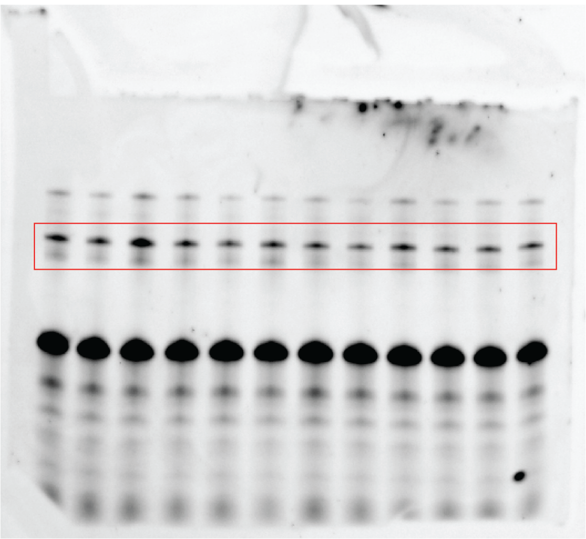

Supplementary Fig.12h, Cy5

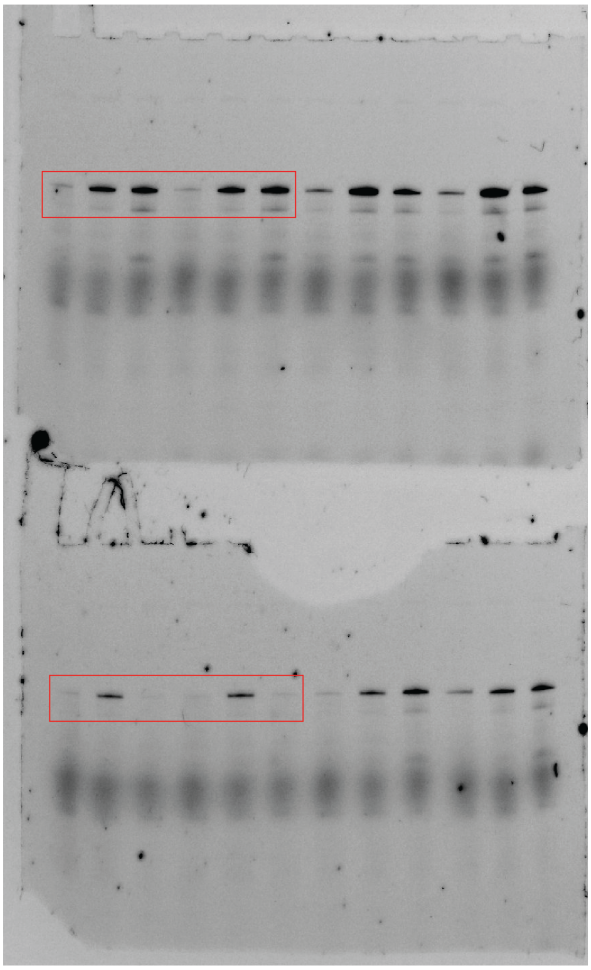

Supplementary Fig.12h, Cy3

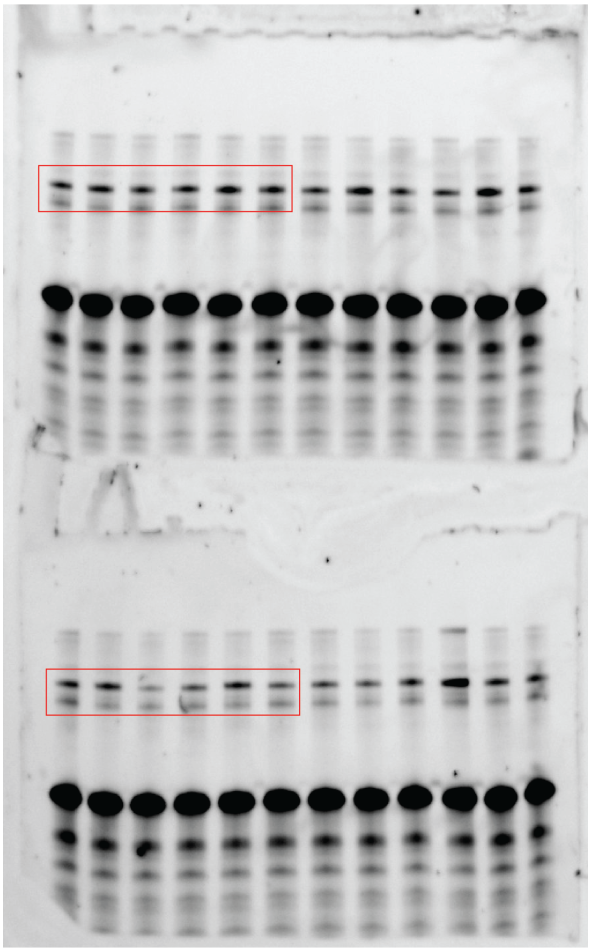

Supplementary Fig.17a, Cy5

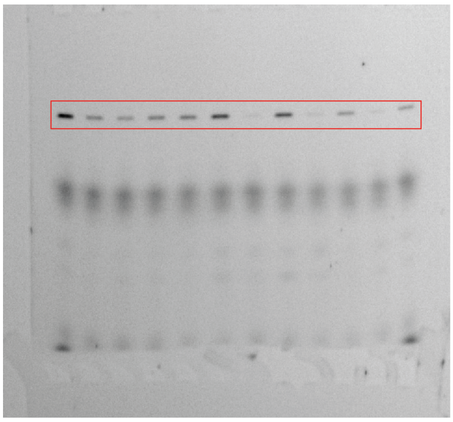

Supplementary Fig.17a, Cy3

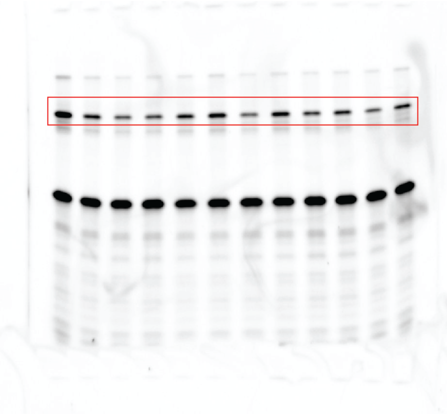

Supplementary Fig.18a, Cy5

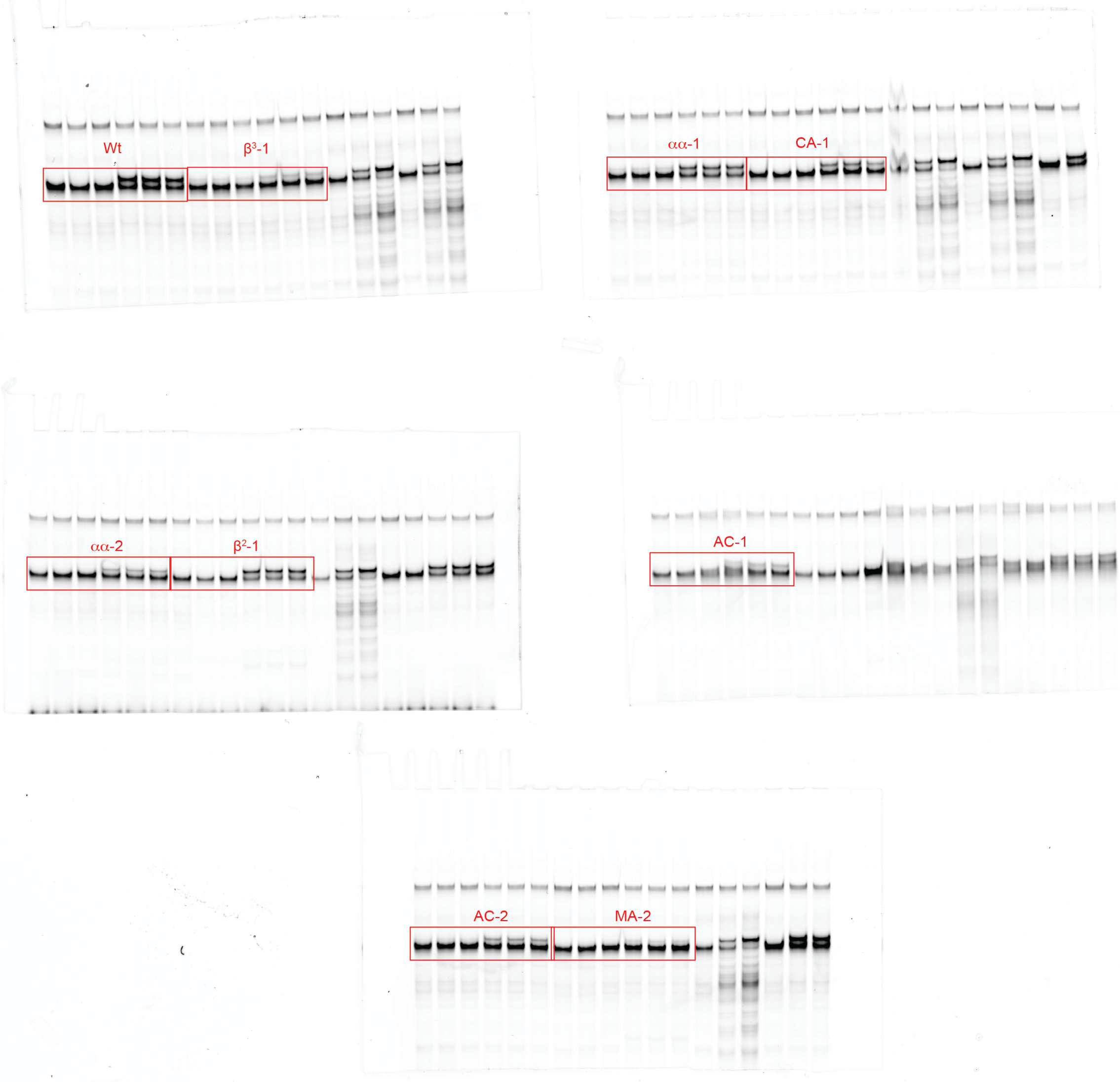

Supplementary Fig.19b Cy5

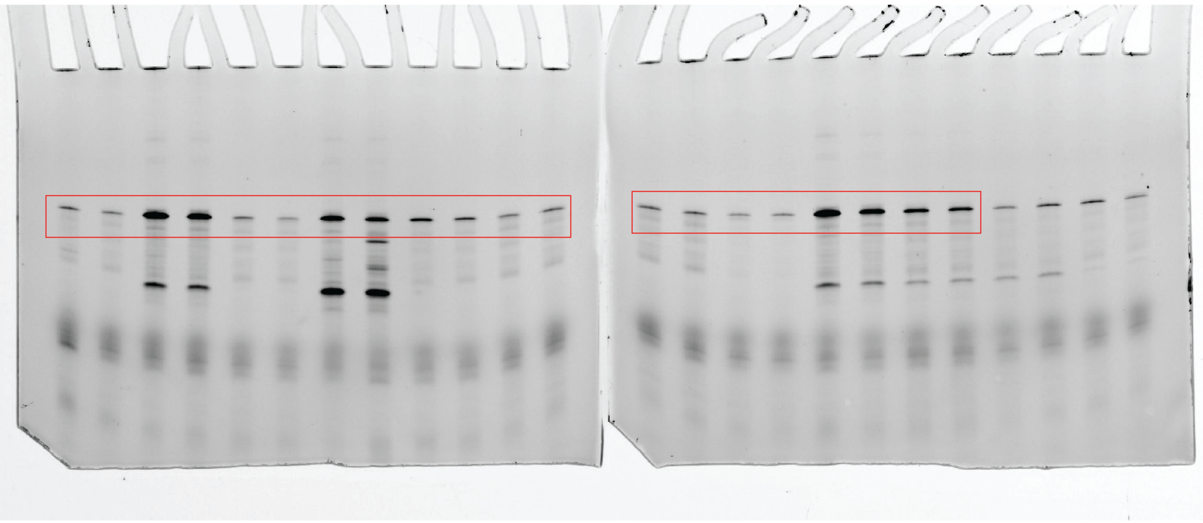

Supplementary Fig.19b Cy3

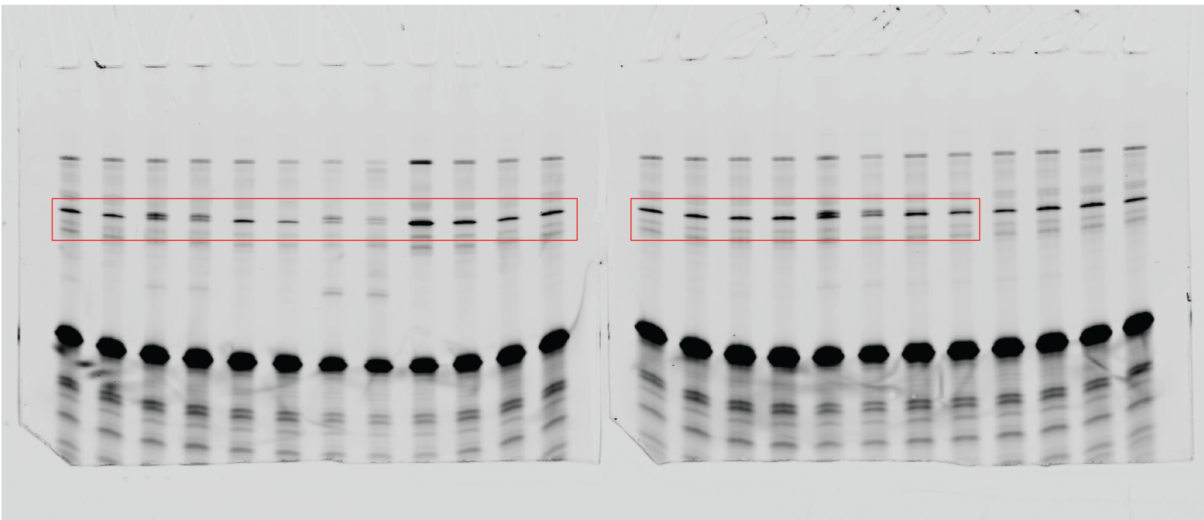

Supplementary Fig.20b Cy5

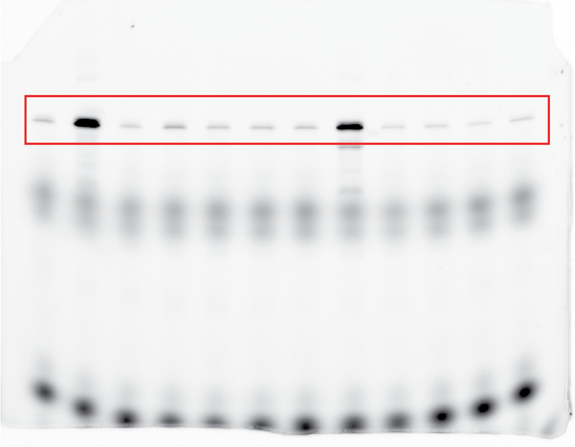

Supplementary Fig.20b Cy3

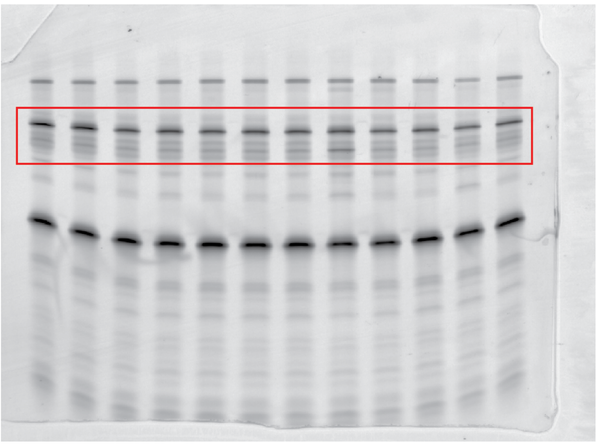

Supplementary Fig.20b Cy5

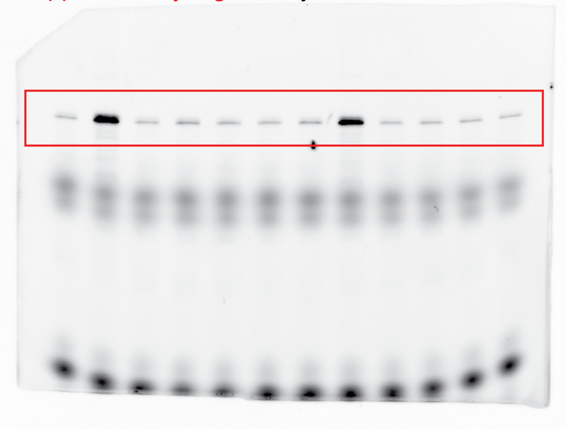

Supplementary Fig.20b Cy3

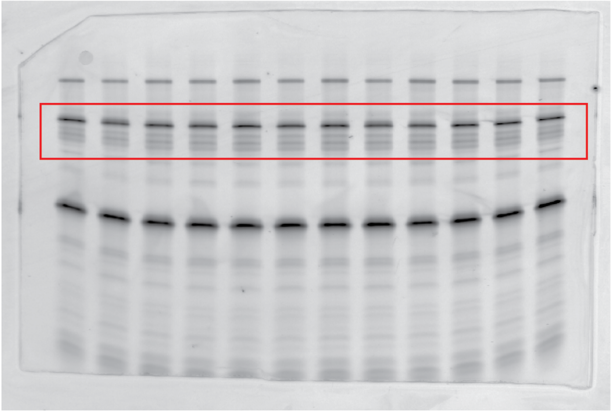

**Supplementary Figure 54. Full scans of gels run in this work.** The Figure in which the gel is shown is indicated at the top of each gel and the segment that was used in that figure is denoted by a box.
